# Bioinformatics analysis of differentially expressed genes in non alcoholic fatty liver disease using next generation sequencing data

**DOI:** 10.1101/2021.12.16.472893

**Authors:** Basavaraj Vastrad, Chanabasayya Vastrad

**Author notes:** Chanabasayya Vastrad Chanabasava Nilaya, Bharthinagar, Dharwad 580001, Karanataka, India.

## Abstract

Non alcoholic fatty liver disease (NAFLD) is the most common metabolic disease in humans, affecting the majority of individuals. In the current investigation, we aim to identify potential key genes linked with NAFLD through bioinformatics analyses of next generation sequencing (NGS) dataset. NGS dataset of GSE135251 from the Gene Expression Omnibus (GEO) database were retrieved. Differentially expressed genes (DEGs) were obtained by DESeq2 package. g:Profiler database was further used to identify the potential gene ontology (GO) and REACTOME pathways. Protein-protein interaction (PPI) network was constructed using the Hippie interactome database. miRNet and NetworkAnalyst databases were used to establish a miRNA-hub gene regulatory network and TF-hub gene regulatory network for the hub genes. Hub genes were verified based on receiver operating characteristic curve (ROC) analysis. Totally, 951 DEGs were identified including 476 up regulated genes and 475 down regulated genes screened in NAFLD and normal control. GO showed that DEGs were significantly enhanced for signaling and regulation of biological quality. REACTOME pathway analysis revealed that DEGs were enriched in signaling by interleukins and extracellular matrix organization. ESR2, JUN, PTN, PTGER3, CEBPB, IKBKG, HSPA8, SFN, CDKN1A and E2F1 were indicated as hub genes from PPI network, miRNA-hub gene regulatory network and TF-hub gene regulatory network. Furthermore, ROC analysis revealed that ESR2, JUN, PTN, PTGER3, CEBPB, IKBKG, HSPA8, SFN, CDKN1A and E2F1 might serve as diagnostic biomarkers in NAFLD. The current investigation provided insights into the molecular mechanism of NAFLD that might be useful in further investigations.

## Introduction

Non alcoholic fatty liver disease (NAFLD) has emerged as the main cause of metabolic disorder, but current treatments are still suboptimal [1]. NAFLD is a heterogeneous liver disorder that is characterized by hepatic steatosis to non-alcoholic steatohepatitis (NASH) and cirrhosis [2]. NAFLD are linked with progression of hepatocellular carcinoma [3], cardiovascular diseases [4], cerebrovascular diseases [5], diabetes mellitus [6], dietary rhythm [7], obesity [8], hypertension [9] and renal diseases [10]. Genetic factors can be used to show the pathogenesis of NAFLD [11]. Our understanding of the occurrence and advancement molecular mechanism of NAFLD has been greatly upgraded; however, the cause and potential molecular pathogenesis of NAFLD is still unclear [12]. Therefore, it is necessary to explore novel hub genes and signaling pathways for understanding the molecular mechanism and discovering key biomarkers for NAFLD.

At present, next generation sequencing (NGS) technology is widely used in molecular mechanism study and has a broad range of application in molecular biology [13]. It offers an efficient method for systematically screening NAFLD genes and signaling pathways, and identifying their regulatory mechanisms with bioinformatics [14]. Genes includes heme oxygenase-1 [15], odd-skipped related 1 (Osr1) [16], serine rich splicing factor 3 (SRSF3) [17], lipocalin-type prostaglandin D2 synthase (L-PGDS) [18] and β-Klotho (KLB) [19] are responsible for advancement of NAFLD. Signaling pathways includes LKB1/AMPK signaling pathway [20], Nrf2 signaling pathway [21], Sirt1/AMPK and NF-κB signaling pathways [22], eIF2α signaling pathway [23] and AMPK and LXR signaling pathways [24] are liable for progression of NAFLD. Therefore, a plenty of valuable clues could be explored for novel studies on the base of these data. Furthermore, many bioinformatics investigation on NAFLD have been produced in recent years [25], which proved that the integrated bioinformatics methods could help us to further investigation and better exploring the molecular mechanisms.

We downloaded NGS dataset GSE135251 [26–27] from Gene Expression Omnibus database (GEO) database (http://www.ncbi.nlm.nih.gov/geo/) [28], which contain expression profiling by high throughput sequencing data from NAFLD samples and normal control samples. We then performed deep bioinformatics analysis, including identifying common DEGs, gene ontology (GO) and pathway enrichment analysis, protein-protein interaction (PPI) network analysis, module analysis, construction and analysis of miRNA-hub gene regulatory network and TF-hub gene regulatory network. The findings were further validated by receiver operating characteristic curve (ROC) analysis. The aim of this study was to identify DEGs and key signaling pathways, and to explore potential candidate biomarkers for the diagnosis and therapeutic targets in NAFLD.

## Materials and methods

### Data resources

Expression profiling by high throughput sequencing dataset GSE135251 [26–27] was downloaded from the GEO database. The data was produced using a GPL18573 Illumina NextSeq 500 (Homo sapiens). The GSE135251 dataset contained data from 216 samples, including 206 NAFLD samples and 10 normal control samples.

### Identification of DEGs

To assess DEGs, using the “DESeq2” package of R software [29]. A moderated t-statistic and a log-odds of DEG was computed for each contrast for each gene. The Benjamini and Hochberg (BH) method was performed to adjust P value to reduce the false positive error [30]. DEGs between NAFLD samples and normal control samples were identified via DESeq2 package of R software with |logFC|□ > 1.295 for up regulated genes, |logFC|□ < -1.368 for down regulated genes and adjust P value <□ 0.05, which were visualized as volcano plots and heat map. “ggplot2” and “gplot” packages of R software was applied to generate volcano plots and heat map.

### GO and pathway enrichment analyses of DEGs

g:Profiler (http://biit.cs.ut.ee/gprofiler/) [31] is an online functional annotation tool to provide a comprehensive understanding of biological information of genes and proteins. GO enrichment analysis (http://www.geneontology.org) [32] is a commonly used approach for defining genes and its RNA or protein product to identify unique biological properties of high-throughput genomic data. GO enrichment analysis including biological process (BP), cellular components (CC), and molecular function (MF). REACTOME (https://reactome.org/) [33] pathway database is a collection of databases dealing with genomes, diseases, biological pathways, drugs, and chemical materials. P value□<□0.05 was considered as the threshold.

### Construction of the PPI network and module analysis

The online Hippie interactome (http://cbdm-01.zdv.uni-mainz.de/~mschaefer/hippie/) [34] database was used to identify potential interaction among the common DEGs. Cytoscape software version 3.8.2 (http://www.cytoscape.org/) [35] was applied to establish the protein interaction network. The Network Analyzer plug-in was used to explore hub genes were generated using node degree [36], betweenness centrality [37], stress centrality [38] and closeness centrality [39]. The intersect function was used to identify the hub genes. Then, PEWCC1 plug-in [40] was performed to monitor PPI network modules with Cytoscape. The functional enhancement analysis of DEGs in the significant module were performed using g:Profiler.

### MiRNA-hub gene regulatory network construction

miRNet database (https://www.mirnet.ca/) [41] is a bioinformatics platform for predicting miRNA-hub gene pairs. In the current investigation, the targets of the hub genes were predicted using fourteen databases: TarBase, miRTarBase, miRecords, miRanda (S mansoni only), miR2Disease, HMDD, PhenomiR, SM2miR, PharmacomiR, EpimiR, starBase, TransmiR, ADmiRE, and TAM 2.0. The screening criterion was that the miRNA target exists in the fourteen databases concurrently. The miRNA-hub gene regulatory network was depicted and visualized using Cytoscape software [35].

### TF-hub gene regulatory network construction

NetworkAnalyst database (https://www.networkanalyst.ca/) [42] is a bioinformatics platform for predicting TF-hub gene pairs. In the current investigation, the targets of the hub genes were predicted using ChEA database. The screening criterion was that the TF target exists in the ChEA database concurrently. The TF-hub gene regulatory network was depicted and visualized using Cytoscape software [35].

### Receiver operating characteristic curve (ROC) analysis

ROC curve analysis was accomplished to assess the sensitivity (true positive rate) and specificity (true negative rate) of the hub genes for NAFLD diagnosis and we studied how large the area under the curve (AUC) was by using the pROC package in R statistical software [43].

## Results

### Identification of DEGs

We obtained Expression profiling by high throughput sequencing from newly diagnosed patients with NAFLD and normal controls from the GSE135251 and analyzed DEGs using DESeq2 package of R software. Setting the cut□off criterion as adjust P value <□0.05, |logFC|□> 1.295 for up regulated genes and |logFC|□< -1.368 for down regulated genes, we identified 951 DEGs from GSE135251. Total 951 DEGs, 476 were up regulated and 475 down regulated genes in NAFLD compared with normal control samples and are listed in Table 1. The heatmap and volcano plot are shown in Fig. 1 and and Fig. 2, respectively.

**Fig. 1.**
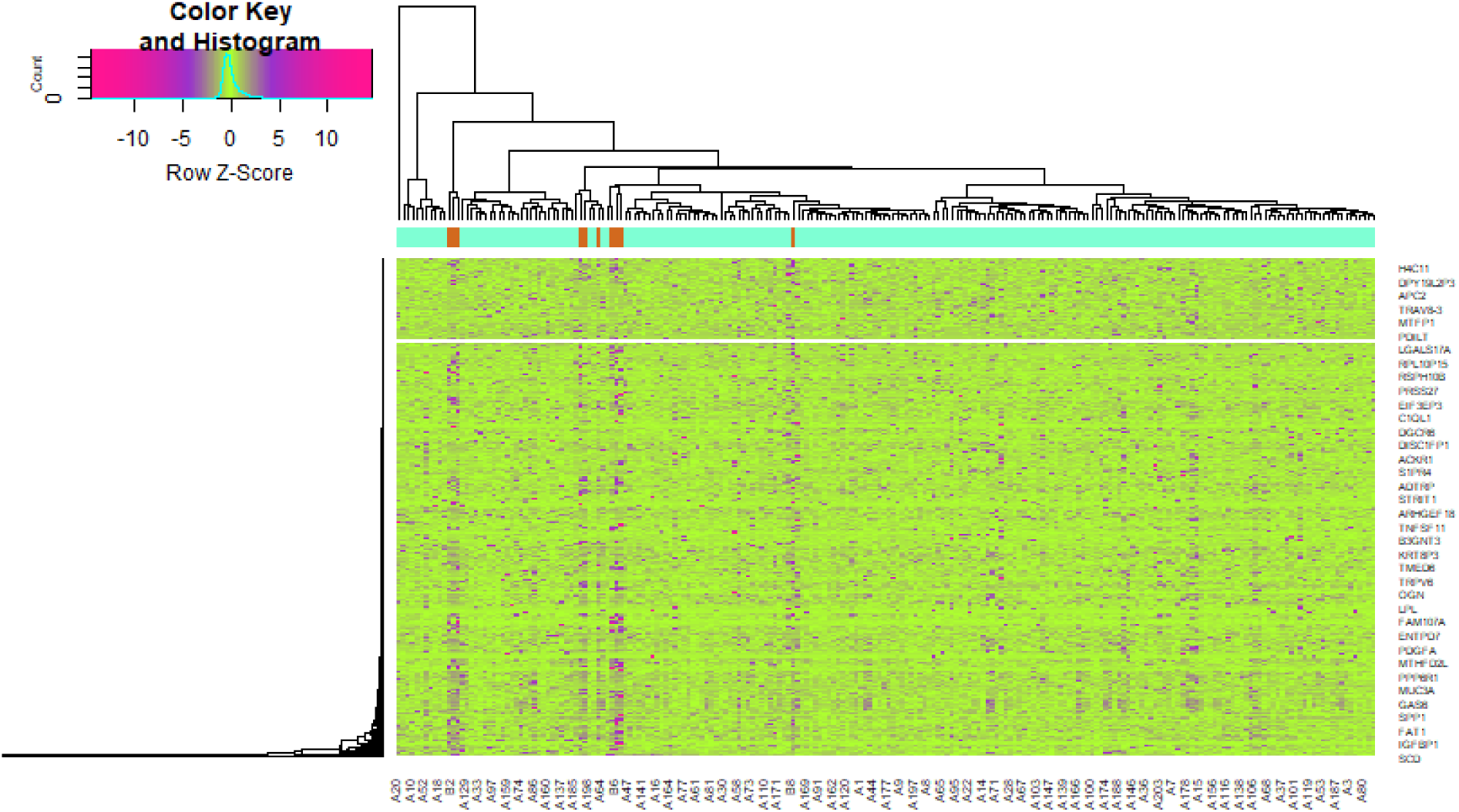
Heat map of differentially expressed genes. Legend on the top left indicate log fold change of genes. (A1 – A206= Non Alcoholic Fatty Liver Disease Samples; B1 – B10= Normal Control Samples)

**Fig. 2.**
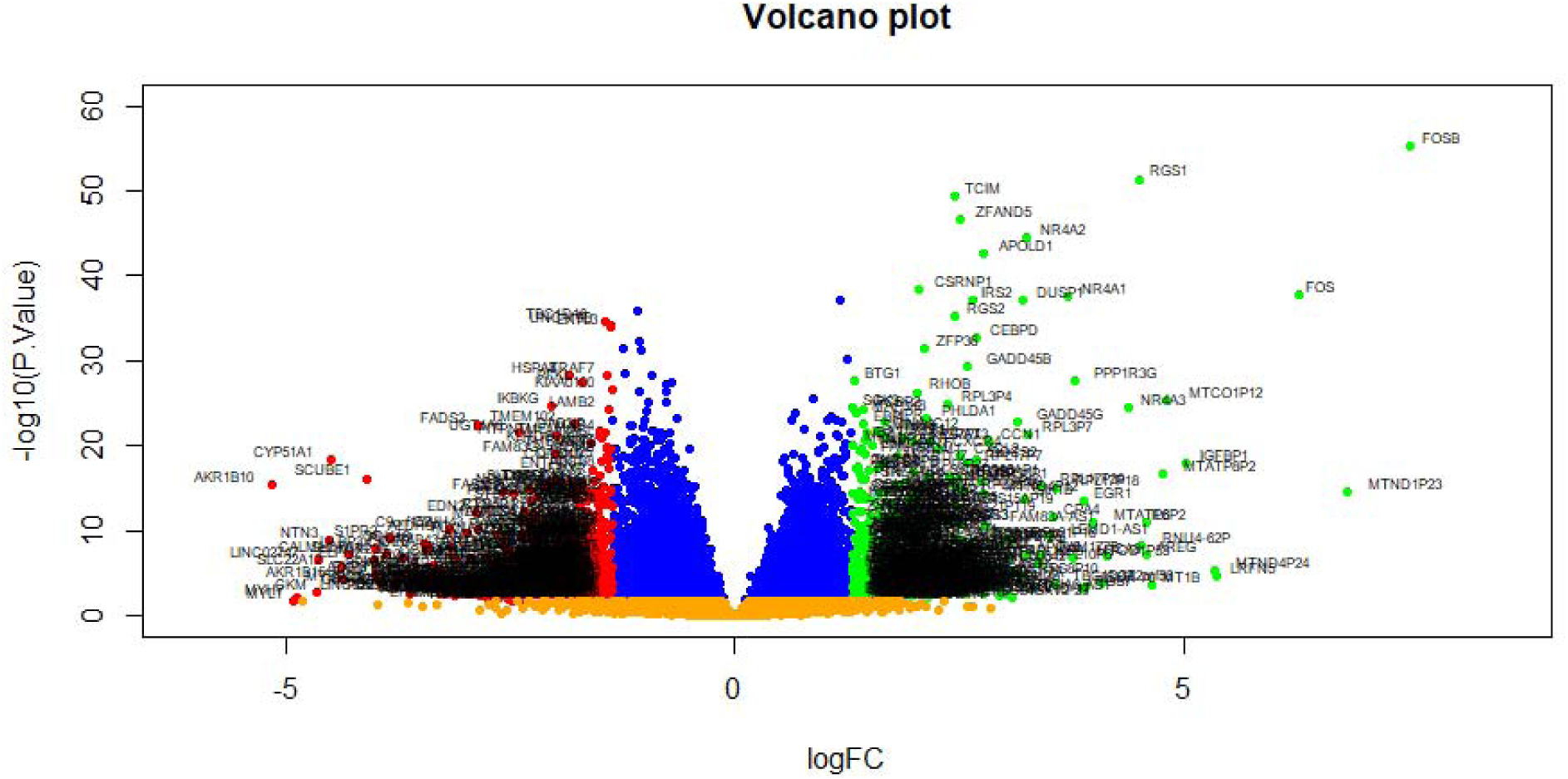
Volcano plot of differentially expressed genes. Genes with a significant change of more than two-fold were selected. Green dot represented up regulated significant genes and red dot represented down regulated significant genes.

**Table 1.**
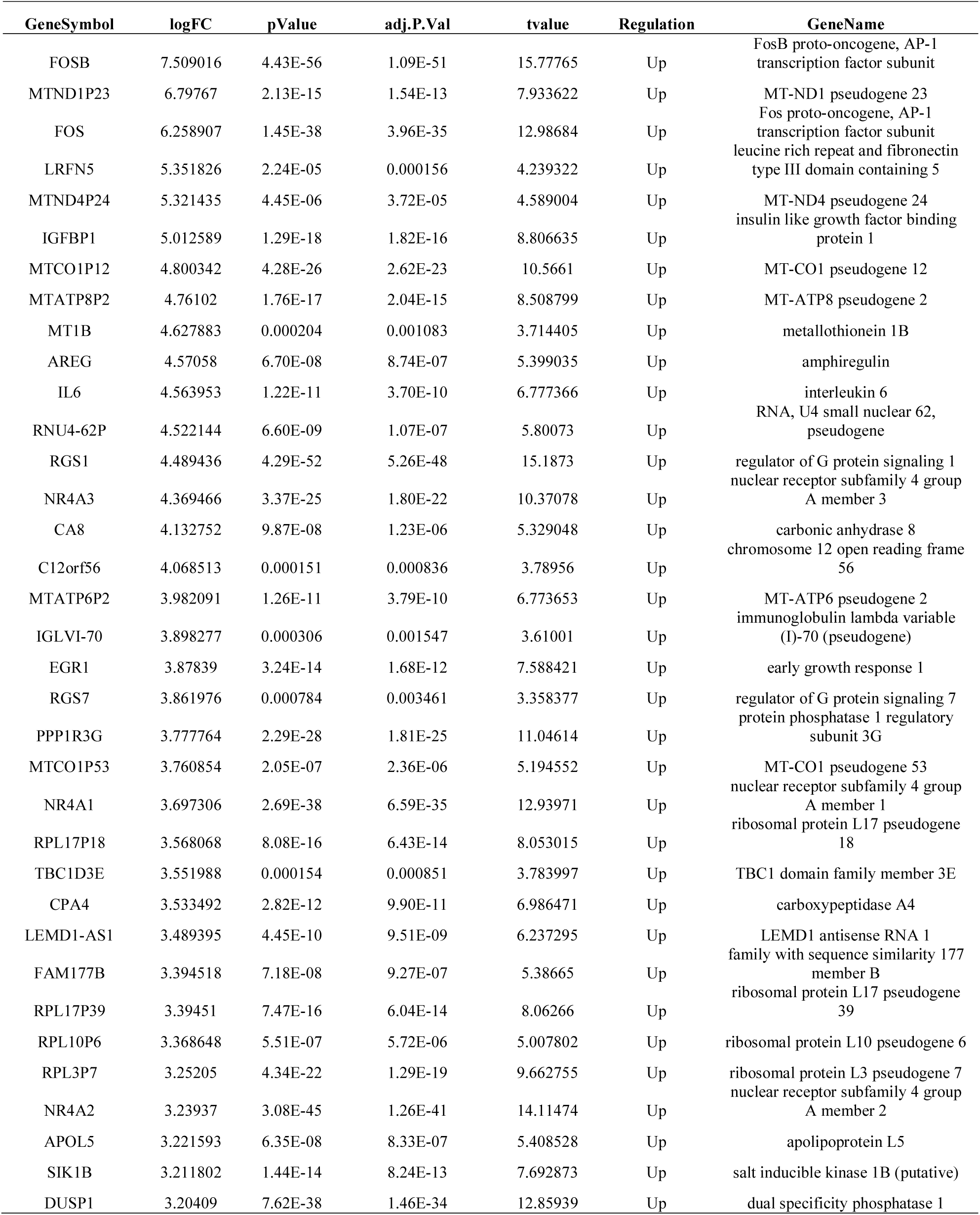
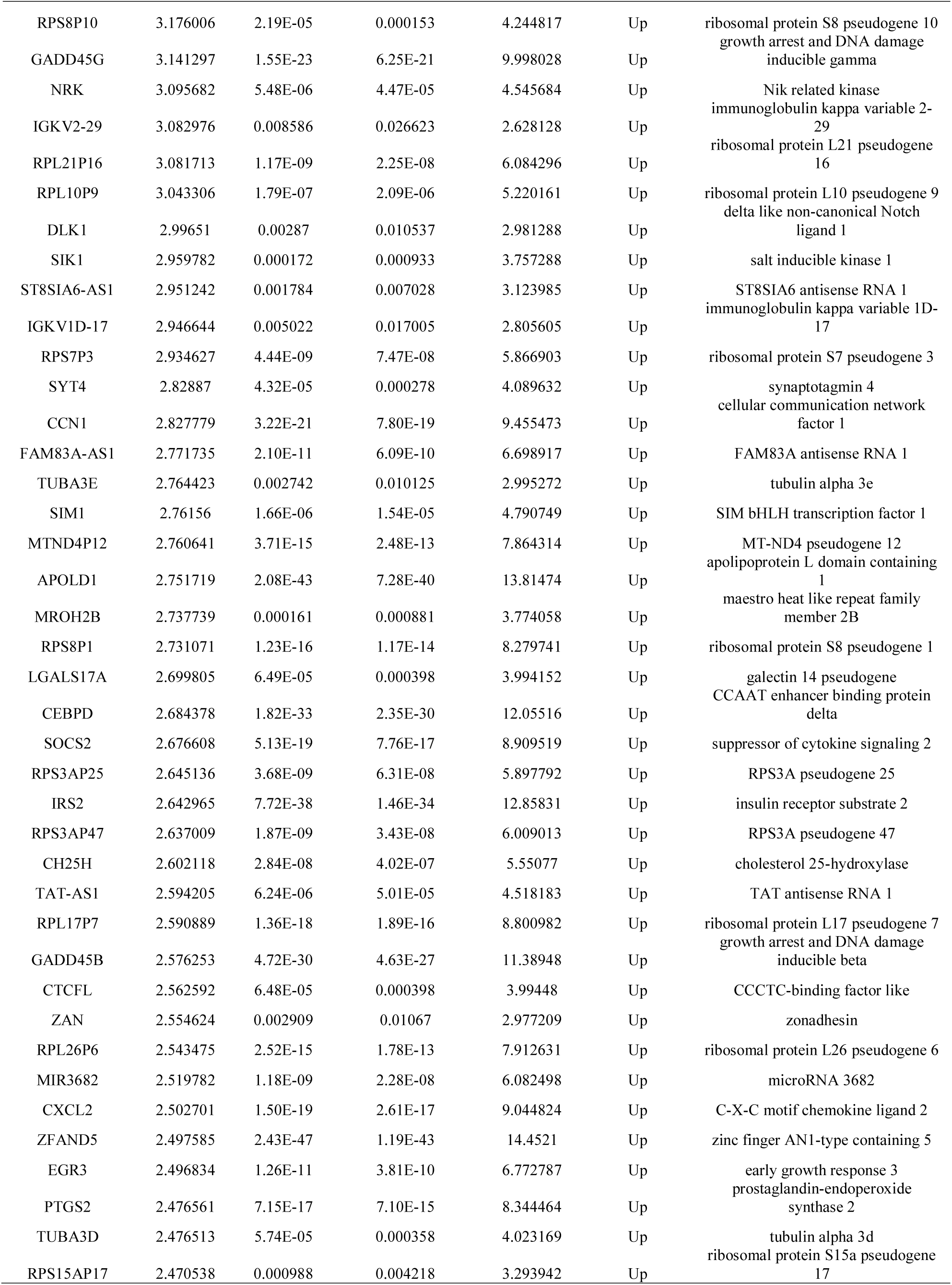
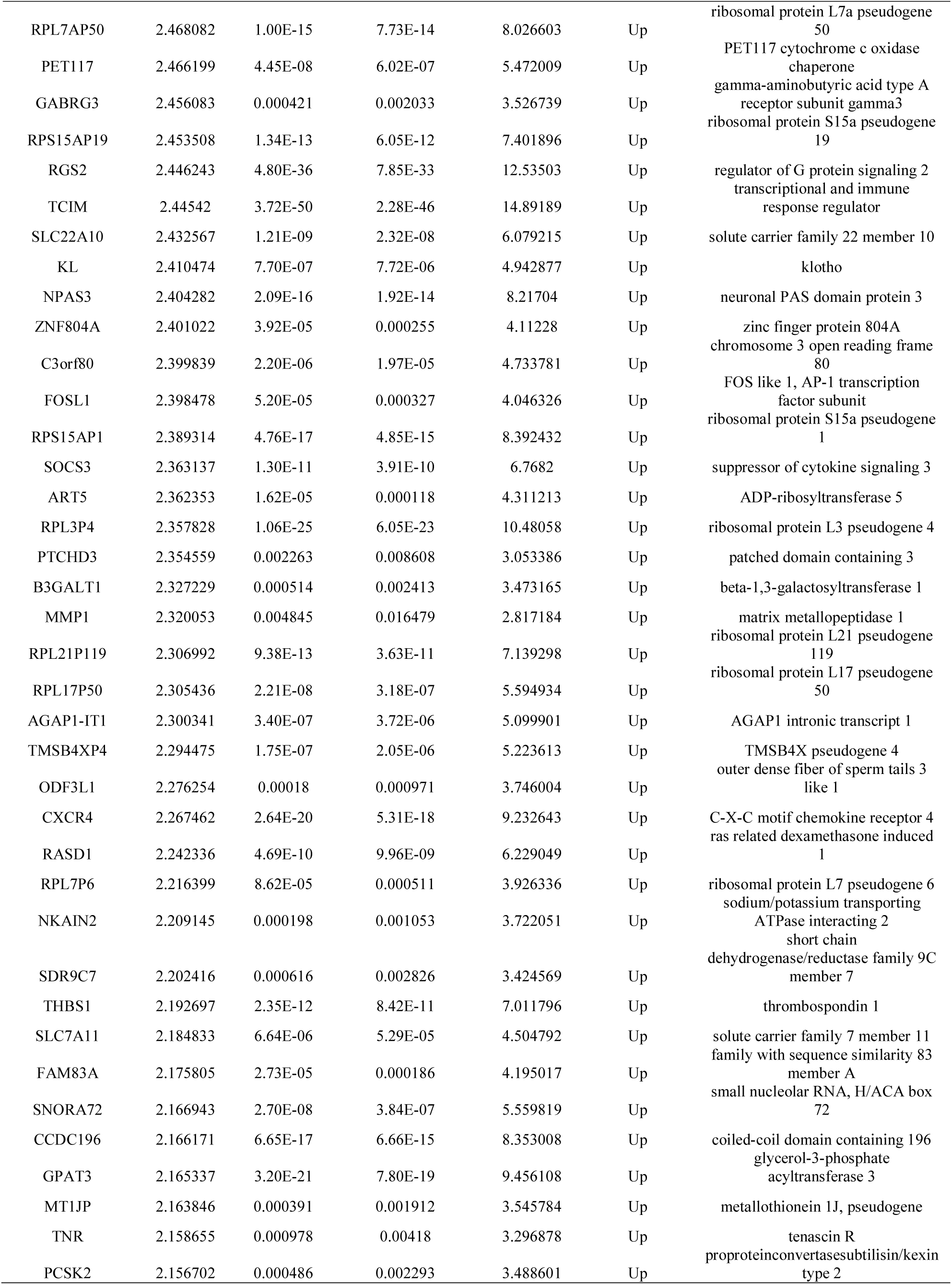
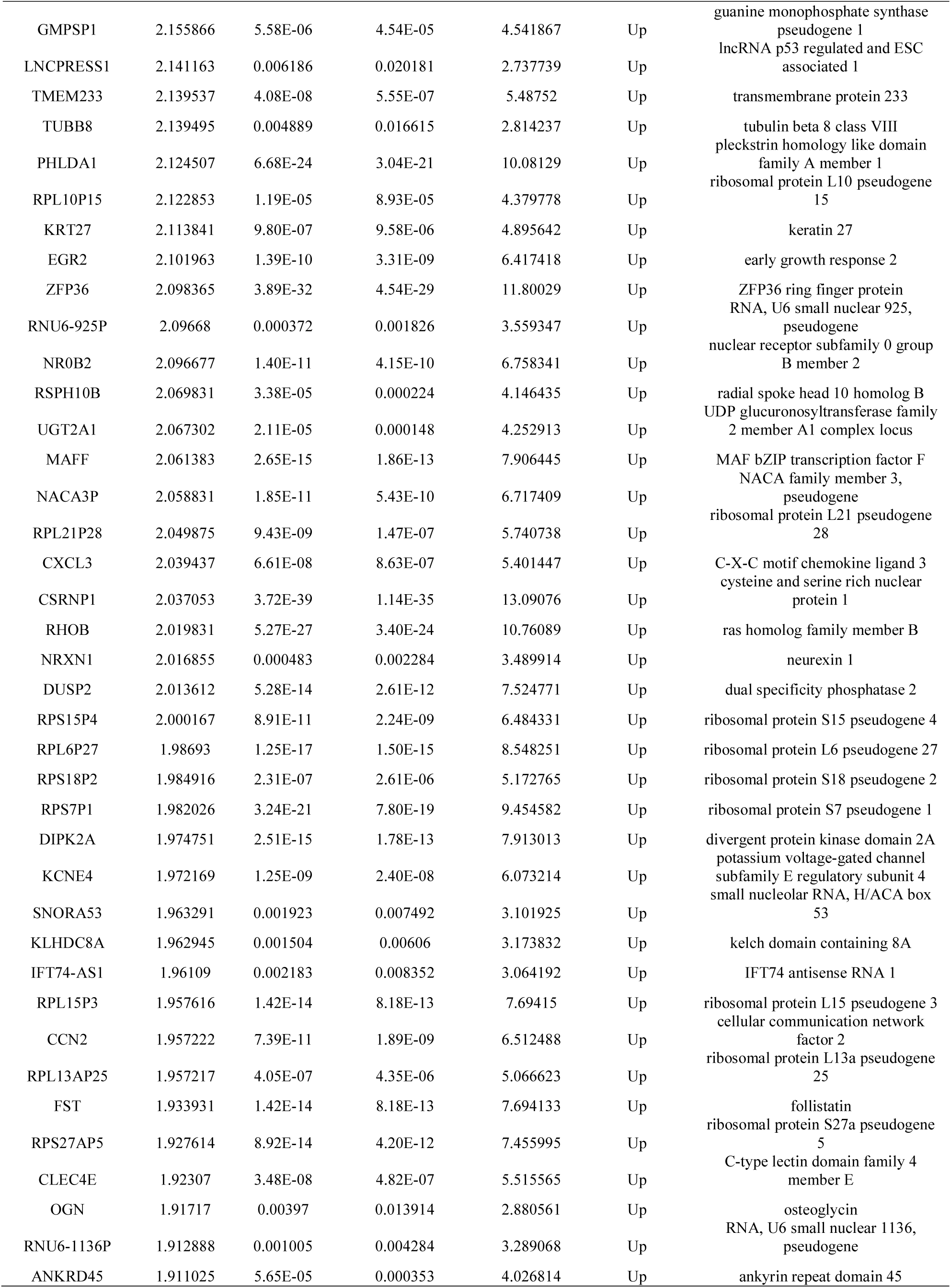
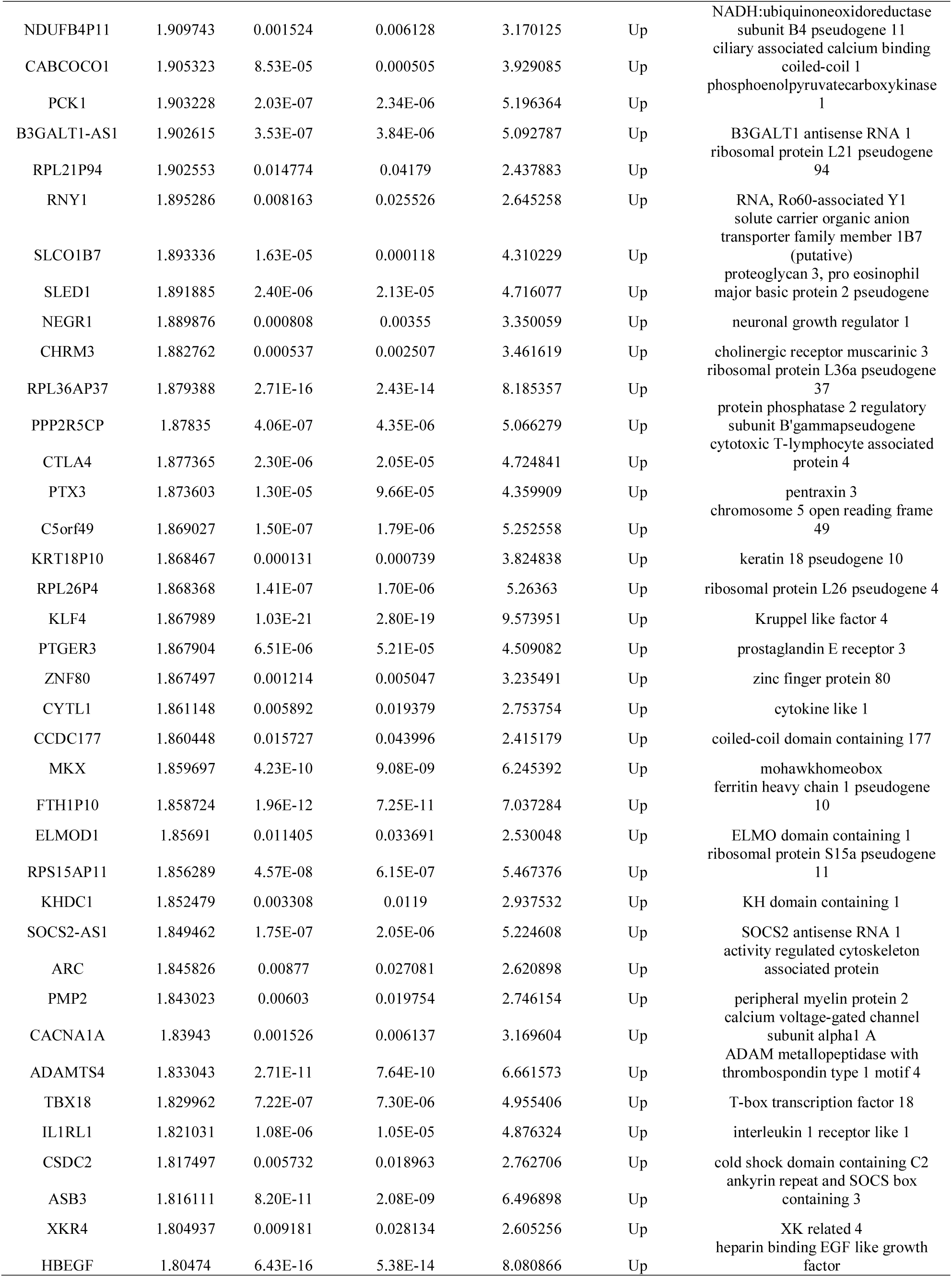
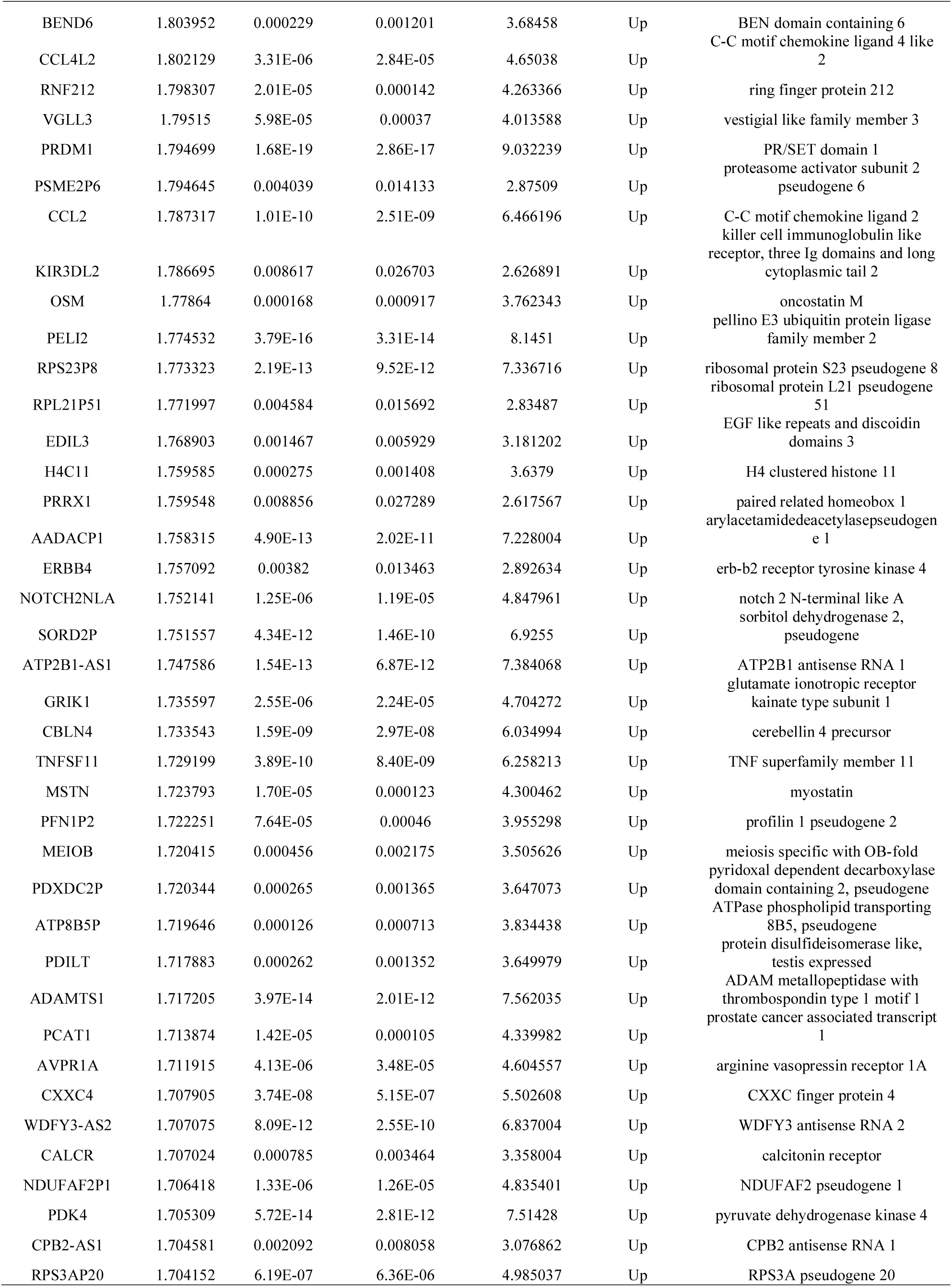
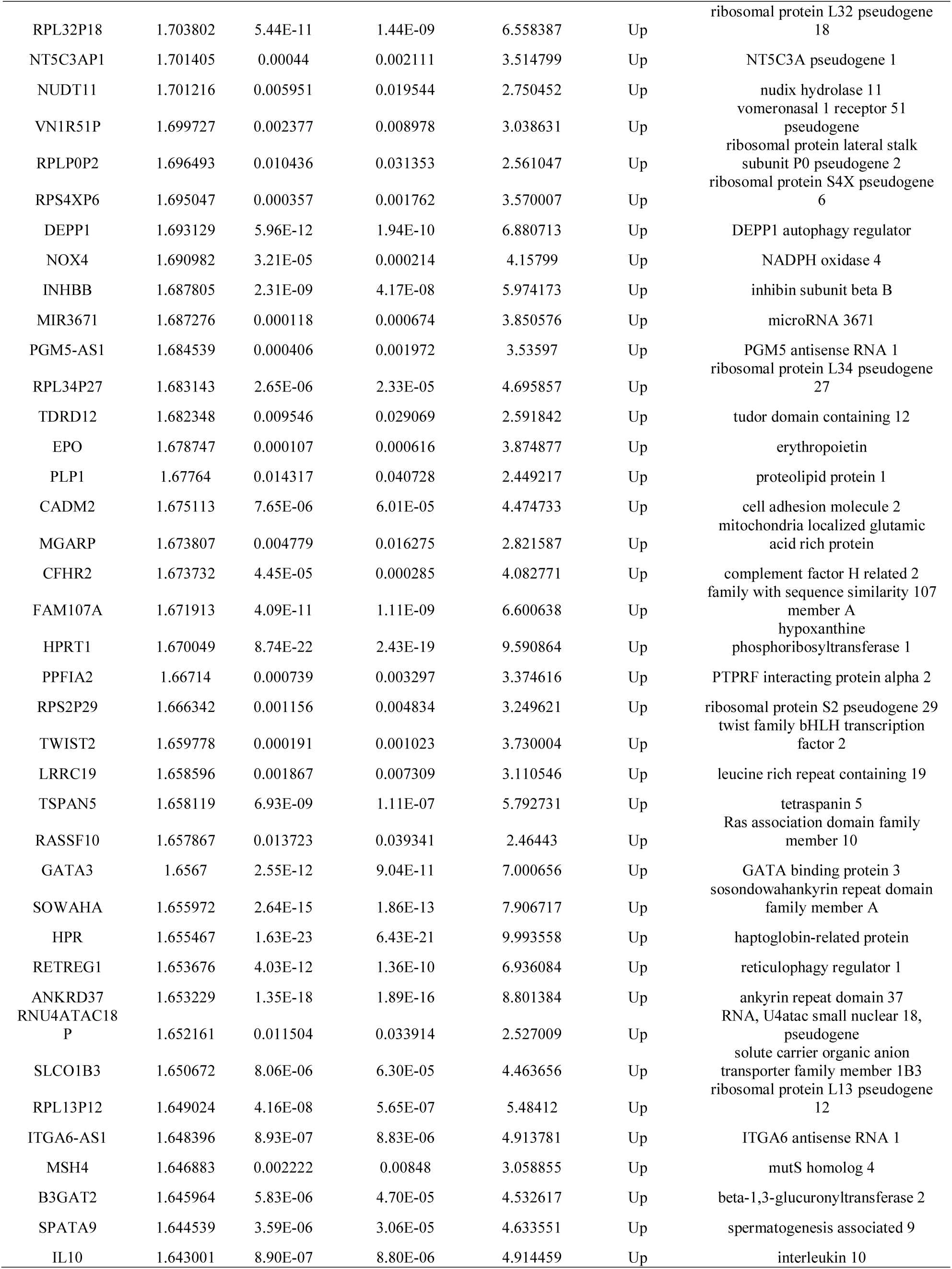
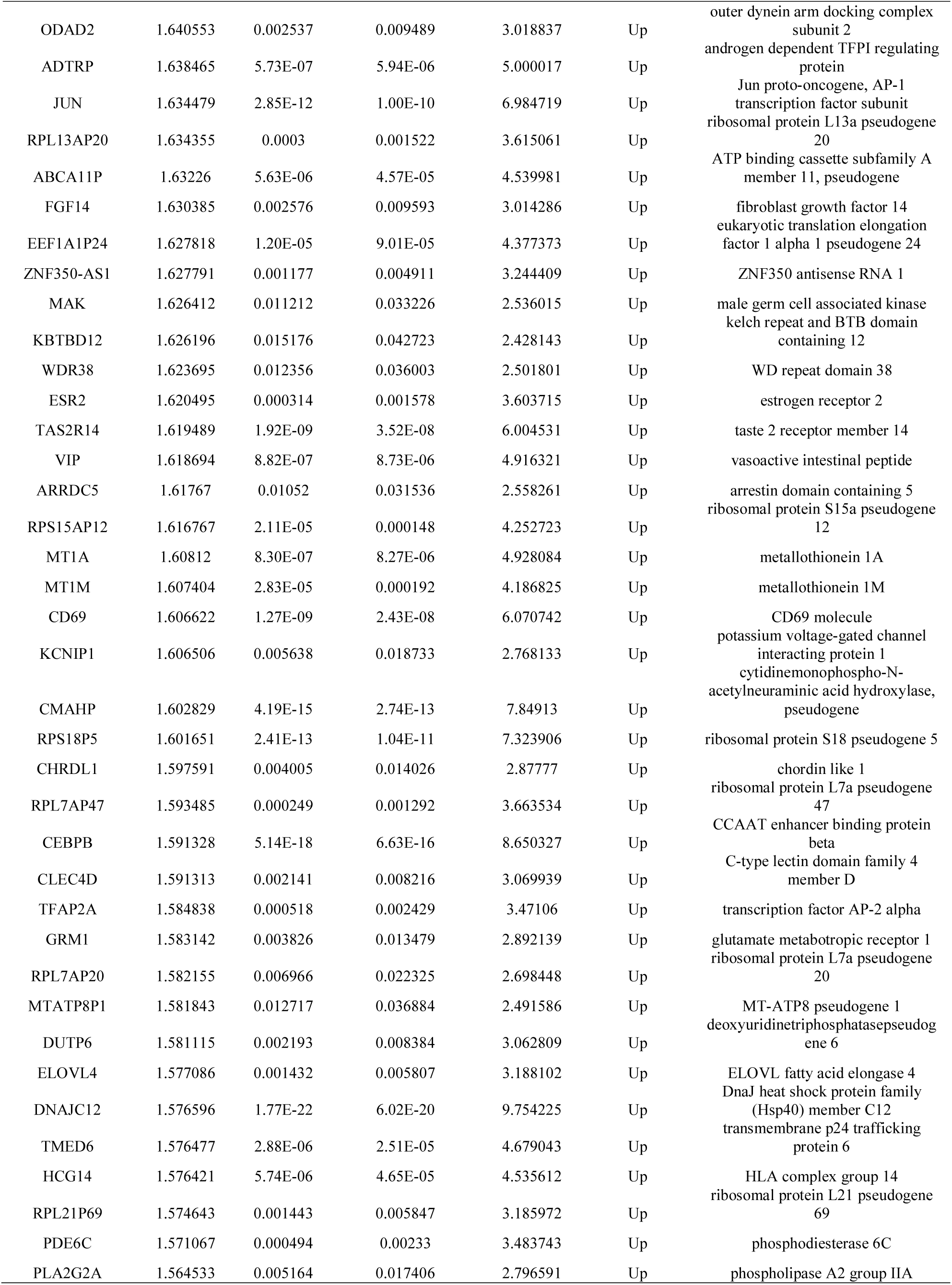
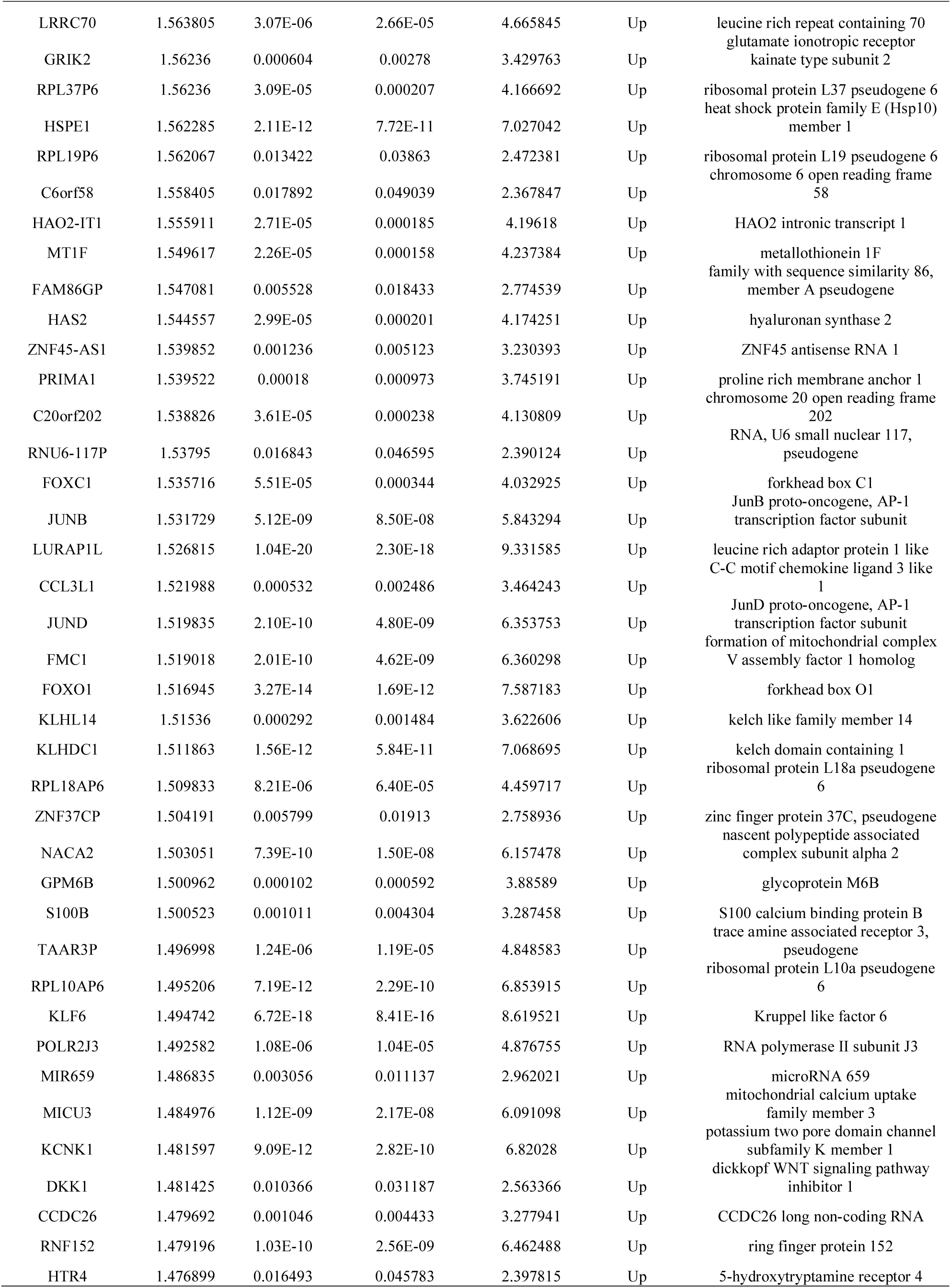
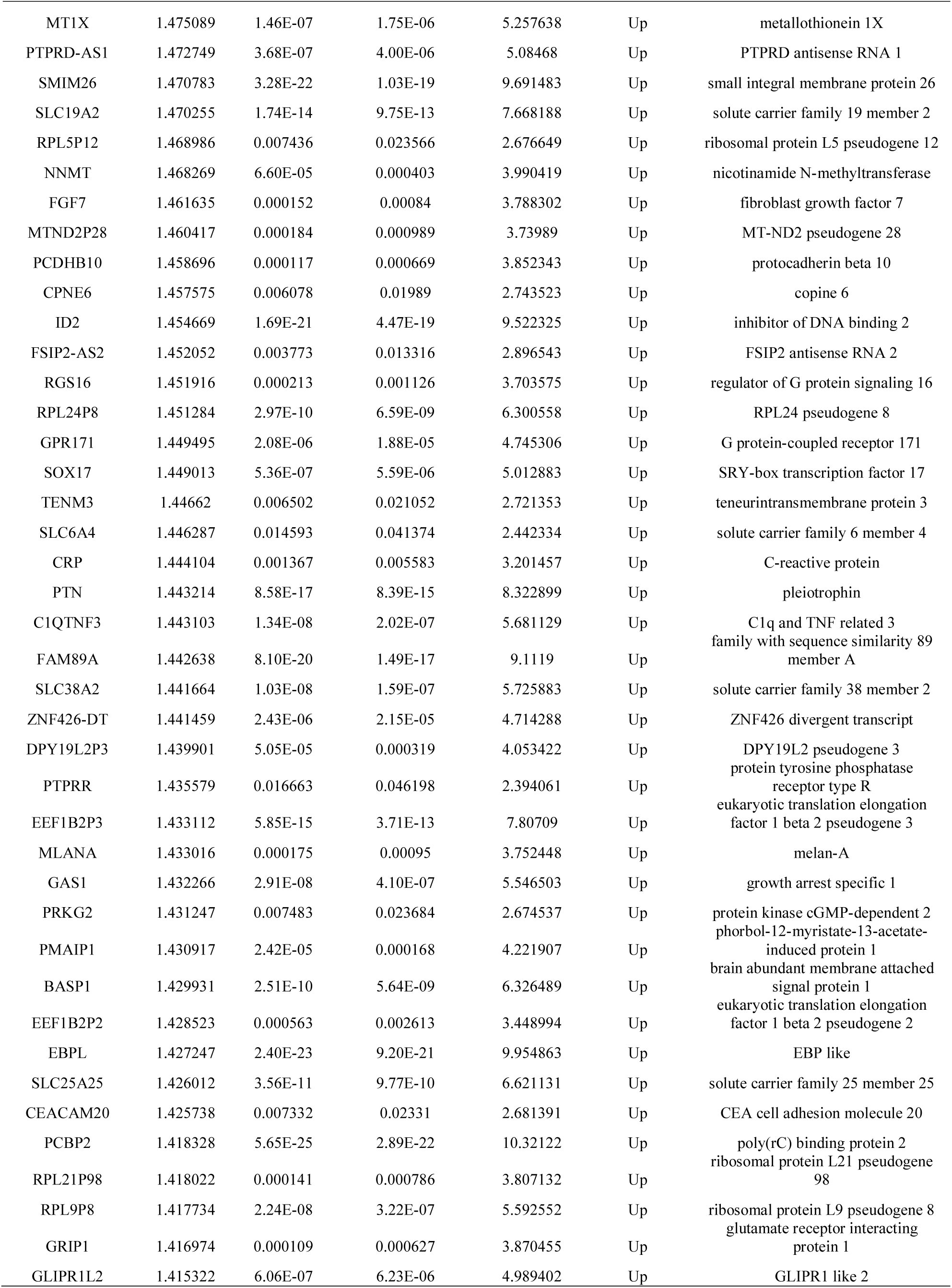
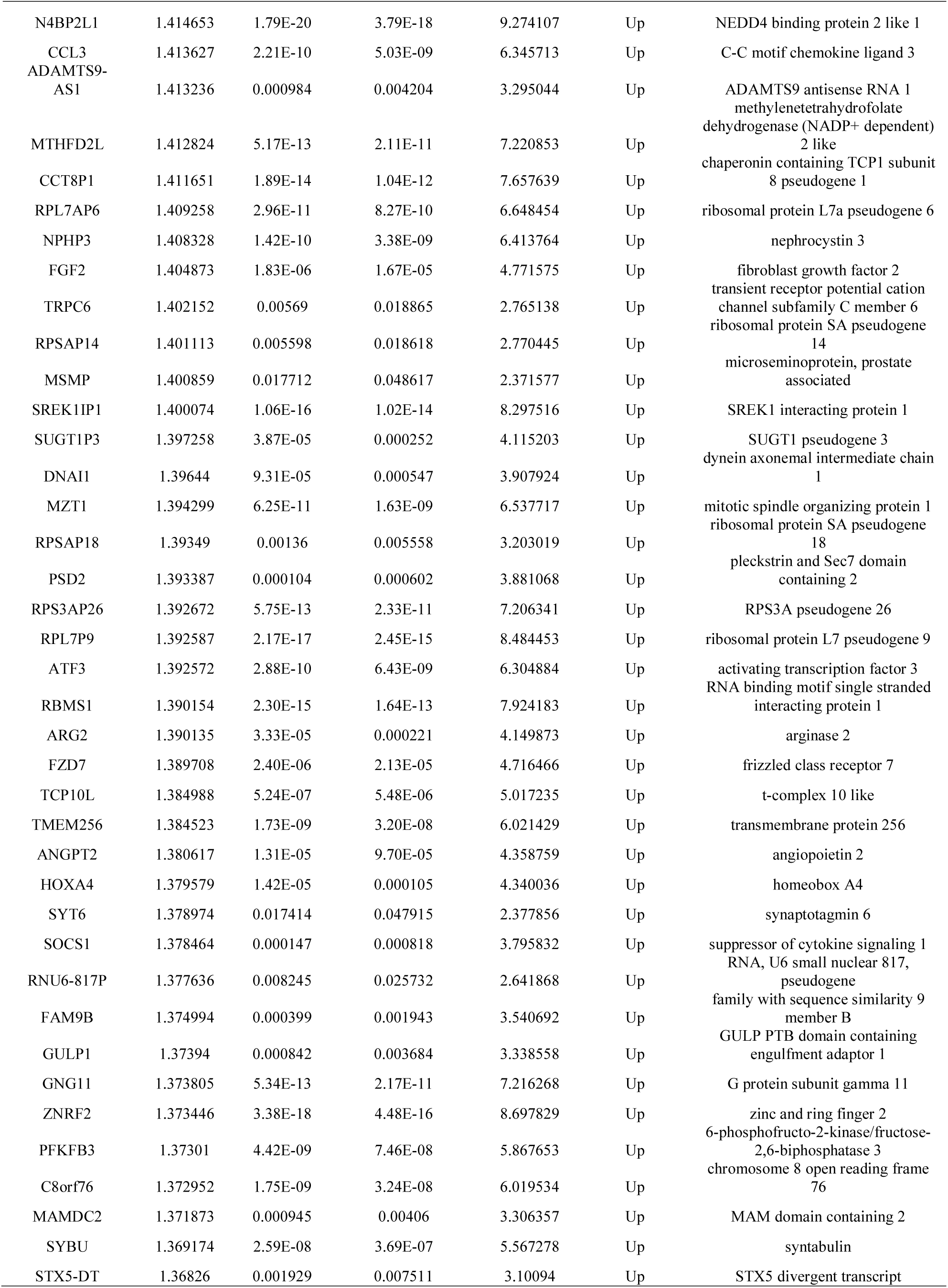
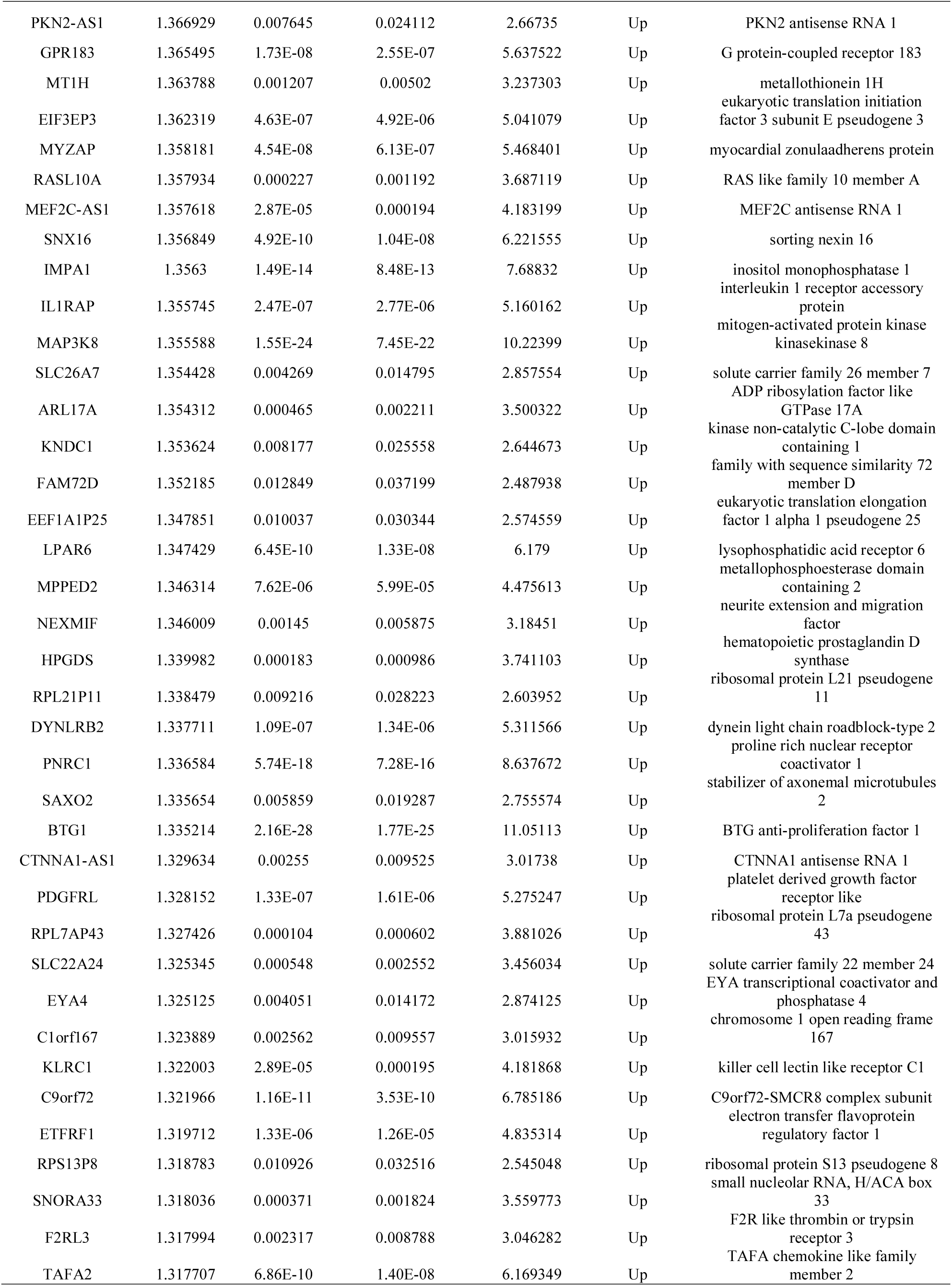
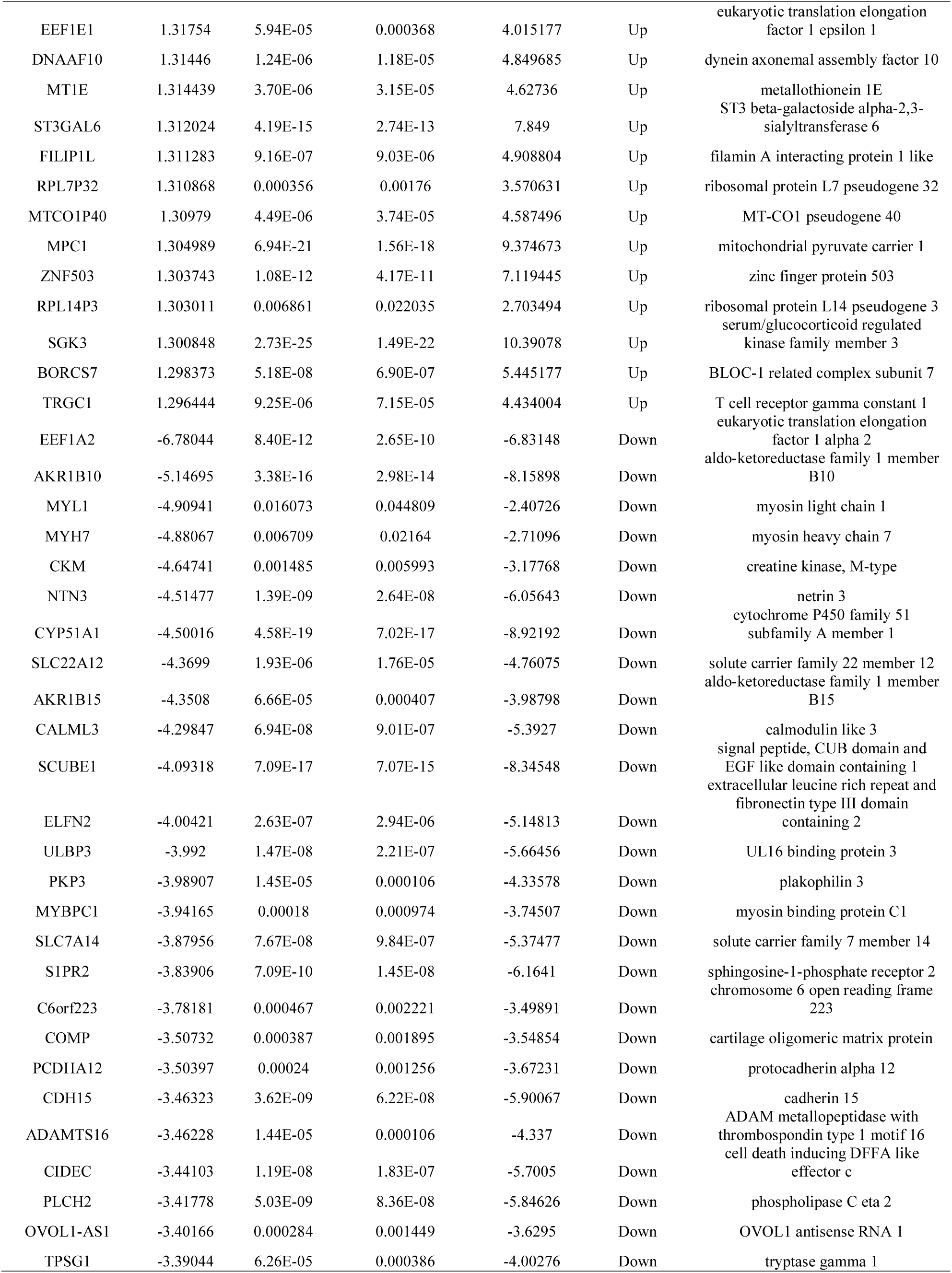
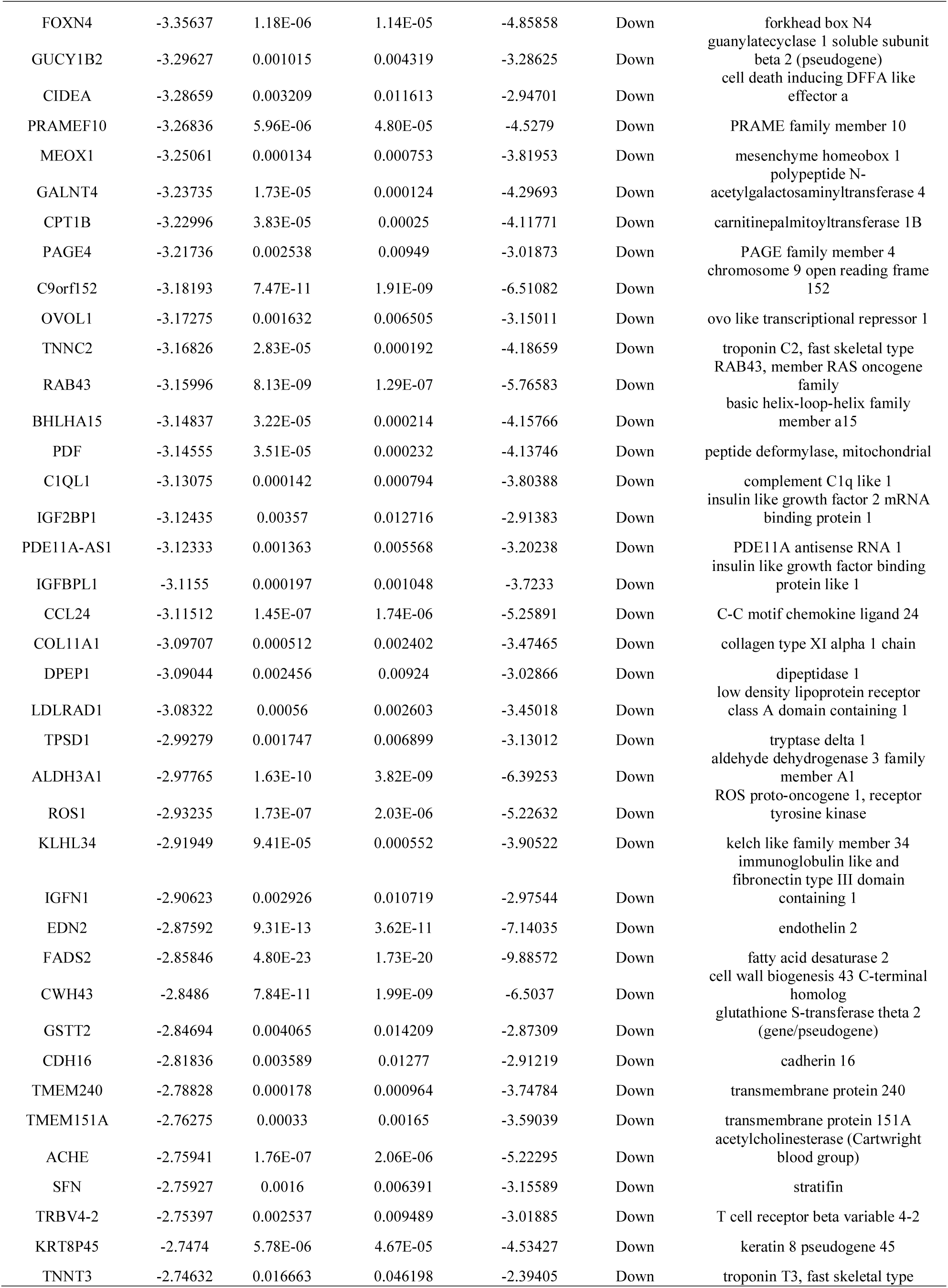
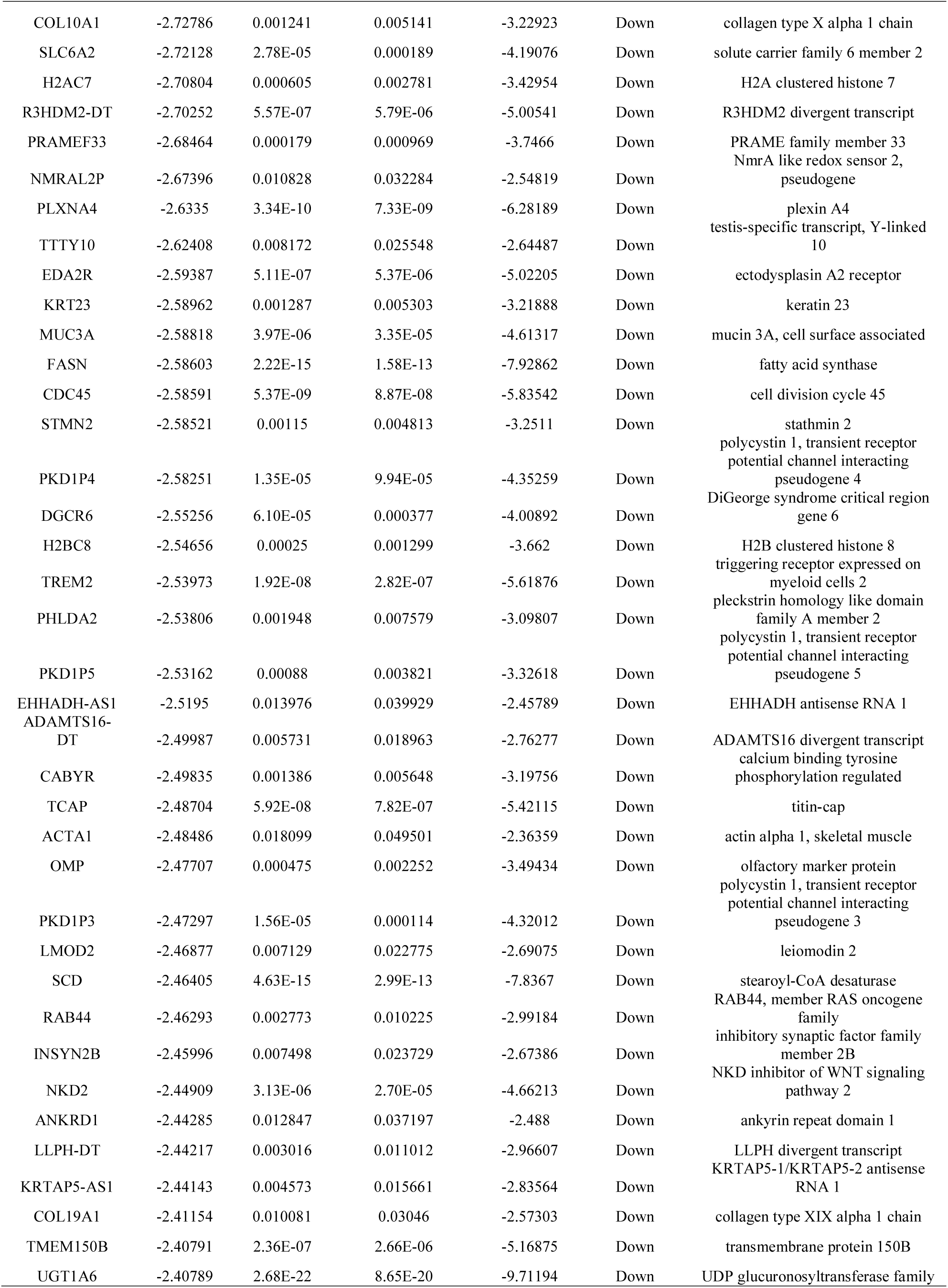
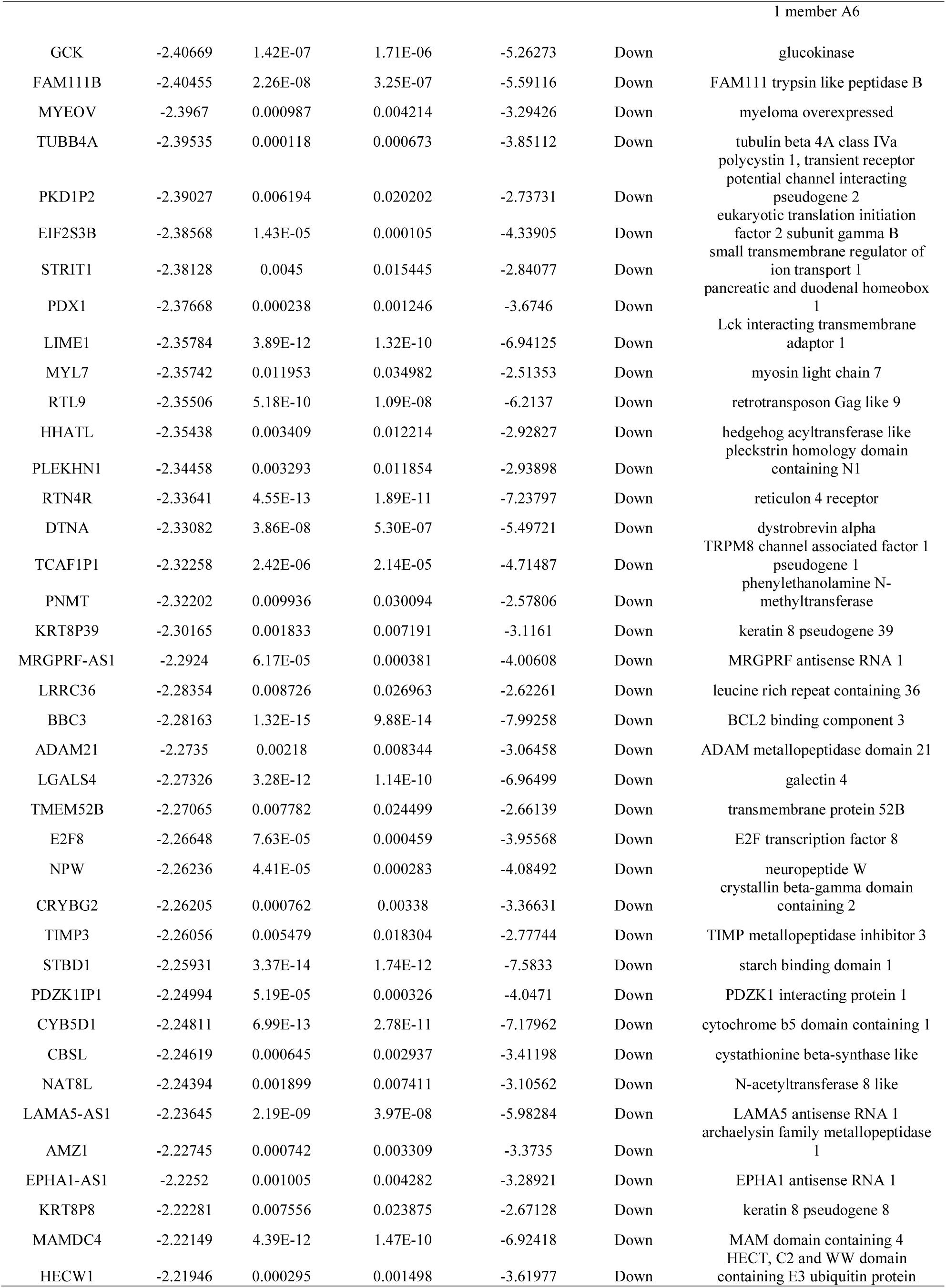
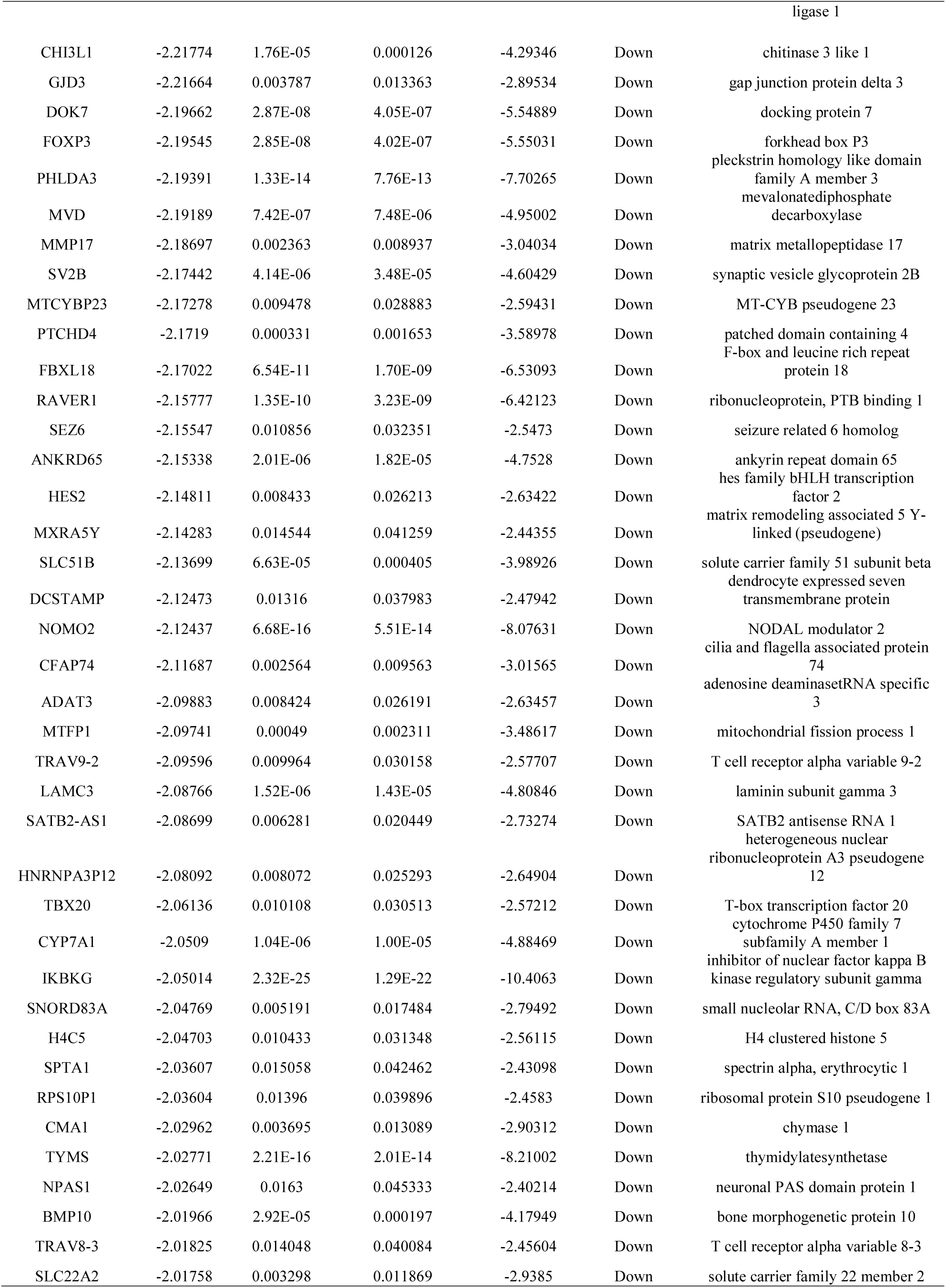
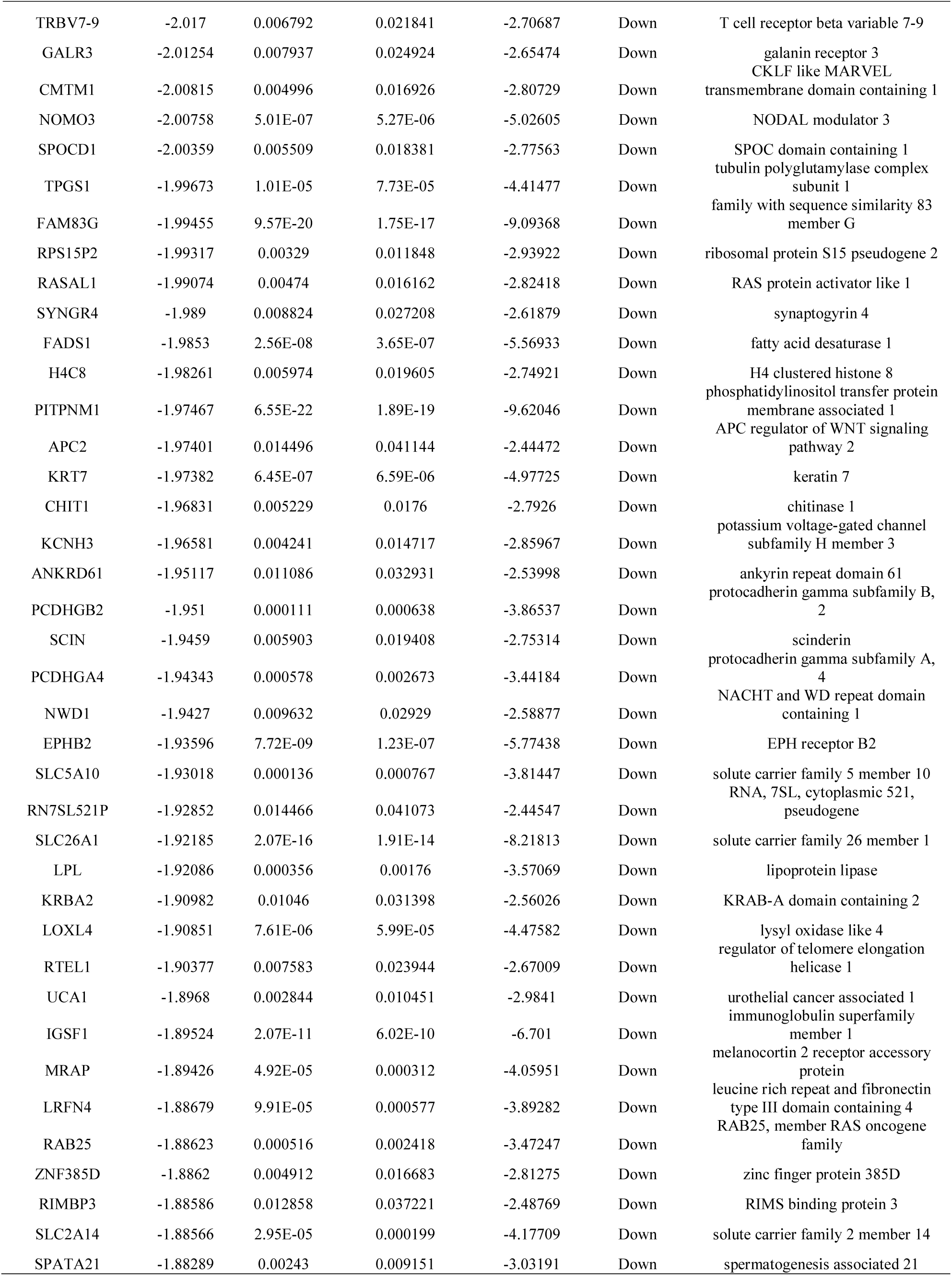
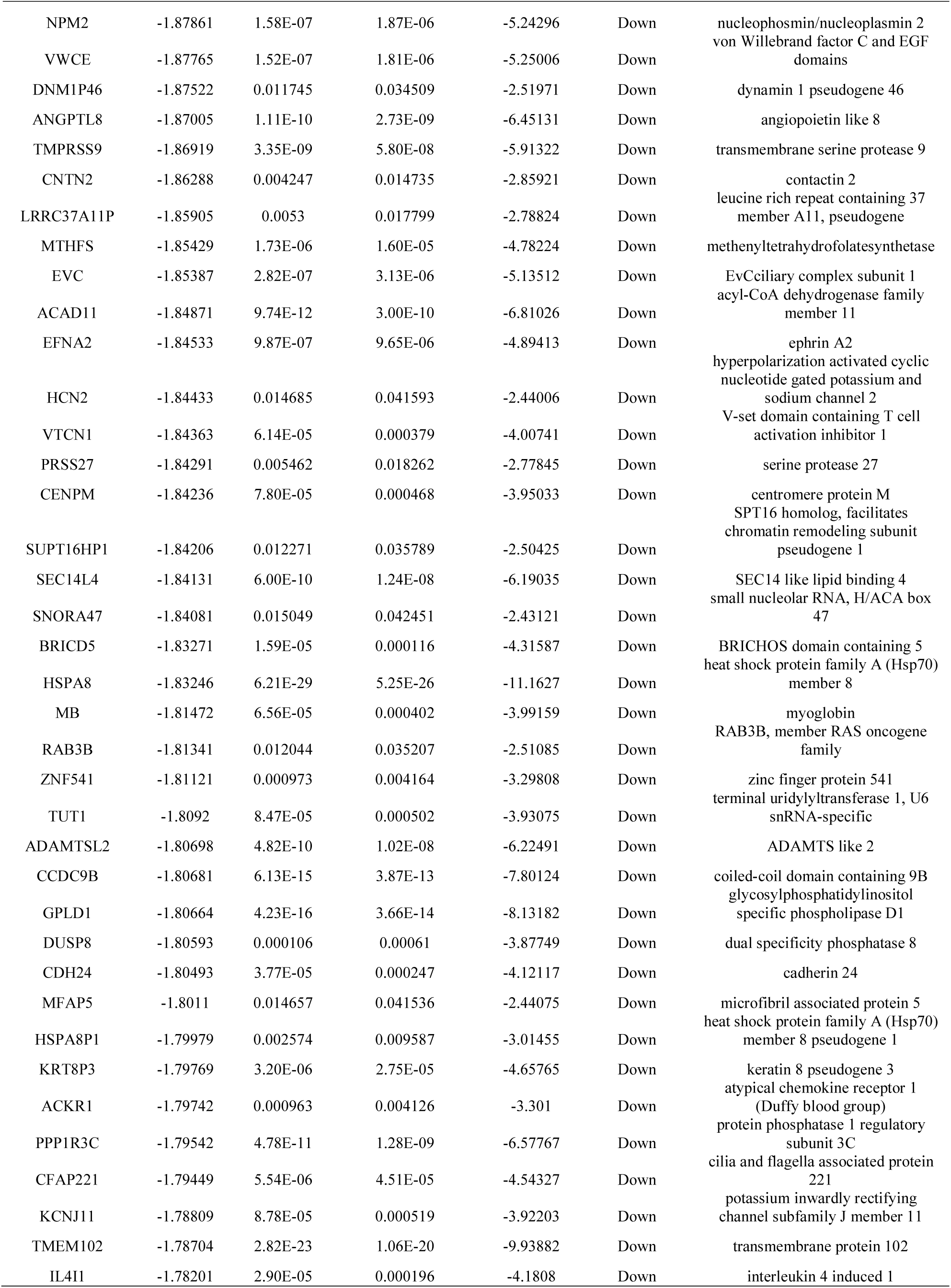
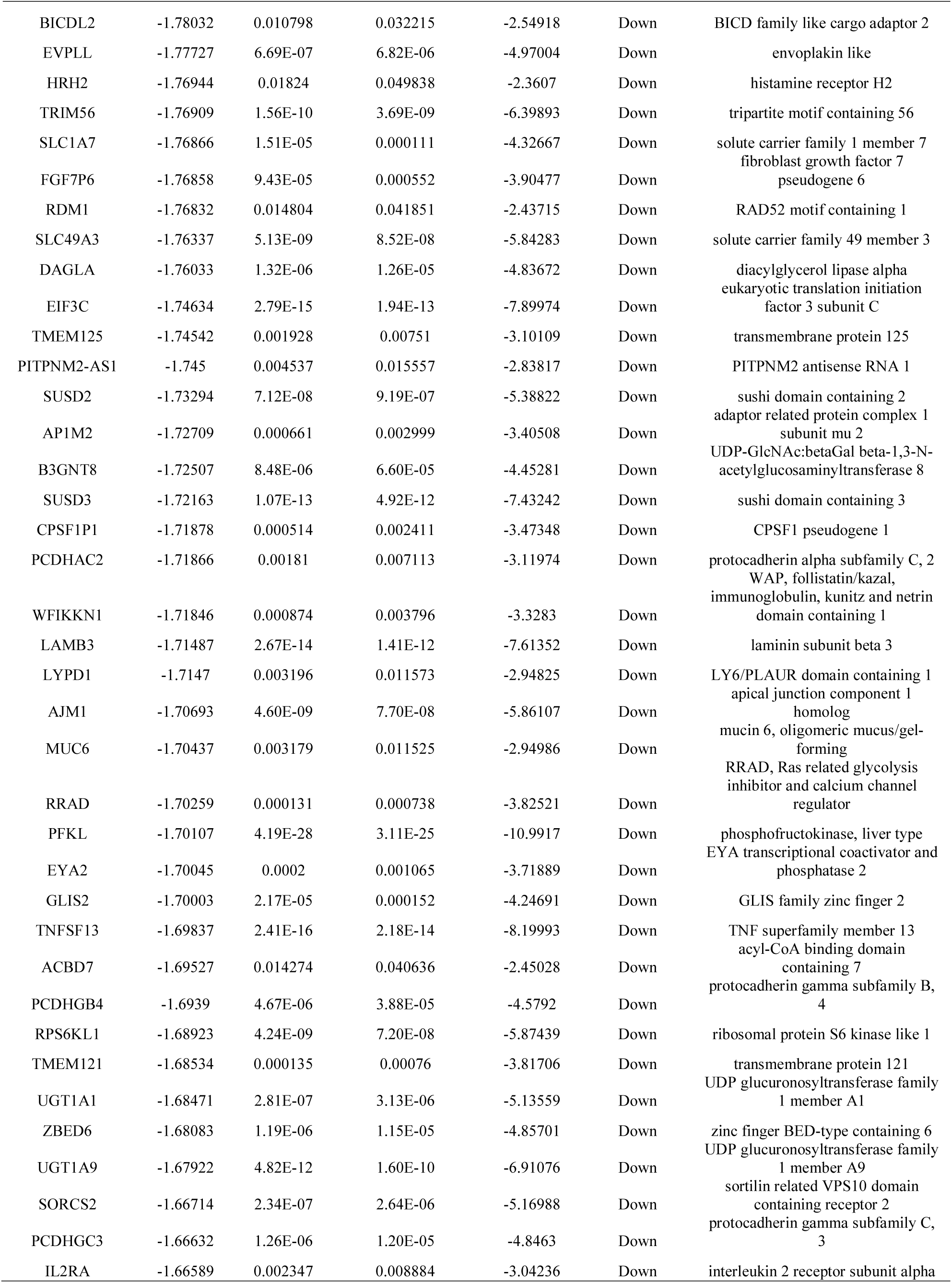
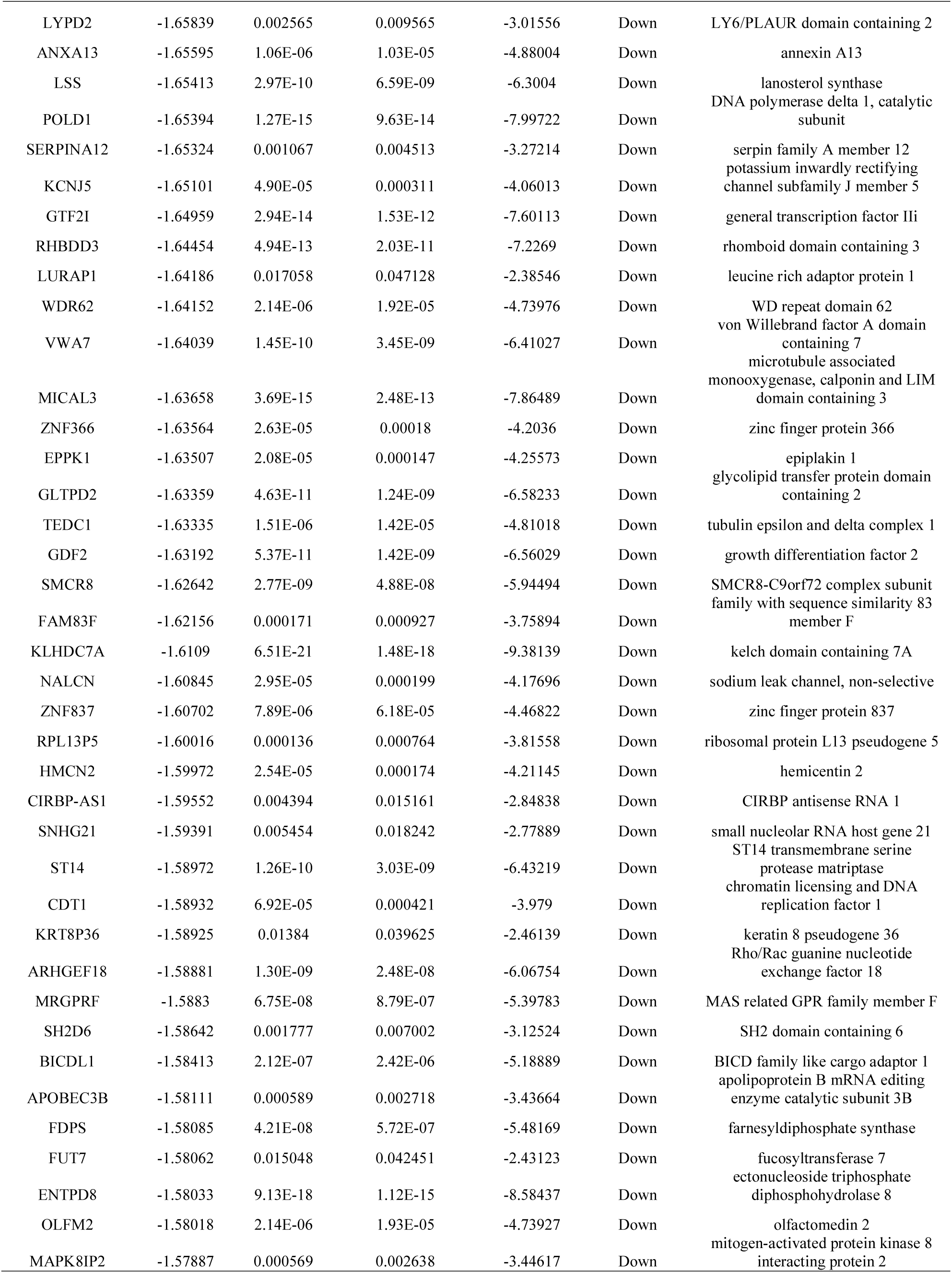
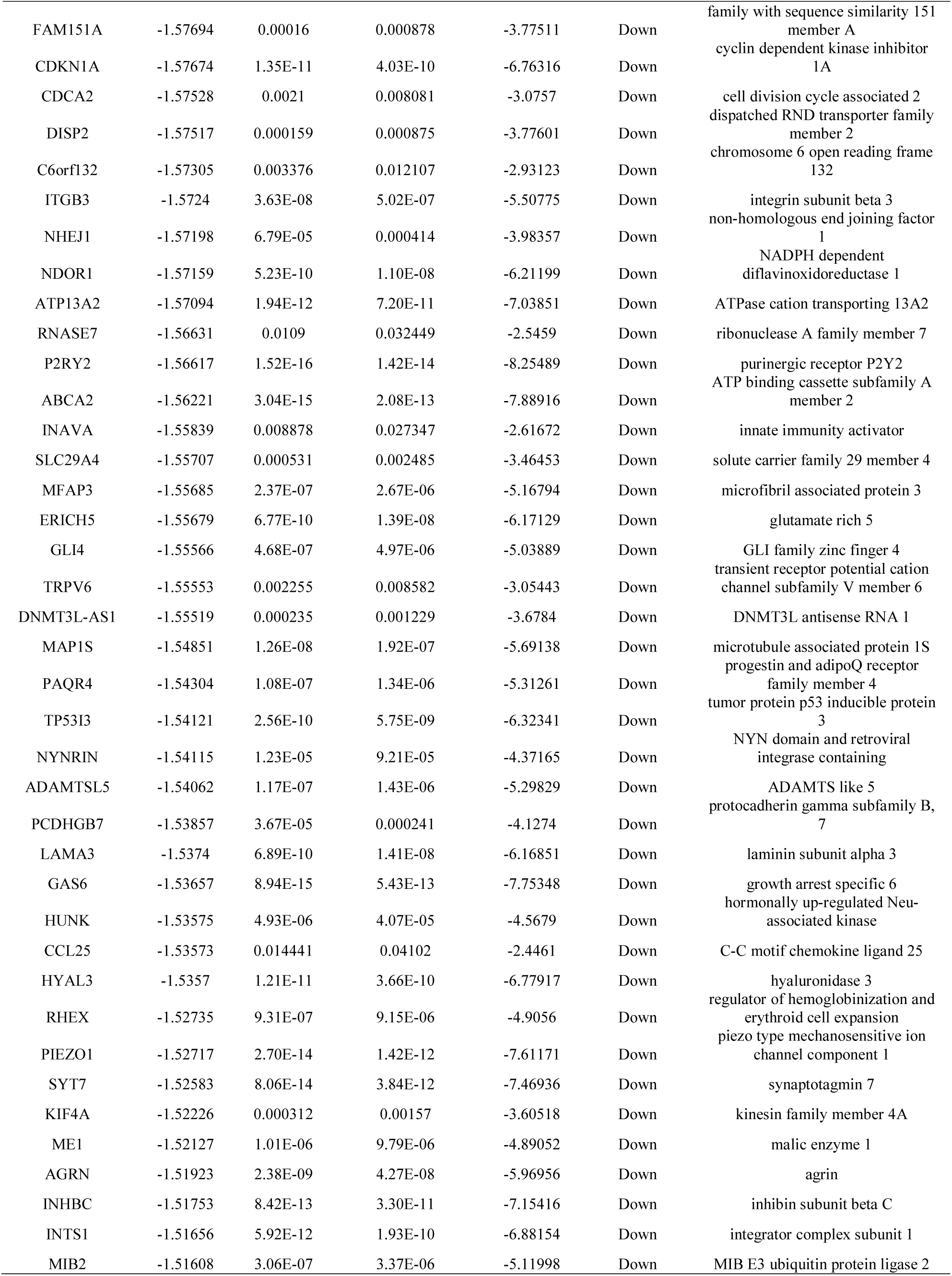
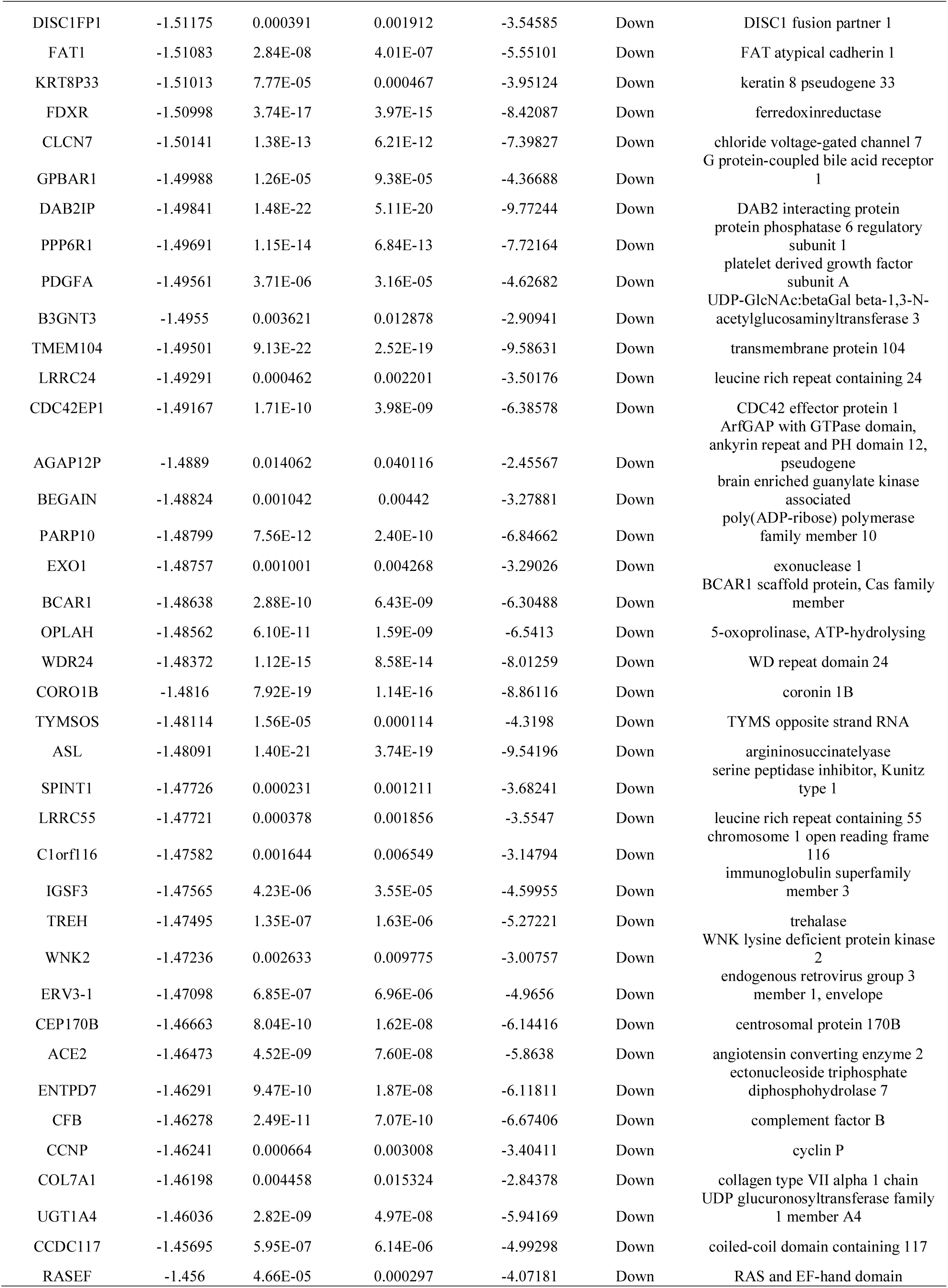
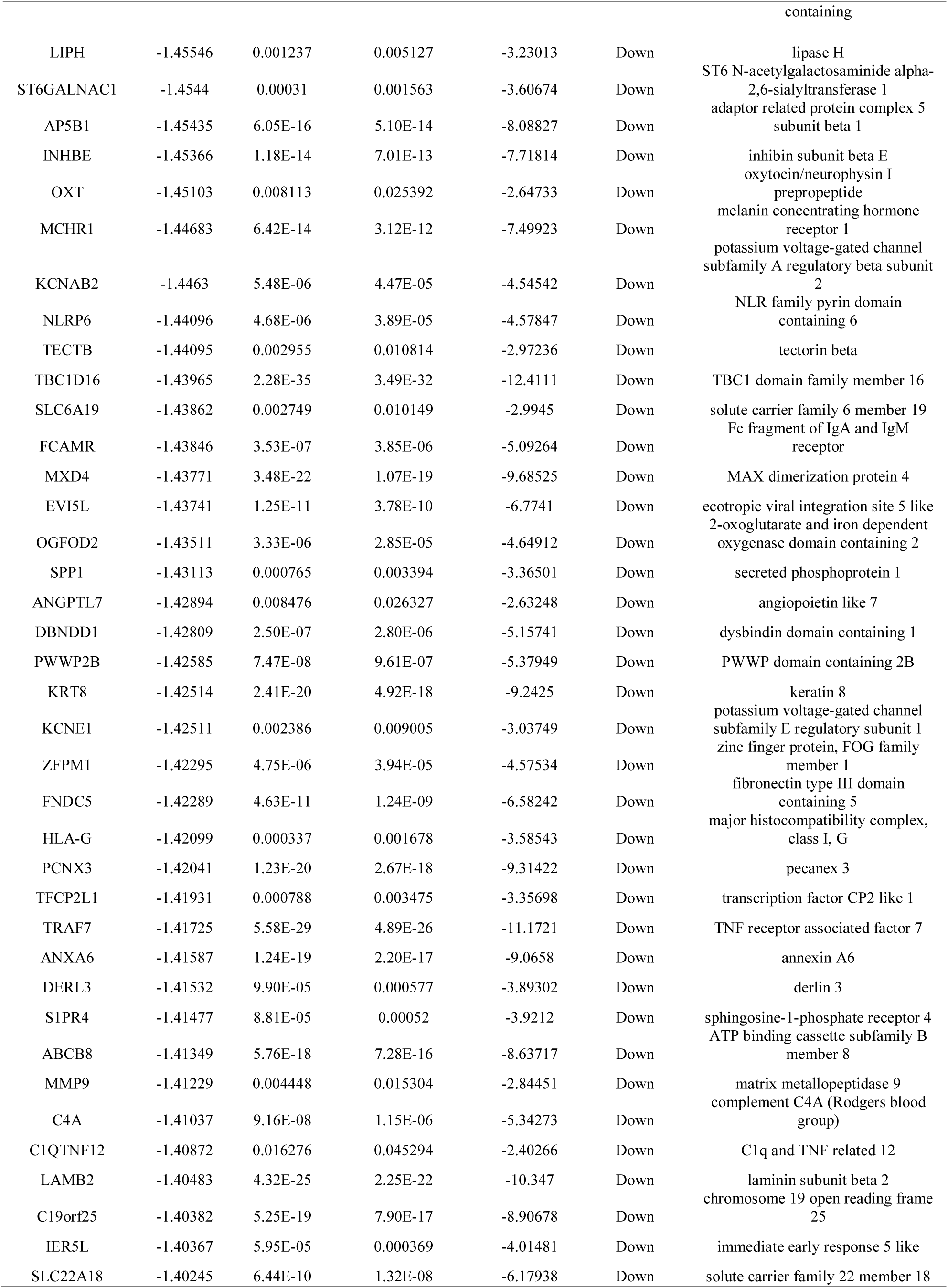
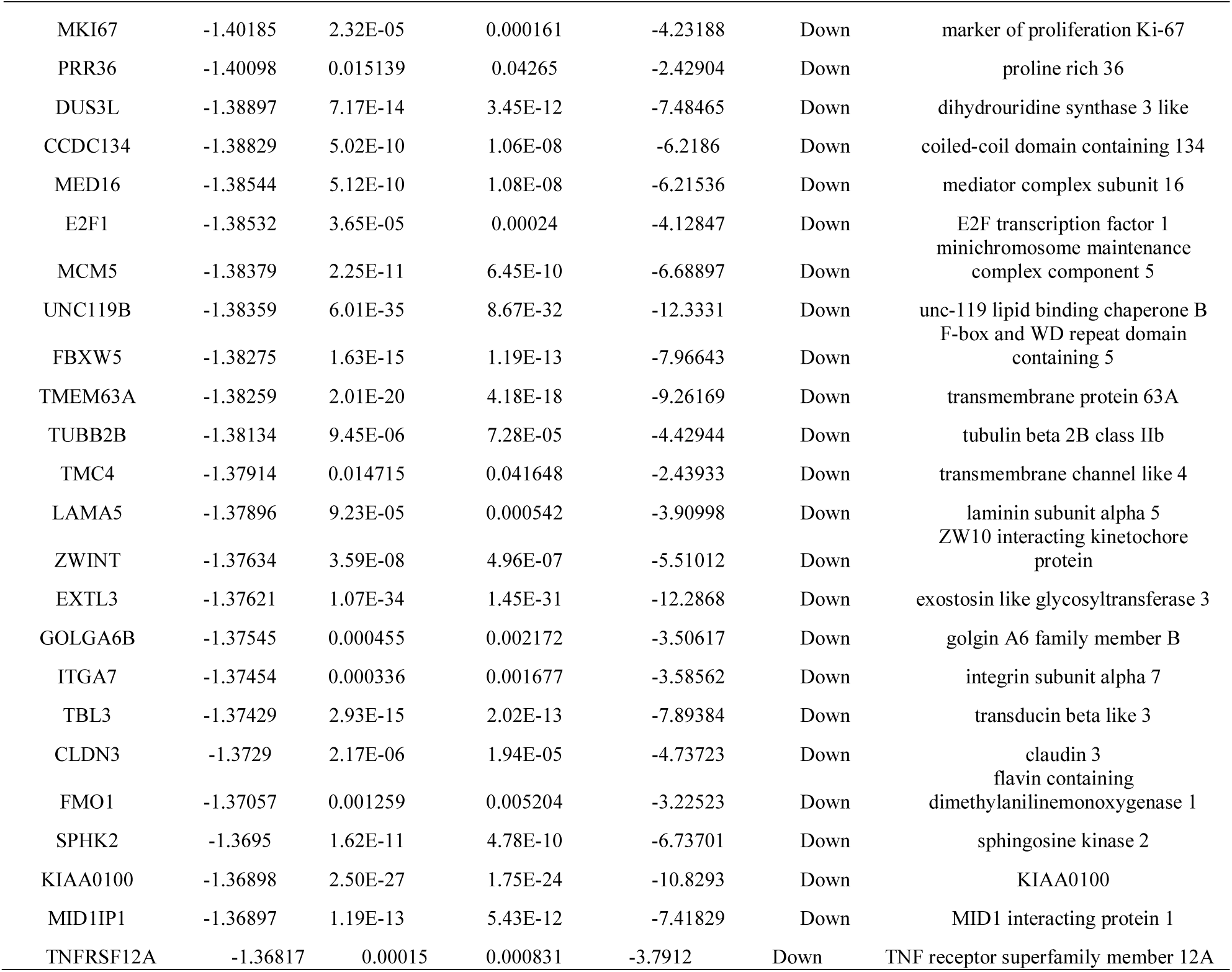
The statistical metrics for key differentially expressed genes (DEGs)

### GO and pathway enrichment analyses of DEGs

The online software g:Profiler was used to identify overrepresented GO categories and REACTOME pathways. The up regulated genes and all of the down regulated genes were uploaded to the gene list separately. GO enrichment analysis showed that the up regulated genes were significantly enriched in BP, including signaling and response to stimulus. The down regulated genes were also significantly enriched in BP, including regulation of biological quality and cell development. As for CC, the up regulated genes were enriched in chromatin and integral component of plasma membrane, and the down regulated genes were enriched in cell periphery and membrane. In addition, with regards to MF, the up regulated genes were enriched in DNA-binding transcription activator activity, RNA polymerase II-specific and signaling receptor binding, and the down regulated genes were enriched in calcium ion binding and structural molecule activity. The results are shown in full in Table 2. REACTOME pathway enrichment analysis found significantly enriched pathways. The up regulated genes were enriched in signaling by interleukins and signal transduction, while all of the down regulated genes were enriched in extracellular matrix organization and biological oxidations. The results are shown in full in Table 3.

**Table 2.**
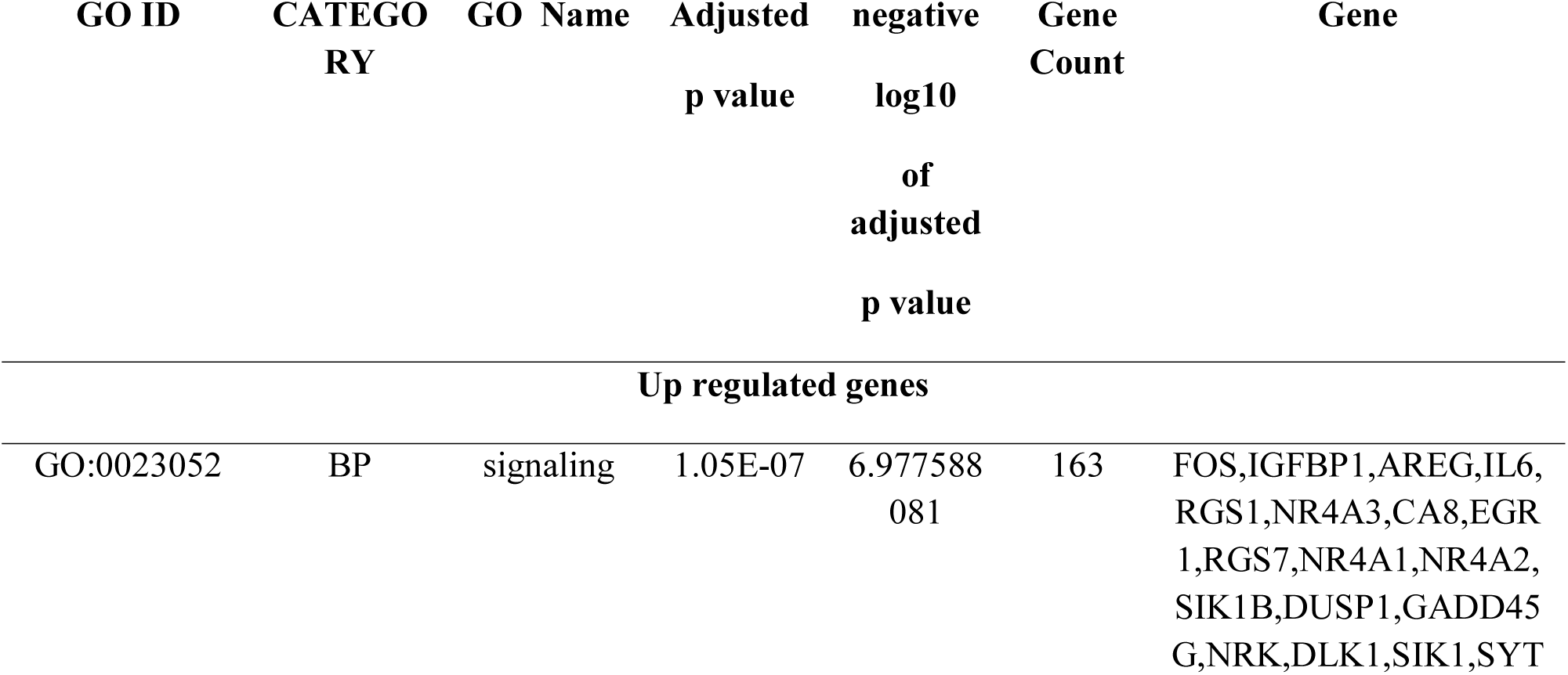
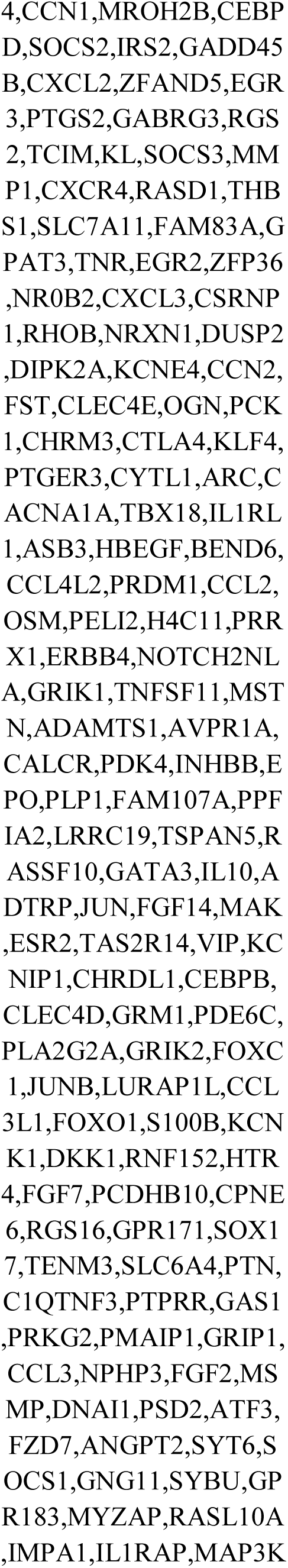
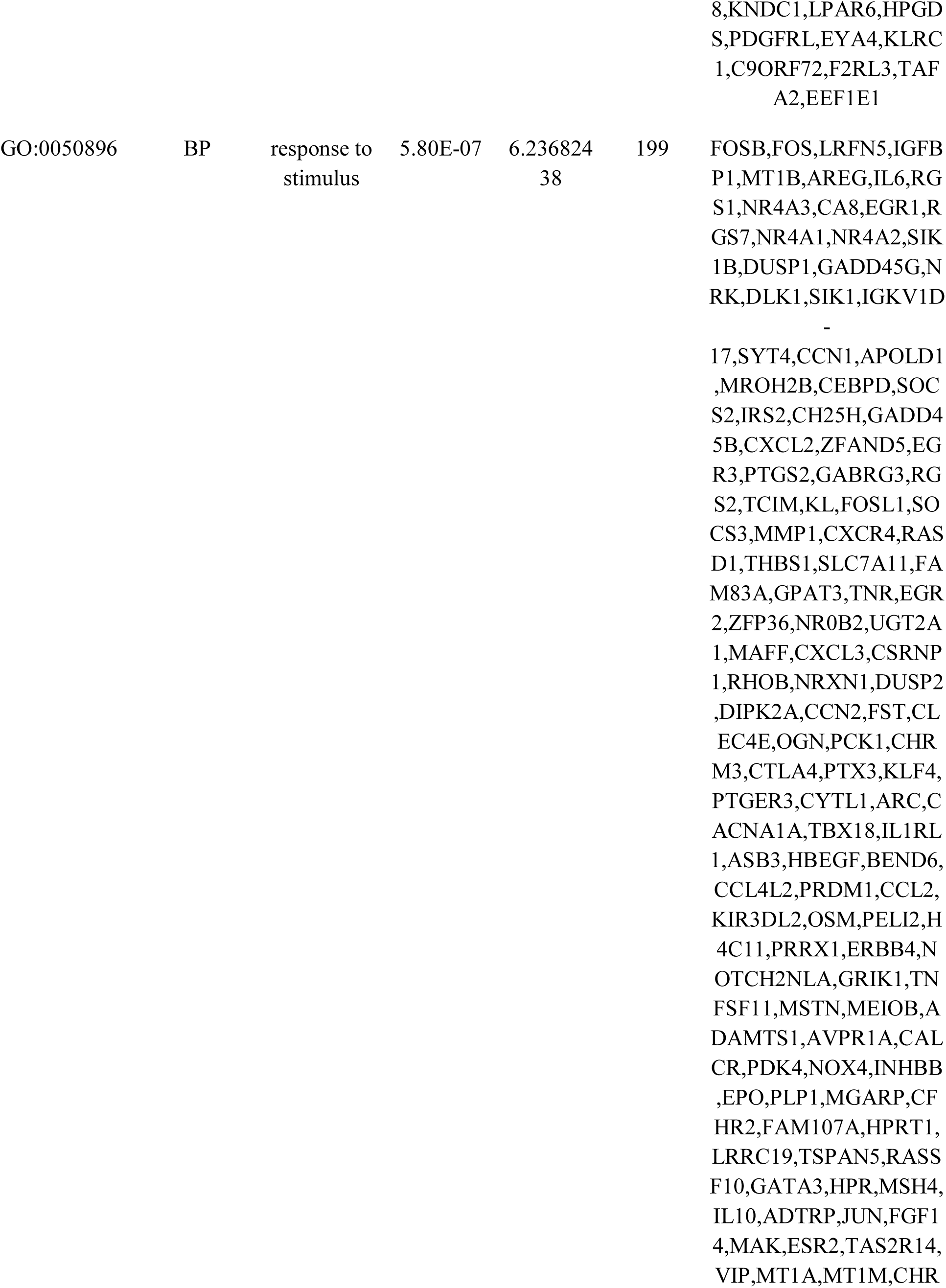
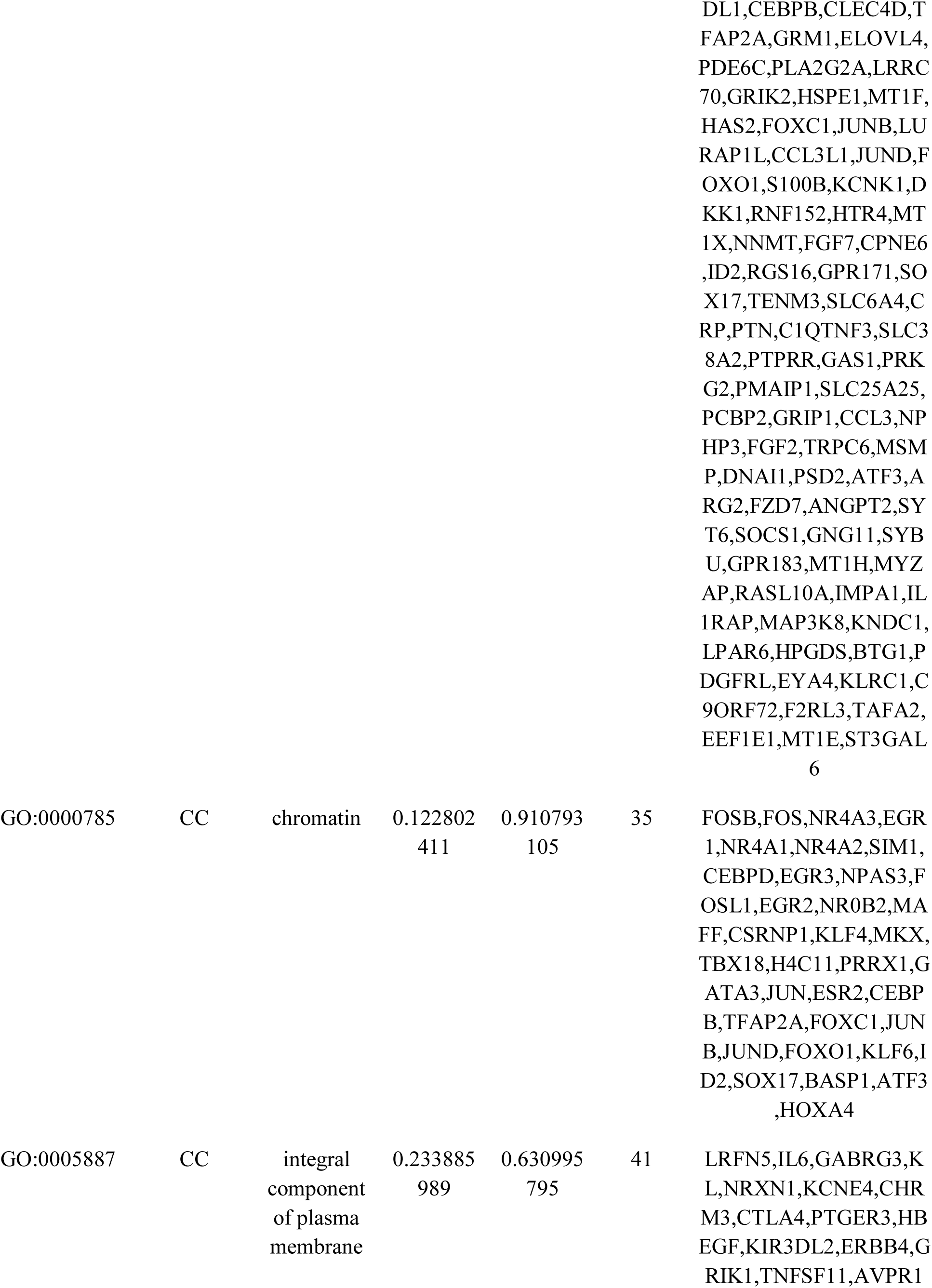
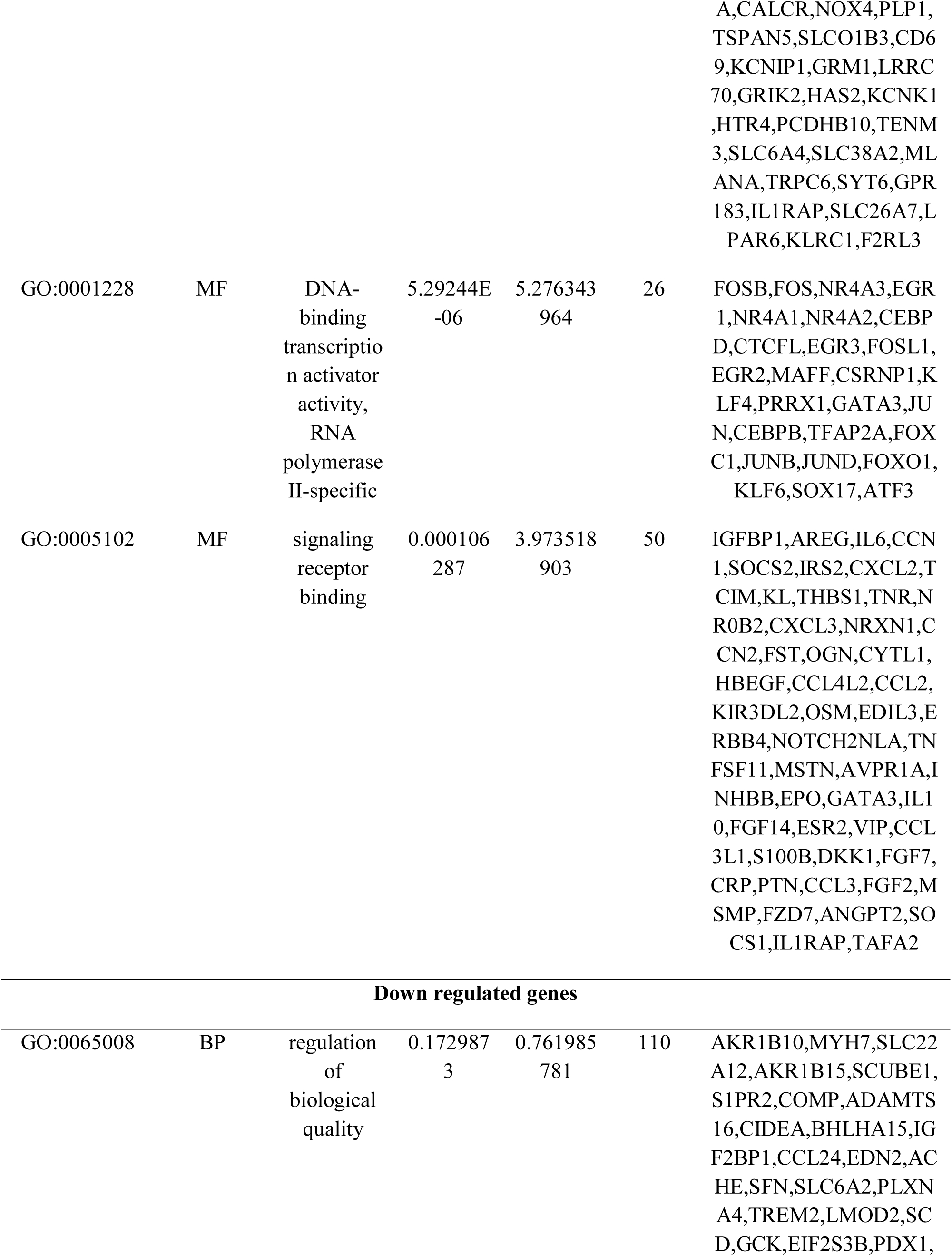
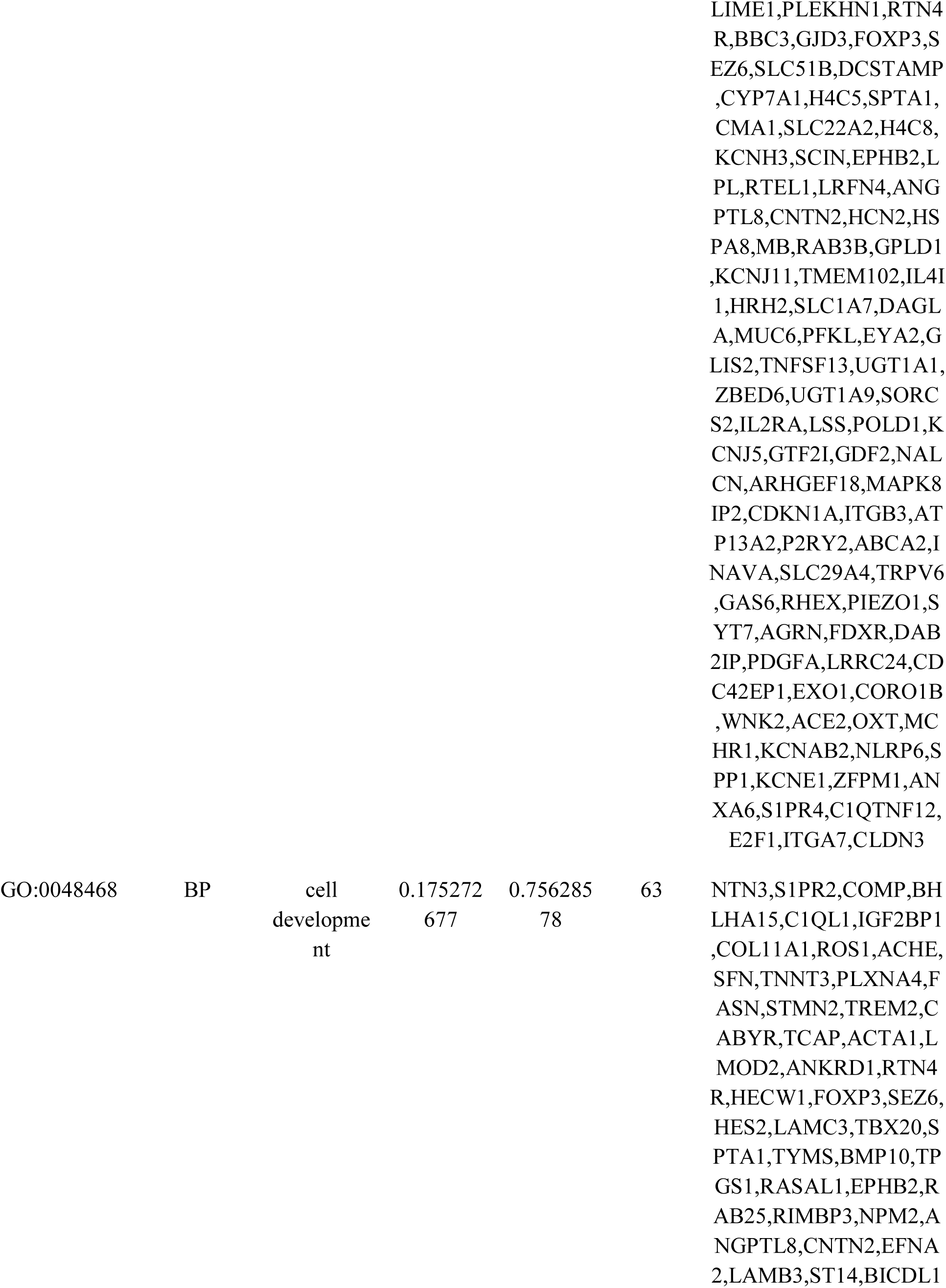
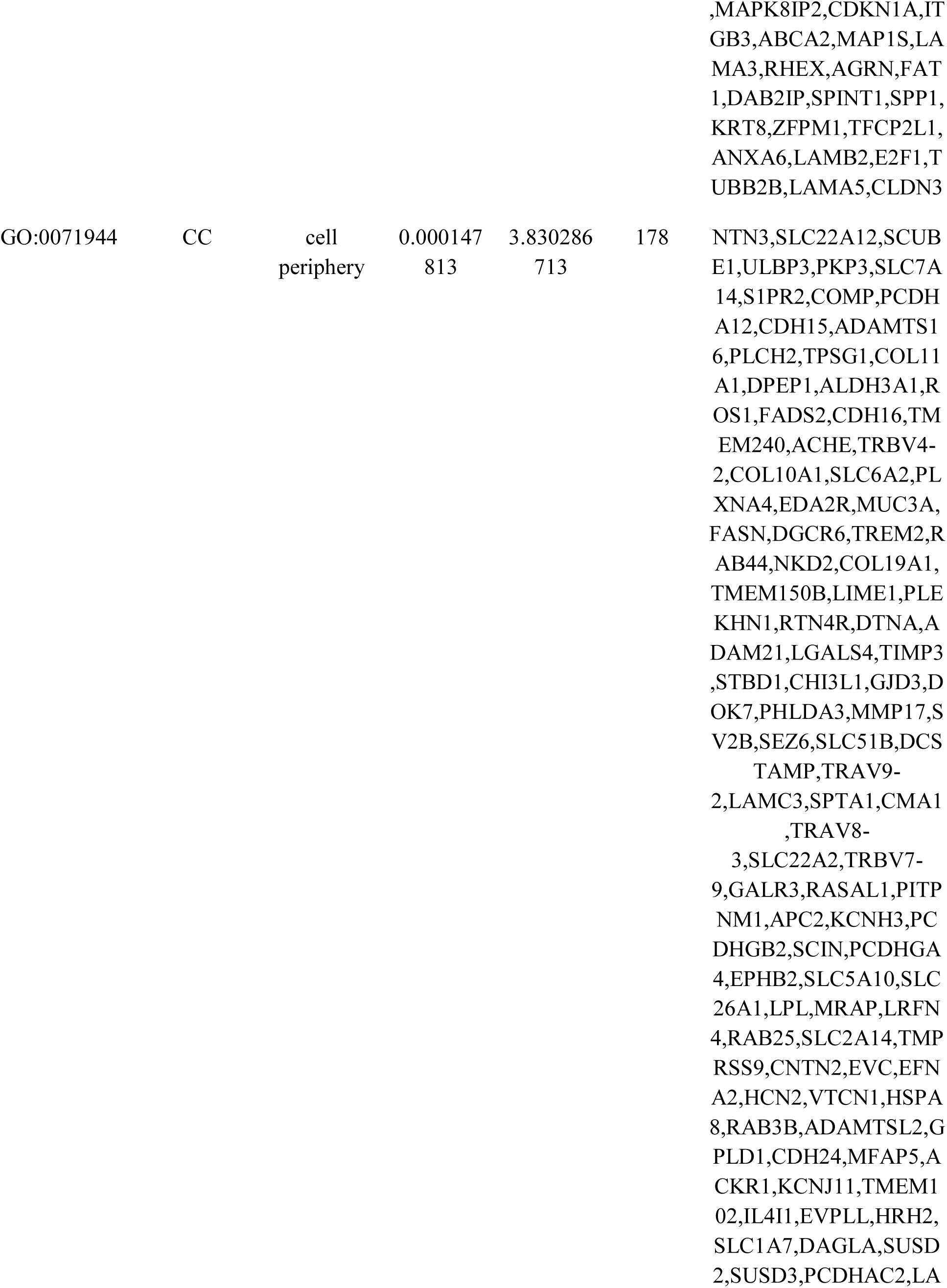
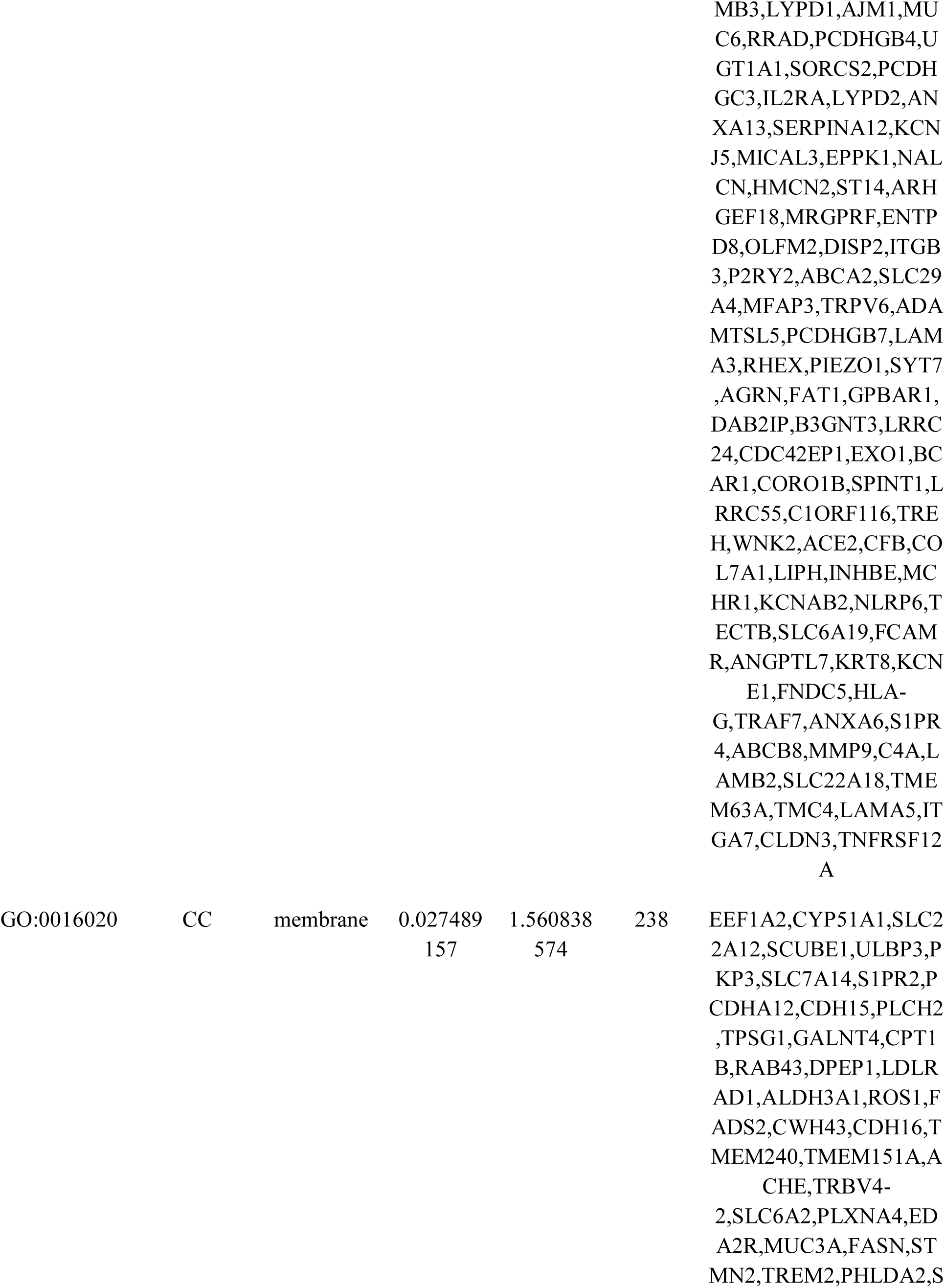
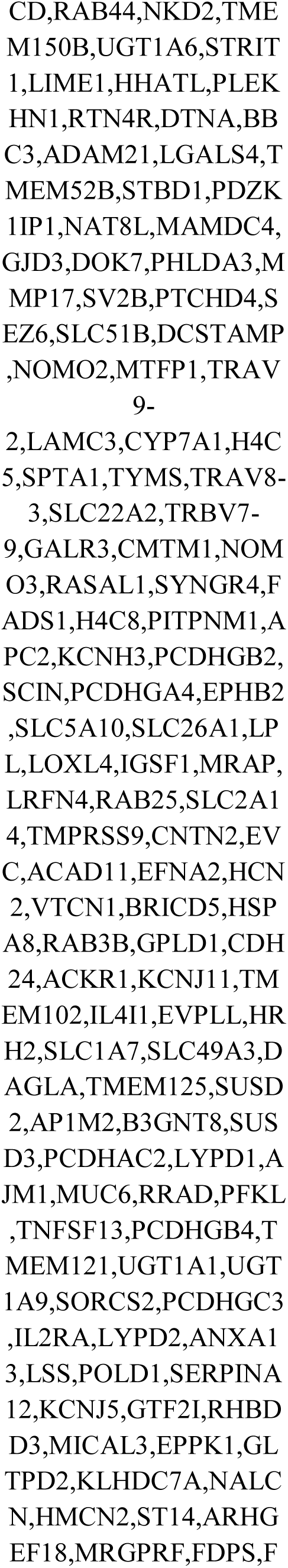
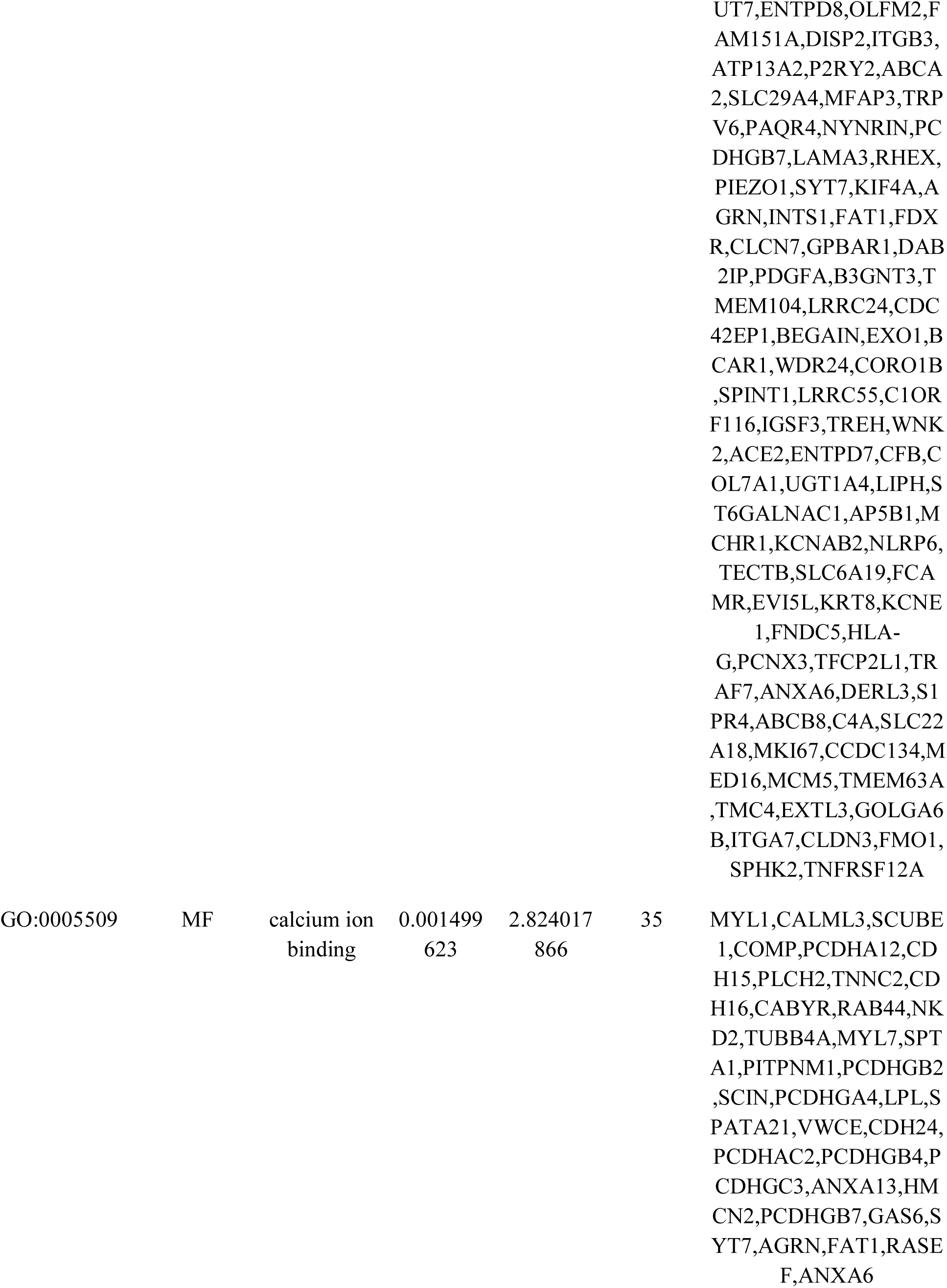
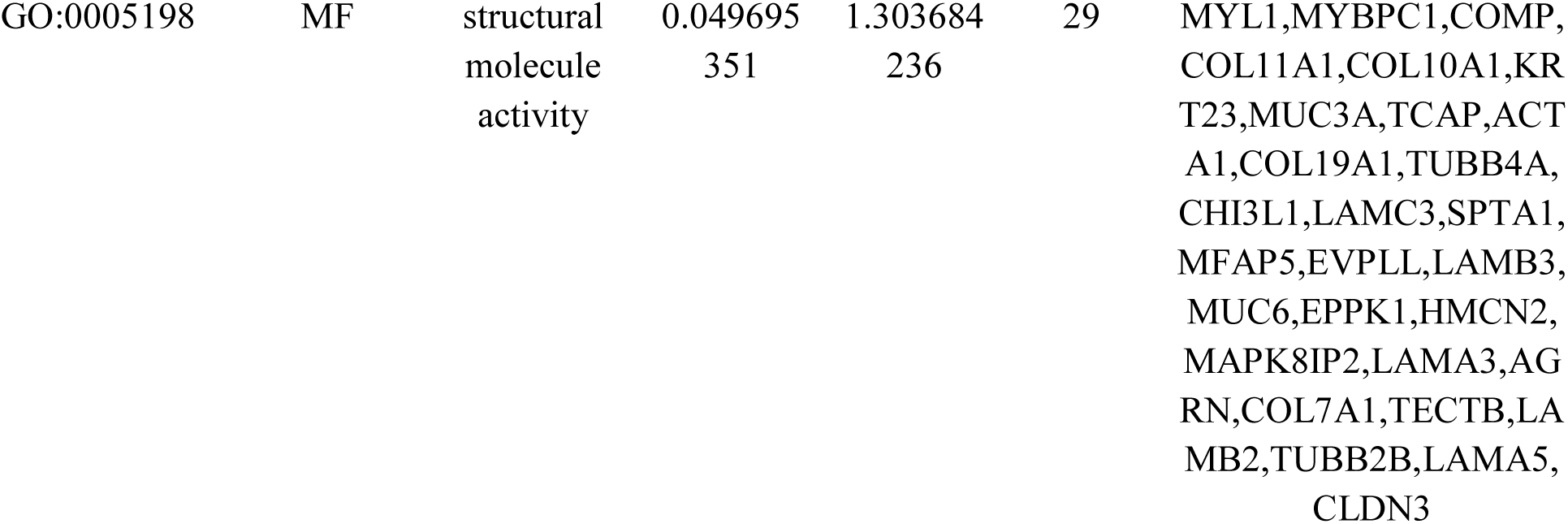
The enriched GO terms of the up and down regulated differentially expressed genes

**Table 3.**
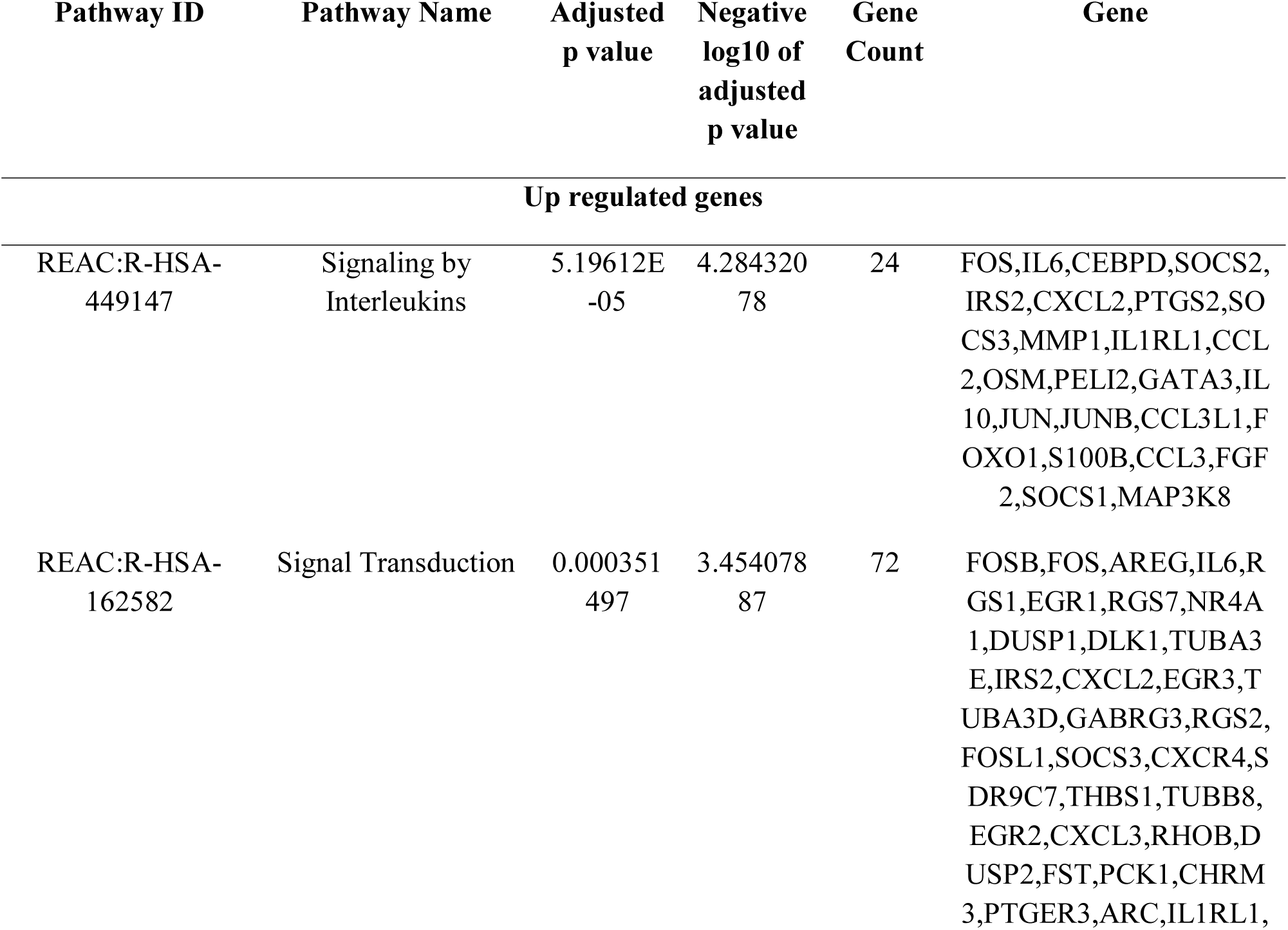
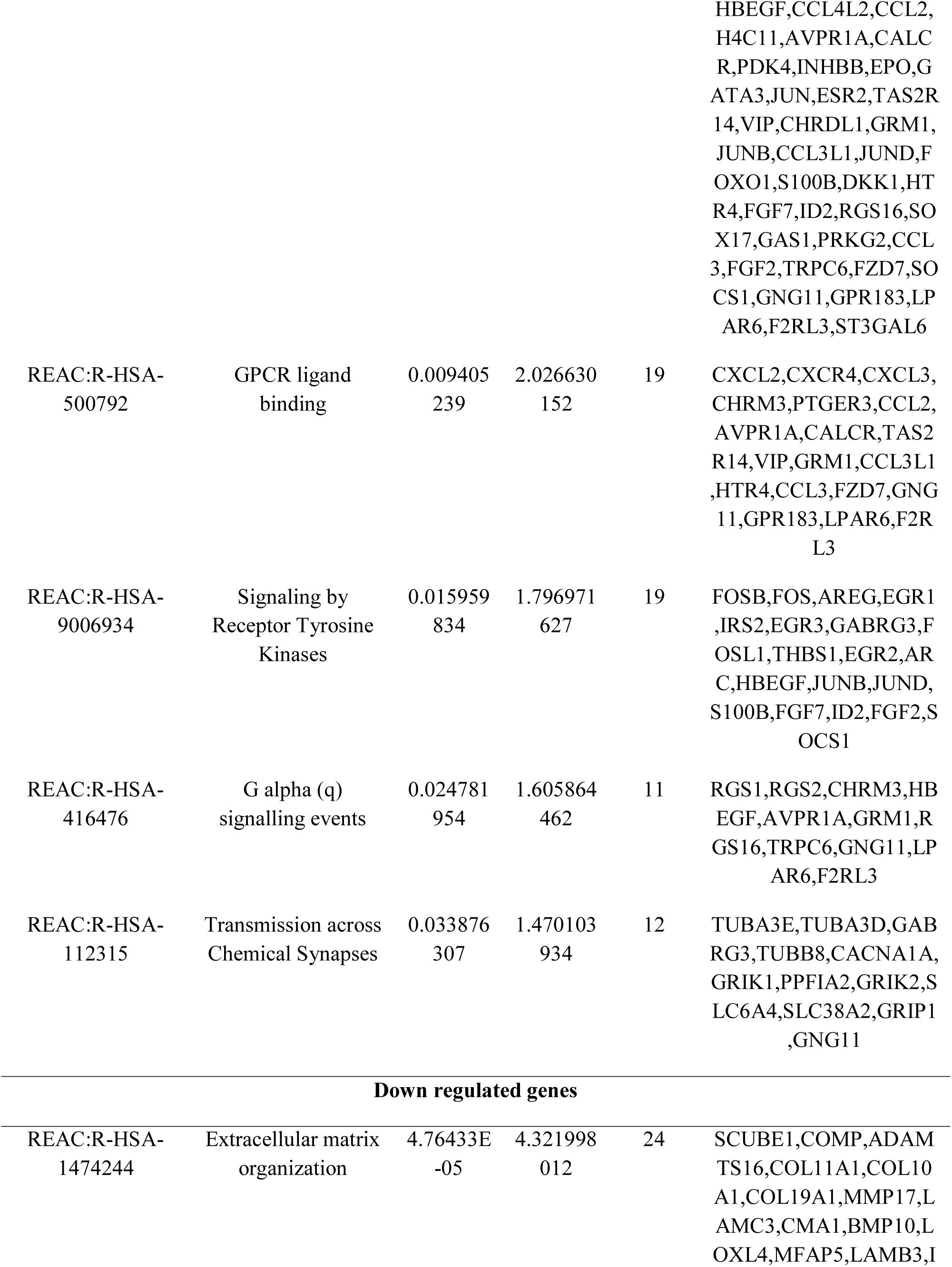
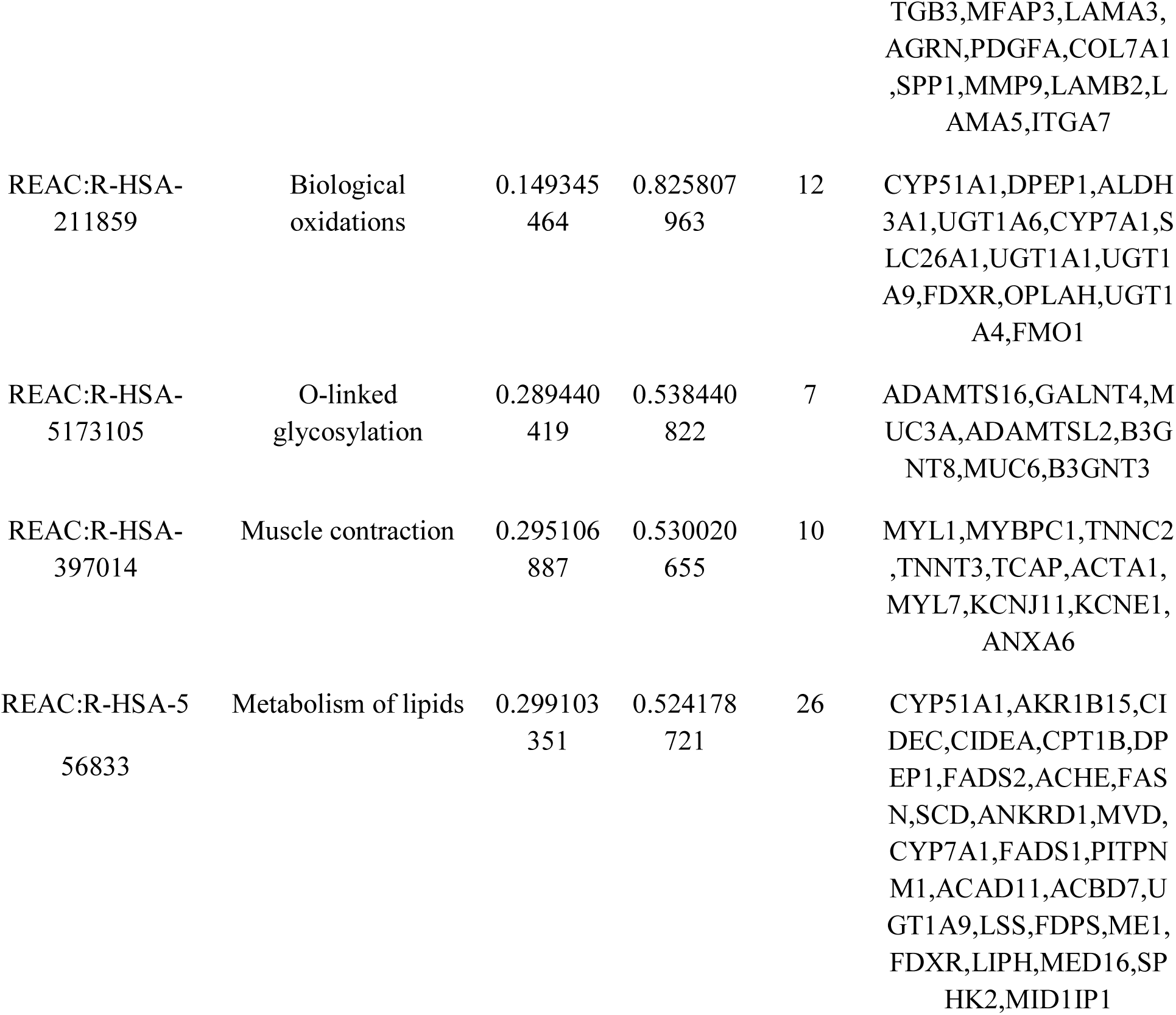
The enriched pathway terms of the up and down regulated differentially expressed genes

### Construction of the PPI network and module analysis

All of the up regulated genes and the down regulated genes were uploaded to Hippie interactome database to draw the PPI network. A total of 3199 nodes and 5019 edges were analyzed in Cytoscape and is displayed in Fig. 3. The hub genes with high node degree, betweenness centrality, stress centrality and closeness centrality were selected as the hub genes of NAFLD. These hub genes were ESR2, JUN, PTN, PTGER3, CEBPB, IKBKG, HSPA8, SFN, CDKN1A and E2F1. All of the results are shown in Tables 4. The two significant modules are displayed in Fig. 4A and Fig. 4B. There are 38 nodes and 121 edges genes involved in module 1 and 45 nodes, and 96 edges genes involved in module 2. These modules were significantly enriched in signaling by interleukins, signaling, signal transduction, response to stimulus, signaling by receptor tyrosine kinases, chromatin, DNA-binding transcription activator activity, RNA polymerase II-specific, metabolism of lipids and regulation of biological quality.

**Fig. 3.**
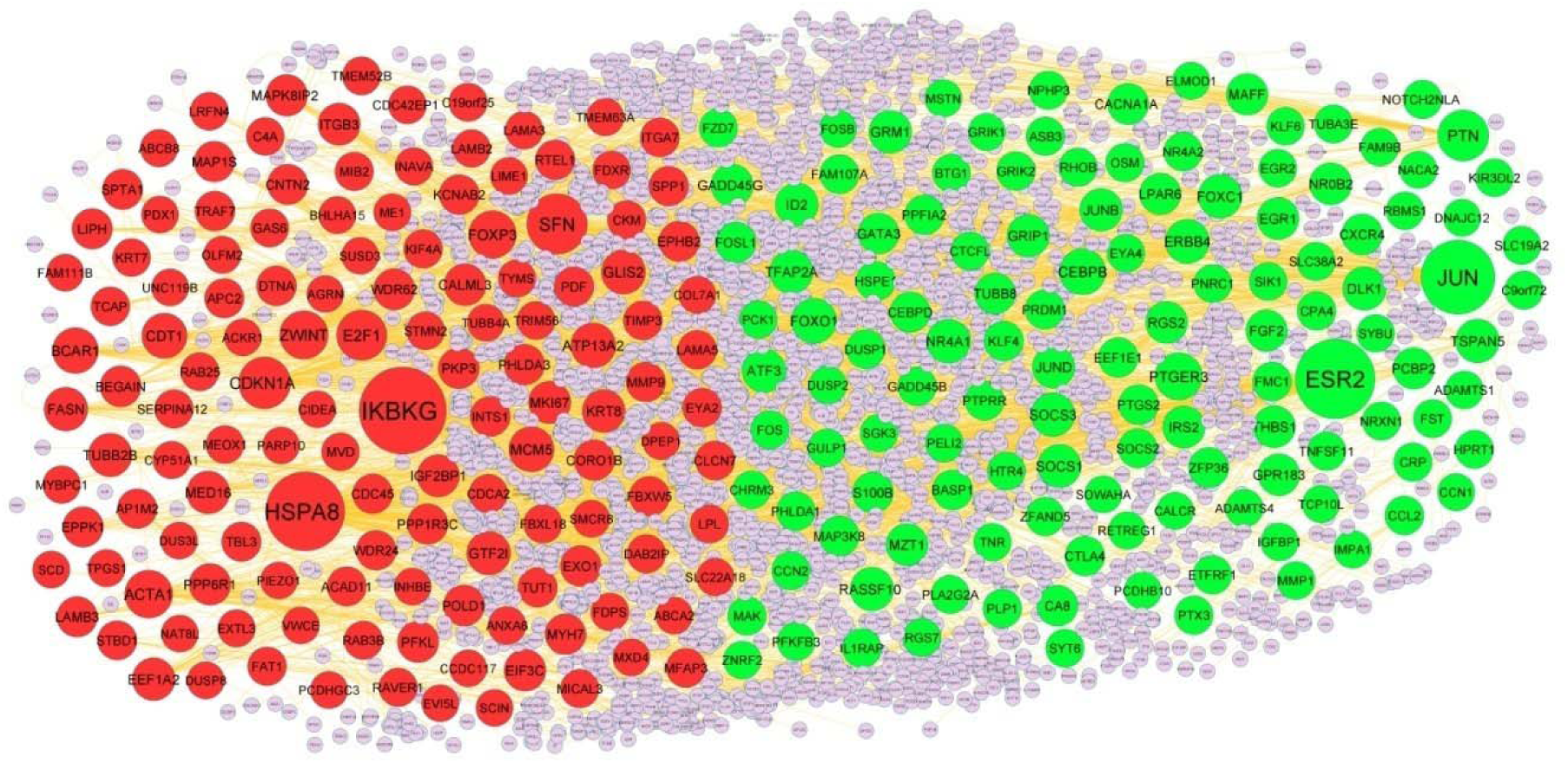
PPI network of DEGs. Up regulated genes are marked in green color; down regulated genes are marked in red color.

**Fig. 4.**
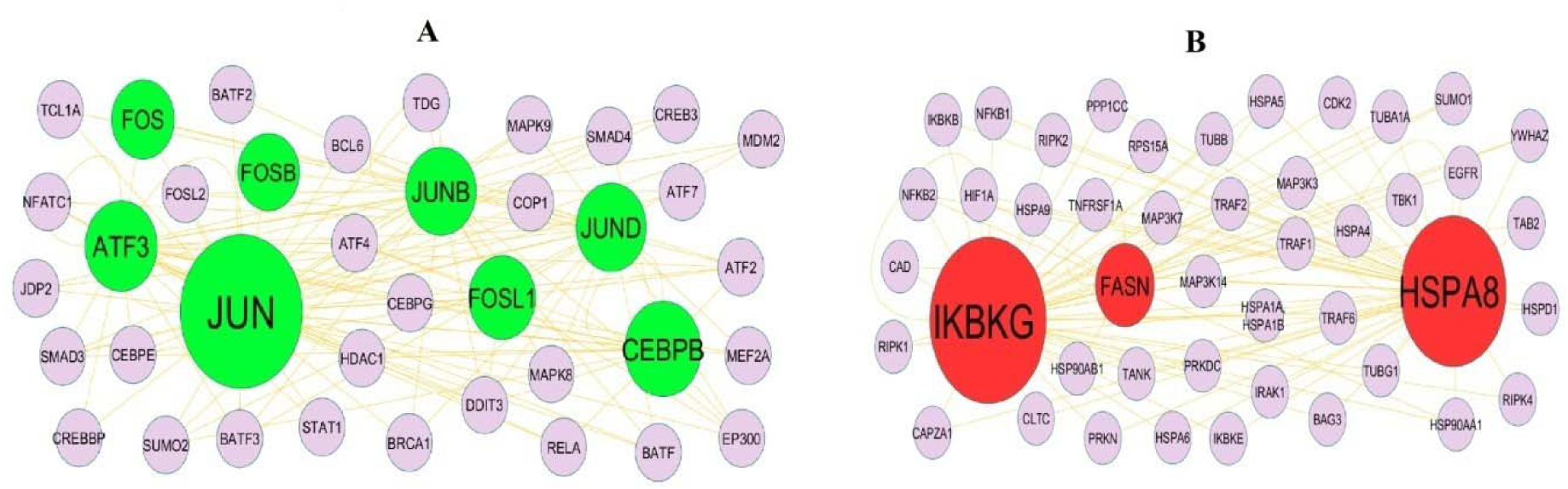
Modules of isolated form PPI of DEGs. (A) The most significant module was obtained from PPI network with 95 nodes and 103 edges for up regulated genes (B) The most significant module was obtained from PPI network with 79 nodes and 131 edges for down regulated genes. Up regulated genes are marked in green; down regulated genes are marked in red

**Table 4.** Topology table for up and down regulated genes.

| Regulation | Node | Degree | Betweenness | Stress | Closeness |
| --- | --- | --- | --- | --- | --- |
| Up | ESR2 | 252 | 0.153302 | 16307122 | 0.327825 |
| Up | JUN | 218 | 0.122226 | 20151740 | 0.318931 |
| Up | PTN | 83 | 0.04122 | 6599240 | 0.261213 |
| Up | PTGER3 | 61 | 0.04852 | 9842220 | 0.243274 |
| Up | CEBPB | 54 | 0.013264 | 2179706 | 0.289255 |
| Up | FOXO1 | 46 | 0.015398 | 1958960 | 0.286178 |
| Up | ERBB4 | 42 | 0.021503 | 4864528 | 0.266298 |
| Up | NR4A1 | 40 | 0.013258 | 2508616 | 0.271305 |
| Up | FOXC1 | 40 | 0.014365 | 2960242 | 0.256879 |
| Up | SOCS3 | 39 | 0.015854 | 1846382 | 0.268846 |
| Up | TSPAN5 | 39 | 0.021529 | 3536542 | 0.179423 |
| Up | ATF3 | 38 | 0.008644 | 888320 | 0.274996 |
| Up | DLK1 | 38 | 0.024667 | 2025172 | 0.198226 |
| Up | RASSF10 | 38 | 0.017409 | 2496598 | 0.256983 |
| Up | SOCS1 | 37 | 0.011257 | 1943118 | 0.266745 |
| Up | TFAP2A | 37 | 0.01778 | 1917734 | 0.284638 |
| Up | ID2 | 35 | 0.014538 | 1684376 | 0.259632 |
| Up | GRM1 | 33 | 0.008081 | 1405306 | 0.252868 |
| Up | JUNB | 33 | 0.003916 | 464160 | 0.27105 |
| Up | JUND | 32 | 0.003605 | 581468 | 0.278421 |
| Up | GRIP1 | 32 | 0.027221 | 4286096 | 0.255638 |
| Up | GADD45G | 31 | 0.018252 | 1295156 | 0.289018 |
| Up | IRS2 | 31 | 0.011813 | 1244018 | 0.280018 |
| Up | NR0B2 | 30 | 0.011956 | 1085608 | 0.273244 |
| Up | THBS1 | 30 | 0.020429 | 1220078 | 0.234295 |
| Up | EGR1 | 29 | 0.008524 | 1240050 | 0.268142 |
| Up | RGS2 | 29 | 0.013439 | 613038 | 0.268846 |
| Up | MZT1 | 28 | 0.01342 | 1489952 | 0.234054 |
| Up | GPR183 | 28 | 0.011006 | 4785362 | 0.187047 |
| Up | MAFF | 27 | 0.0076 | 914614 | 0.244436 |
| Up | PCBP2 | 26 | 0.007207 | 976748 | 0.251885 |
| Up | LPAR6 | 26 | 0.013687 | 1227722 | 0.21789 |
| Up | FGF2 | 25 | 0.018904 | 2947892 | 0.245305 |
| Up | CACNA1A | 24 | 0.014346 | 1032998 | 0.245419 |
| Up | CA8 | 23 | 0.011729 | 1626402 | 0.212977 |
| Up | CXCR4 | 23 | 0.011272 | 1056072 | 0.2569 |
| Up | S100B | 22 | 0.00813 | 1264412 | 0.248279 |
| Up | GATA3 | 22 | 0.005995 | 1401804 | 0.251685 |
| Up | TUBB8 | 22 | 0.010257 | 1276630 | 0.244926 |
| Up | FOSL1 | 19 | 0.004451 | 290374 | 0.270819 |
| Up | ZFP36 | 19 | 0.005886 | 855226 | 0.252586 |
| Up | MAP3K8 | 18 | 0.002239 | 664468 | 0.252687 |
| Up | CEBPD | 18 | 0.003277 | 452042 | 0.256009 |
| Up | BASP1 | 18 | 0.004385 | 972442 | 0.25077 |
| Up | PTGS2 | 17 | 0.00354 | 684102 | 0.250098 |
| Up | CTLA4 | 16 | 0.004535 | 561138 | 0.232408 |
| Up | EEF1E1 | 16 | 0.006062 | 993528 | 0.230033 |
| Up | PPFIA2 | 15 | 0.004649 | 430712 | 0.240218 |
| Up | HSPE1 | 14 | 0.005538 | 963234 | 0.251486 |
| Up | KLF4 | 14 | 0.002381 | 401354 | 0.249803 |
| Up | PRDM1 | 14 | 0.004147 | 663736 | 0.233674 |
| Up | PTPRR | 13 | 0.002529 | 397948 | 0.230167 |
| Up | SOCS2 | 12 | 2.25E-04 | 29864 | 0.225731 |
| Up | FAM107A | 12 | 0.004314 | 391420 | 0.23653 |
| Up | DUSP1 | 11 | 8.29E-04 | 200904 | 0.236671 |
| Up | IL1RAP | 11 | 0.005075 | 490034 | 0.226294 |
| Up | CCL2 | 11 | 0.004963 | 301058 | 0.179221 |
| Up | GRIK2 | 10 | 0.002374 | 159532 | 0.213235 |
| Up | PHLDA1 | 10 | 0.002297 | 380826 | 0.244135 |
| Up | ETFRF1 | 10 | 0.004407 | 230860 | 0.208651 |
| Up | CPA4 | 10 | 0.005088 | 610960 | 0.203854 |
| Up | FMC1 | 10 | 0.003795 | 509566 | 0.219229 |
| Up | TNFSF11 | 9 | 0.006207 | 499546 | 0.228066 |
| Up | GADD45B | 9 | 0.001894 | 221200 | 0.226326 |
| Up | PTX3 | 9 | 0.003429 | 288438 | 0.200316 |
| Up | PLP1 | 9 | 0.002263 | 205084 | 0.214083 |
| Up | ZFAND5 | 9 | 0.00214 | 106574 | 0.270013 |
| Up | NR4A2 | 9 | 0.00141 | 126664 | 0.222549 |
| Up | NOTCH2NLA | 9 | 0.002972 | 234194 | 0.220463 |
| Up | CRP | 8 | 0.002804 | 195986 | 0.192761 |
| Up | OSM | 8 | 0.003812 | 215368 | 0.210981 |
| Up | NPHP3 | 8 | 0.003428 | 308932 | 0.202993 |
| Up | C9orf72 | 8 | 0.004954 | 494952 | 0.210004 |
| Up | NRXN1 | 8 | 0.004407 | 299852 | 0.204906 |
| Up | ASB3 | 8 | 0.001938 | 714486 | 0.184868 |
| Up | MAK | 8 | 0.001972 | 159538 | 0.226601 |
| Up | RHOB | 8 | 0.003182 | 393652 | 0.197094 |
| Up | TUBA3E | 8 | 0.003786 | 289220 | 0.20426 |
| Up | SLC38A2 | 8 | 0.002598 | 229690 | 0.192015 |
| Up | FST | 7 | 0.003149 | 1000482 | 0.16387 |
| Up | HPRT1 | 7 | 0.002519 | 265684 | 0.190757 |
| Up | PLA2G2A | 7 | 0.001582 | 115638 | 0.227657 |
| Up | FZD7 | 7 | 0.001934 | 696944 | 0.182393 |
| Up | CCN2 | 7 | 0.008071 | 354698 | 0.266723 |
| Up | SLC19A2 | 7 | 0.003778 | 297990 | 0.165545 |
| Up | SYT6 | 7 | 0.002519 | 132836 | 0.208679 |
| Up | FOS | 6 | 0.001001 | 233150 | 0.246889 |
| Up | BTG1 | 6 | 0.001312 | 158508 | 0.220019 |
| Up | RGS7 | 6 | 0.001373 | 68622 | 0.193655 |
| Up | MMP1 | 6 | 0.002086 | 85636 | 0.162162 |
| Up | ADAMTS1 | 6 | 0.002369 | 102166 | 0.18102 |
| Up | CTCFL | 6 | 0.001763 | 246802 | 0.228066 |
| Up | PELI2 | 5 | 6.63E-04 | 105816 | 0.222643 |
| Up | KLF6 | 5 | 6.77E-04 | 75600 | 0.236073 |
| Up | EYA4 | 5 | 7.59E-04 | 91984 | 0.197794 |
| Up | KIR3DL2 | 5 | 1 | 6 | 1 |
| Up | PFKFB3 | 5 | 3.28E-05 | 6506 | 0.215391 |
| Up | PNRC1 | 5 | 0.00139 | 114074 | 0.222036 |
| Up | RBMS1 | 5 | 0.001488 | 192558 | 0.183883 |
| Up | SIK1 | 5 | 0.00146 | 115492 | 0.232988 |
| Up | HTR4 | 5 | 5.45E-05 | 12090 | 0.216095 |
| Up | EGR2 | 5 | 6.76E-04 | 100080 | 0.220401 |
| Up | FAM9B | 5 | 1 | 20 | 1 |
| Up | ZNRF2 | 4 | 7.56E-04 | 76784 | 0.23151 |
| Up | SGK3 | 4 | 0.001265 | 170160 | 0.224899 |
| Up | IGFBP1 | 4 | 0.00189 | 182178 | 0.15723 |
| Up | IMPA1 | 4 | 6.30E-04 | 43712 | 0.20232 |
| Up | SYBU | 4 | 3.36E-04 | 36546 | 0.209022 |
| Up | CHRM3 | 4 | 6.68E-04 | 81146 | 0.215508 |
| Up | GULP1 | 4 | 3.48E-04 | 43006 | 0.22665 |
| Up | ADAMTS4 | 4 | 0.001457 | 106596 | 0.193195 |
| Up | DUSP2 | 3 | 5.91E-06 | 2284 | 0.216389 |
| Up | TNR | 3 | 6.61E-04 | 34996 | 0.19117 |
| Up | RETREG1 | 3 | 9.48E-04 | 107682 | 0.176079 |
| Up | FOSB | 2 | 0 | 0 | 0.242753 |
| Up | NACA2 | 2 | 6.30E-04 | 71244 | 0.19482 |
| Up | PCK1 | 2 | 1.12E-04 | 24282 | 0.233795 |
| Up | TCP10L | 1 | 0 | 0 | 1 |
| Up | MSTN | 1 | 0 | 0 | 0.213838 |
| Up | CALCR | 1 | 0 | 0 | 0.189833 |
| Up | GRIK1 | 1 | 0 | 0 | 0.203605 |
| Up | ELMOD1 | 1 | 0 | 0 | 0.217412 |
| Up | DNAJC12 | 1 | 0 | 0 | 0.175398 |
| Up | SOWAHA | 1 | 0 | 0 | 0.19951 |
| Up | CCN1 | 1 | 0 | 0 | 0.199209 |
| Up | PCDHB10 | 1 | 0 | 0 | 0.21973 |
| Down | IKBKG | 297 | 0.219998 | 28468976 | 0.339538 |
| Down | HSPA8 | 249 | 0.216486 | 22912020 | 0.352393 |
| Down | SFN | 139 | 0.096144 | 11786478 | 0.307827 |
| Down | CDKN1A | 89 | 0.044712 | 9732644 | 0.290314 |
| Down | E2F1 | 79 | 0.027908 | 5678252 | 0.277108 |
| Down | ZWINT | 68 | 0.034002 | 4334154 | 0.248746 |
| Down | ACTA1 | 65 | 0.031398 | 5160900 | 0.270358 |
| Down | FOXP3 | 63 | 0.026408 | 4439834 | 0.261644 |
| Down | BCAR1 | 53 | 0.026528 | 2498042 | 0.283595 |
| Down | MCM5 | 49 | 0.015828 | 3100420 | 0.270151 |
| Down | CDT1 | 49 | 0.0142 | 2142496 | 0.27268 |
| Down | ATP13A2 | 48 | 0.040851 | 1858586 | 0.281583 |
| Down | GLIS2 | 43 | 0.012738 | 2971676 | 0.267803 |
| Down | GTF2I | 40 | 0.013965 | 3548098 | 0.267352 |
| Down | FASN | 40 | 0.013278 | 1923754 | 0.294052 |
| Down | TUBB2B | 39 | 0.014139 | 3409940 | 0.261536 |
| Down | EEF1A2 | 35 | 0.011846 | 1154354 | 0.283722 |
| Down | MED16 | 35 | 0.010267 | 546374 | 0.274829 |
| Down | KRT8 | 35 | 0.022582 | 2216918 | 0.295889 |
| Down | ITGB3 | 32 | 0.017613 | 1787526 | 0.25309 |
| Down | MKI67 | 32 | 0.012591 | 2306860 | 0.26999 |
| Down | IGF2BP1 | 32 | 0.011046 | 1817704 | 0.264964 |
| Down | CALML3 | 30 | 0.016001 | 3689680 | 0.234624 |
| Down | EIF3C | 29 | 0.013833 | 1898480 | 0.255309 |
| Down | MAPK8IP2 | 28 | 0.014621 | 2092782 | 0.252888 |
| Down | PPP6R1 | 28 | 0.014191 | 2543882 | 0.236742 |
| Down | PFKL | 26 | 0.011293 | 2060970 | 0.262422 |
| Down | POLD1 | 25 | 0.01216 | 1380436 | 0.254103 |
| Down | WDR62 | 25 | 0.009598 | 1556774 | 0.257776 |
| Down | BEGAIN | 25 | 0.008918 | 621044 | 0.231358 |
| Down | EXO1 | 24 | 0.009378 | 611262 | 0.26679 |
| Down | PKP3 | 24 | 0.013902 | 706668 | 0.277788 |
| Down | CORO1B | 23 | 0.008296 | 1392340 | 0.240911 |
| Down | MYH7 | 23 | 0.010919 | 466678 | 0.228214 |
| Down | PPP1R3C | 22 | 0.009067 | 1597620 | 0.236231 |
| Down | MAP1S | 22 | 0.012693 | 971324 | 0.247177 |
| Down | PHLDA3 | 22 | 0.008565 | 1300372 | 0.238826 |
| Down | EPHB2 | 21 | 0.007088 | 1027538 | 0.246658 |
| Down | CNTN2 | 21 | 0.008116 | 442262 | 0.277885 |
| Down | EPPK1 | 21 | 0.006527 | 714440 | 0.285972 |
| Down | LAMB3 | 21 | 0.011464 | 1023764 | 0.243929 |
| Down | LIPH | 21 | 0.010728 | 701480 | 0.212692 |
| Down | SPTA1 | 20 | 0.006833 | 602872 | 0.253717 |
| Down | DTNA | 18 | 0.011282 | 1247794 | 0.206439 |
| Down | FBXW5 | 18 | 0.005902 | 970060 | 0.233657 |
| Down | WDR24 | 18 | 0.007909 | 812806 | 0.222925 |
| Down | KCNAB2 | 18 | 0.006806 | 539108 | 0.227821 |
| Down | INTS1 | 18 | 0.011943 | 1432184 | 0.240473 |
| Down | TRAF7 | 17 | 0.006461 | 891258 | 0.235268 |
| Down | CDC45 | 17 | 0.005438 | 340314 | 0.255083 |
| Down | AP1M2 | 16 | 0.009031 | 859896 | 0.213953 |
| Down | MMP9 | 16 | 0.010164 | 605996 | 0.202217 |
| Down | SPP1 | 16 | 0.007637 | 828418 | 0.229087 |
| Down | RTEL1 | 16 | 0.008419 | 692556 | 0.241976 |
| Down | APC2 | 15 | 0.003969 | 540810 | 0.238342 |
| Down | TBL3 | 15 | 0.005688 | 802840 | 0.241939 |
| Down | TUT1 | 15 | 0.005993 | 741238 | 0.224676 |
| Down | ACAD11 | 15 | 0.007604 | 390702 | 0.21475 |
| Down | PDF | 15 | 0.005927 | 815818 | 0.211967 |
| Down | STBD1 | 14 | 0.008739 | 588312 | 0.219411 |
| Down | MFAP3 | 14 | 0.008403 | 592372 | 0.208063 |
| Down | CLCN7 | 14 | 0.00565 | 345496 | 0.206506 |
| Down | ITGA7 | 13 | 0.004528 | 295182 | 0.203305 |
| Down | EYA2 | 12 | 0.005337 | 373532 | 0.226294 |
| Down | TCAP | 12 | 0.005861 | 210724 | 0.271979 |
| Down | LAMA5 | 12 | 0.004464 | 378806 | 0.203788 |
| Down | TIMP3 | 12 | 0.005346 | 371996 | 0.169851 |
| Down | KIF4A | 11 | 0.004493 | 663298 | 0.235007 |
| Down | FAT1 | 11 | 0.004601 | 383862 | 0.219821 |
| Down | RAB3B | 10 | 0.005664 | 629802 | 0.166921 |
| Down | DAB2IP | 10 | 0.001509 | 308434 | 0.247196 |
| Down | CKM | 10 | 0.003777 | 334172 | 0.239873 |
| Down | MICAL3 | 10 | 0.001711 | 278900 | 0.225747 |
| Down | MIB2 | 10 | 0.002635 | 175896 | 0.23745 |
| Down | SERPINA12 | 10 | 0.003286 | 1359806 | 0.162038 |
| Down | FBXL18 | 10 | 0.002126 | 410424 | 0.215728 |
| Down | RAVER1 | 9 | 0.002356 | 184440 | 0.232255 |
| Down | LPL | 9 | 0.002077 | 256246 | 0.201844 |
| Down | C19orf25 | 9 | 0.005017 | 454386 | 0.212322 |
| Down | CDCA2 | 9 | 0.002925 | 429662 | 0.232937 |
| Down | RAB25 | 9 | 0.003803 | 599076 | 0.208569 |
| Down | GAS6 | 8 | 0.004121 | 284918 | 0.204339 |
| Down | TMEM52B | 8 | 0.00642 | 1133566 | 0.226536 |
| Down | FDPS | 7 | 0.001512 | 77870 | 0.268983 |
| Down | STMN2 | 7 | 0.00198 | 112084 | 0.21843 |
| Down | LRFN4 | 7 | 7.76E-04 | 119920 | 0.2404 |
| Down | MEOX1 | 7 | 0.003778 | 227130 | 0.152699 |
| Down | TRIM56 | 7 | 0.00154 | 235690 | 0.240546 |
| Down | CDC42EP1 | 7 | 0.00145 | 147522 | 0.240382 |
| Down | ANXA6 | 6 | 8.32E-04 | 55866 | 0.210477 |
| Down | FDXR | 6 | 0.00196 | 213120 | 0.185019 |
| Down | C4A | 6 | 0.003488 | 217398 | 0.188659 |
| Down | LAMA3 | 6 | 0.002394 | 218314 | 0.225346 |
| Down | KRT7 | 6 | 0.001611 | 172556 | 0.239493 |
| Down | MVD | 6 | 0.002525 | 163688 | 0.227445 |
| Down | MYBPC1 | 5 | 0.001022 | 80464 | 0.199799 |
| Down | INAVA | 5 | 0.001535 | 221534 | 0.195974 |
| Down | LAMB2 | 5 | 0.001901 | 130904 | 0.193643 |
| Down | TYMS | 4 | 6.30E-04 | 48926 | 0.213594 |
| Down | SLC22A18 | 4 | 0.001272 | 66132 | 0.188749 |
| Down | CYP51A1 | 4 | 0.00189 | 204342 | 0.14622 |
| Down | DPEP1 | 3 | 0 | 0 | 1 |
| Down | ABCB8 | 3 | 0.00126 | 110762 | 0.178958 |
| Down | TPGS1 | 3 | 0.00126 | 231982 | 0.176275 |
| Down | TMEM63A | 3 | 6.52E-04 | 38766 | 0.160157 |
| Down | OLFM2 | 3 | 1 | 6 | 1 |
| Down | EXTL3 | 3 | 1 | 6 | 1 |
| Down | DUSP8 | 2 | 1.55E-06 | 342 | 0.219169 |
| Down | COL7A1 | 2 | 2.58E-04 | 22290 | 0.200038 |
| Down | BHLHA15 | 2 | 1.00E-07 | 2 | 0.170948 |
| Down | TUBB4A | 2 | 2.31E-04 | 62654 | 0.256443 |
| Down | ME1 | 2 | 3.08E-05 | 8426 | 0.195829 |
| Down | PCDHGC3 | 2 | 6.30E-04 | 51874 | 0.15275 |
| Down | SCD | 2 | 6.30E-04 | 30588 | 0.206816 |
| Down | SMCR8 | 1 | 0 | 0 | 0.173566 |
| Down | MXD4 | 1 | 0 | 0 | 1 |
| Down | UNC119B | 1 | 0 | 0 | 0.168749 |
| Down | ACKR1 | 1 | 0 | 0 | 0.15199 |
| Down | INHBE | 1 | 0 | 0 | 0.140804 |
| Down | SCIN | 1 | 0 | 0 | 0.212834 |
| Down | DUS3L | 1 | 0 | 0 | 0.194772 |
| Down | LIME1 | 1 | 0 | 0 | 0.19047 |
| Down | NAT8L | 1 | 0 | 0 | 0.152135 |
| Down | SUSD3 | 1 | 0 | 0 | 0.190049 |
| Down | CCDC117 | 1 | 0 | 0 | 0.260591 |
| Down | PIEZO1 | 1 | 0 | 0 | 0.152135 |
| Down | CIDEA | 1 | 0 | 0 | 0.139449 |
| Down | EVI5L | 1 | 0 | 0 | 0.195684 |
| Down | FAM111B | 1 | 0 | 0 | 0.21973 |
| Down | ABCA2 | 1 | 0 | 0 | 0.196108 |
| Down | AGRN | 1 | 0 | 0 | 0.197069 |
| Down | PDX1 | 1 | 0 | 0 | 0.215698 |
| Down | PARP10 | 1 | 0 | 0 | 0.253494 |
| Down | VWCE | 1 | 0 | 0 | 1 |

### MiRNA-hub gene regulatory network construction

To investigate the molecular mechanisms underlying the hub genes and miRNAs were searched by bioinformatics methods. The miRNAs of hub genes were predicted by miRNet database. The miRNA-hub gene regulatory network is consisted of 2546 nodes and 13598 edges, including 2204 miRNAs and 252 hub genes (Fig. 5). As shown in Table 5, miRNAs of hub genes were displayed. JUN might be the target of 118 miRNAs (ex; hsa-mir-199a-5p), FOXO1 might be the target of 63 miRNAs (ex; hsa-mir-33a-3p), FOXC1 might be the target of 62 miRNAs (ex; hsa-mir-561-3p), SOCS3 might be the target of 48 miRNAs (ex; hsa-mir-22-5p), CEBPB might be the target of 47 miRNAs (ex; hsa-mir-3613-3p), CDKN1A might be the target of 187 miRNAs (ex; hsa-mir-30b-3p), HSPA8 might be the target of 116 miRNAs (ex; hsa-mir-338-5p), FASN might be the target of 116 miRNAs (ex; hsa-mir-103a-3p), E2F1 might be the target of 74 miRNAs (ex; hsa-mir-150-5p) and CDT1 might be the target of 45 miRNAs (ex; hsa-mir-202-3p).

**Fig. 5.**
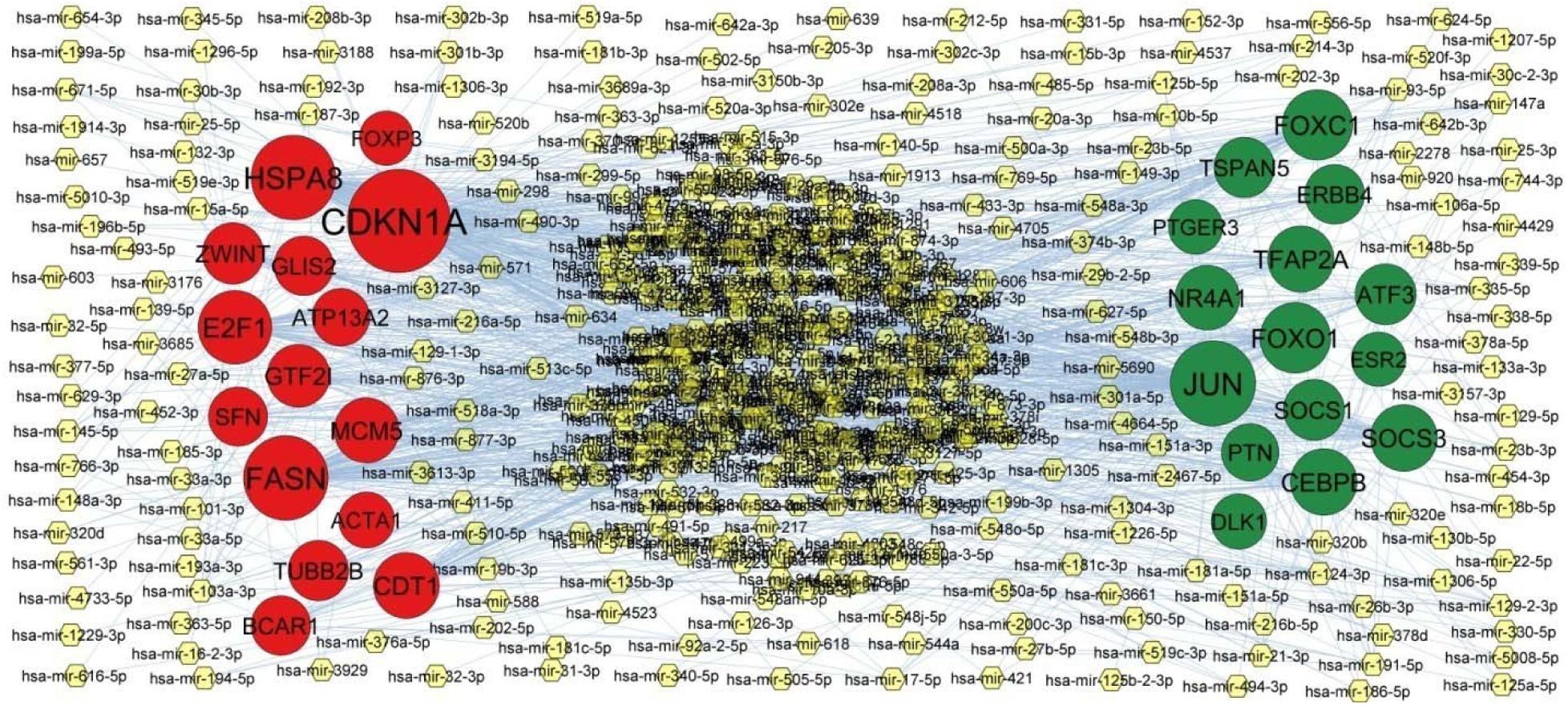
Target gene - miRNA regulatory network between target genes. The yellow color diamond nodes represent the key miRNAs; up regulated genes are marked in green color; down regulated genes are marked in red color.

**Table 5.** miRNA - target gene and TF - target gene interaction

| Regulation | Target Genes | Degree | MicroRNA | Regulation | Target Genes | Degree | TF |
| --- | --- | --- | --- | --- | --- | --- | --- |
| Up | JUN | 118 | hsa-mir-199a-5p | Up | JUN | 61 | GATA2 |
| Up | FOXO1 | 63 | hsa-mir-33a-3p | Up | SOCS3 | 56 | RUNX1 |
| Up | FOXC1 | 62 | hsa-mir-561-3p | Up | TFAP2A | 55 | SUZ12 |
| Up | SOCS3 | 48 | hsa-mir-22-5p | Up | ATF3 | 48 | NUCKS1 |
| Up | CEBPB | 47 | hsa-mir-3613-3p | Up | CEBPB | 46 | FOXP2 |
| Up | TFAP2A | 46 | hsa-mir-1306-3p | Up | FOXO1 | 43 | PAX3 |
| Up | NR4A1 | 40 | hsa-mir-125a-5p | Up | SOCS1 | 39 | STAT5A |
| Up | SOCS1 | 26 | hsa-mir-21-3p | Up | FOXC1 | 37 | SALL4 |
| Up | ATF3 | 26 | hsa-mir-147a | Up | NR4A1 | 36 | MYC |
| Up | TSPAN5 | 24 | hsa-mir-181a-5p | Up | PTGER3 | 33 | TFCP2L1 |
| Up | ERBB4 | 21 | hsa-mir-433-3p | Up | TSPAN5 | 30 | ASH2L |
| Up | PTN | 12 | hsa-mir-376a-5p | Up | ERBB4 | 29 | MEF2A |
| Up | DLK1 | 5 | hsa-mir-15a-5p | Up | ESR2 | 28 | RCOR1 |
| Up | ESR2 | 1 | hsa-mir-335-5p | Up | DLK1 | 28 | LMO2 |
| Up | PTGER3 | 1 | hsa-mir-148a-3p | Up | PTN | 27 | MYCN |
| Down | CDKN1A | 187 | hsa-mir-30b-3p | Down | CDKN1A | 57 | TP53 |
| Down | HSPA8 | 116 | hsa-mir-338-5p | Down | HSPA8 | 39 | KLF4 |
| Down | FASN | 116 | hsa-mir-103a-3p | Down | FASN | 39 | EGR1 |
| Down | E2F1 | 74 | hsa-mir-150-5p | Down | ZWINT | 37 | AHR |
| Down | CDT1 | 45 | hsa-mir-202-3p | Down | BCAR1 | 37 | TRIM28 |
| Down | MCM5 | 41 | hsa-mir-145-5p | Down | E2F1 | 36 | MITF |
| Down | GTF2I | 33 | hsa-mir-124-3p | Down | CDT1 | 34 | SIN3B |
| Down | ZWINT | 28 | hsa-mir-126-3p | Down | TUBB2B | 34 | SMAD3 |
| Down | TUBB2B | 24 | hsa-mir-101-3p | Down | GTF2I | 33 | AR |
| Down | BCAR1 | 21 | hsa-mir-1296-5p | Down | ATP13A2 | 28 | TCF7 |
| Down | GLIS2 | 20 | hsa-mir-31-3p | Down | MCM5 | 27 | KLF1 |
| Down | SFN | 19 | hsa-mir-200c-3p | Down | SFN | 21 | REST |
| Down | ATP13A2 | 13 | hsa-mir-4523 | Down | GLIS2 | 19 | SOX2 |
| Down | ACTA1 | 7 | hsa-mir-27a-5p | Down | ACTA1 | 13 | SRY |
| Down | FOXP3 | 1 | hsa-mir-372-3p | Down | FOXP3 | 10 | CCND1 |

### TF-hub gene regulatory network construction

To investigate the molecular mechanisms underlying the hub genes and TFs were searched by bioinformatics methods. The TFs of hub genes were predicted by NetworkAnalyst database. The TF-hub gene regulatory network is consisted of 441 nodes and 6738 edges, including 181 TFs and 260 hub genes (Fig. 6). As shown in Table 5, TFs of hub genes were displayed. JUN might be the target of 61 TFs (ex; GATA2), SOCS3 might be the target of 56 TFs (ex; RUNX1), TFAP2A might be the target of 55 TFs (ex; SUZ12), ATF3 might be the target of 48 TFs (ex; NUCKS1), CEBPB might be the target of 46 TFs (ex; FOXP2), CDKN1A might be the target of 57 TFs (ex; TP53), HSPA8 might be the target of 39 TFs (ex; KLF4), FASN might be the target of 39 TFs (ex; EGR1), ZWINT might be the target of 37 TFs (ex; AHR) and BCAR1 might be the target of 37 TFs (ex; TRIM28).

**Fig. 6.**
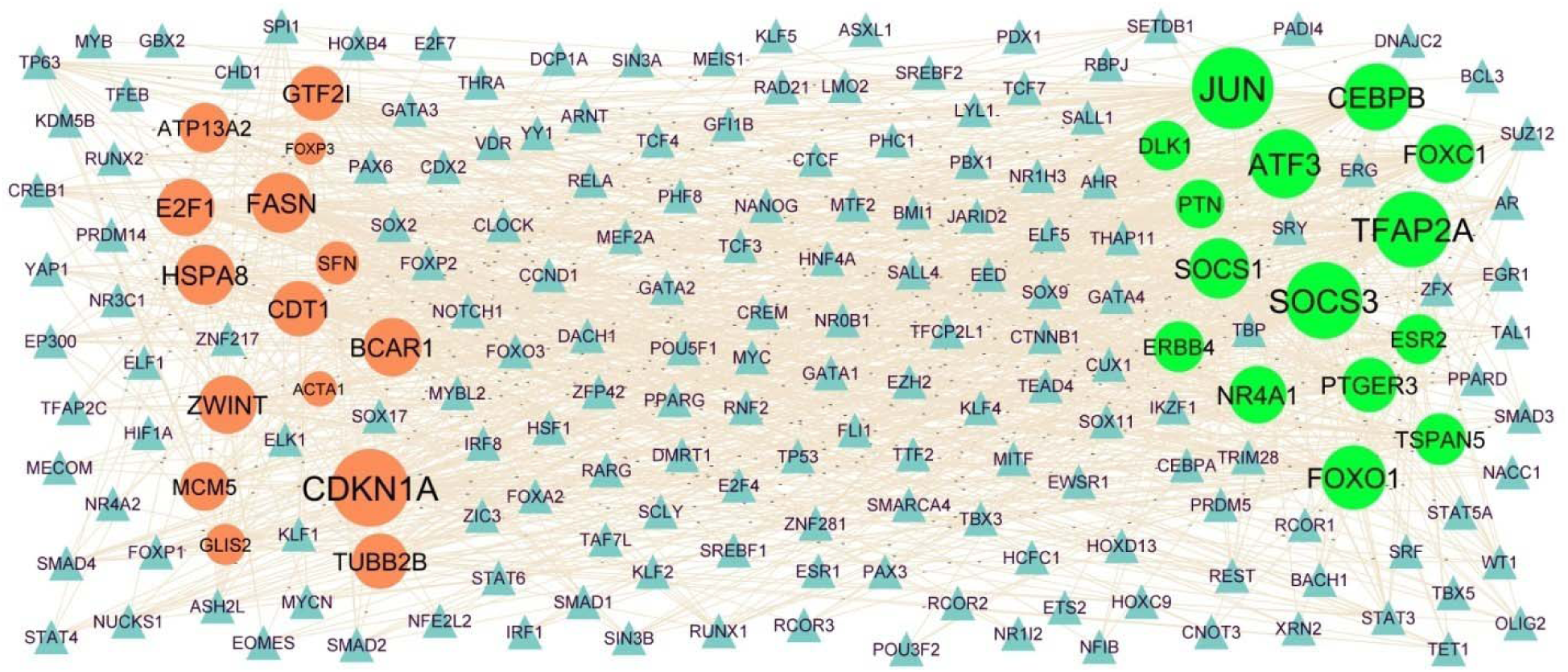
Target gene - TF regulatory network between target genes. The blue color triangle nodes represent the key TFs; up regulated genes are marked in green color; down regulated genes are marked in red color.

**Fig. 7.**
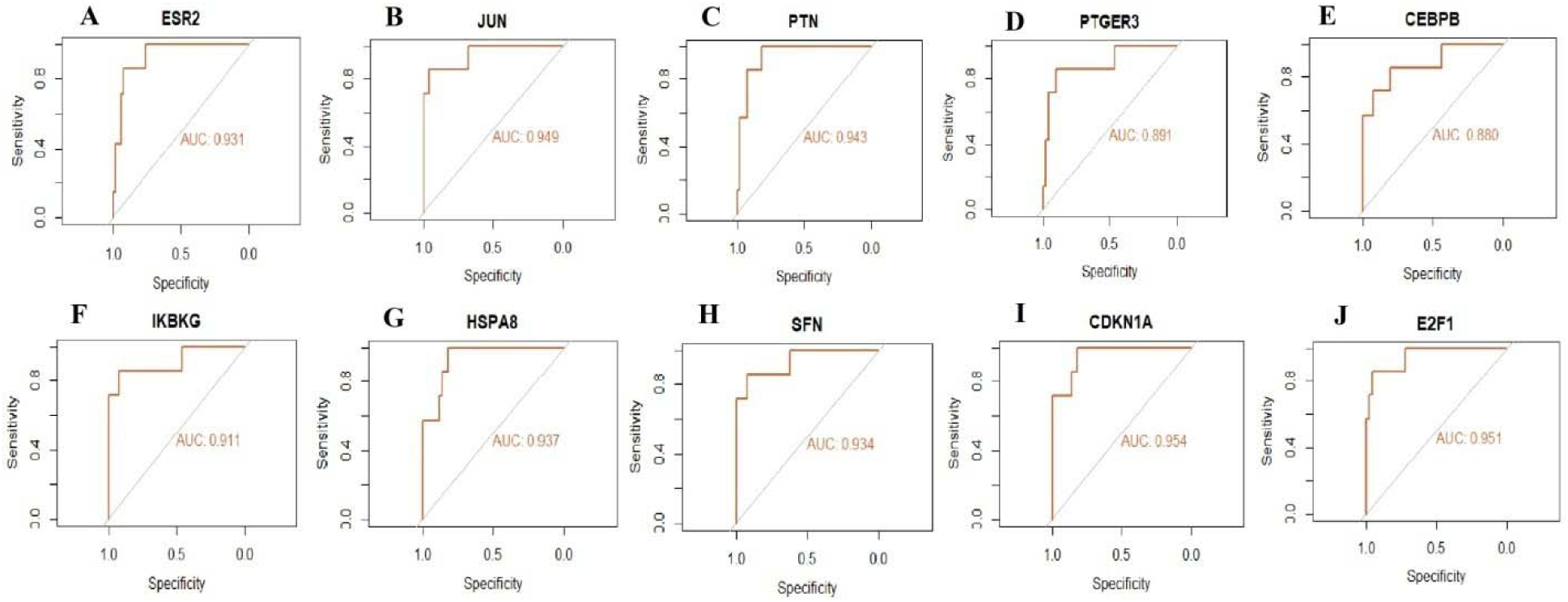
ROC curve analyses of hub genes. A) ESR2 B) JUN C) PTN D) PTGER3 E) CEBPB F) IKBKG G) SFN I) CDKN1A J) E2F1

### Receiver operating characteristic curve (ROC) analysis

ROC curve analysis using “pROC” in R statistical software packages was performed to calculate the capacity of ten hub genes to distinguish NAFLD samples from normal control samples. ESR2, JUN, PTN, PTGER3, CEBPB, IKBKG, HSPA8, SFN, CDKN1A and E2F1 all exhibited excellent diagnostic efficiency (AUC > 0.88) (Fig. 6).

## Discussion

The NAFLD is a metabolic disease in light of molecular characteristics, distinct morphologies, metabolic pathways, therapeutic response and clinical outcomes. Therefore, it is important to explore the mechanisms of NAFLD progression to limit its advancement. The NGS platforms for detection of gene expression have been progressing briskly in diseases advancement, which provides the basis of novel targets discovery for diagnosis and therapy for NAFLD.

In the current investigation, NGS dataset, GSE135251 was extracted to identify the DEGs between NAFLD samples and normal control samples. The results showed that there are 476 up regulated and 475 down regulated genes between the NAFLD samples and normal control samples by using bioinformatics analysis. Altered expression of FOS (Fos proto-oncogene, AP-1 transcription factor subunit) [44], IGFBP1 [45], IL6 [46], RGS1 [47], NR4A3 [48] and SLC22A12 [49] occurs in hypertension, but these genes might be novel targets for NAFLD. Pan et al. [50], McKee et al. [51], Simon et al. [52] and Shi et al. [53] proved that IGFBP1, AREG (amphiregulin), IL6 and AKR1B10 were involved in NAFLD progression. Ross et al. [54] and Lin et al. [55] further demonstrated that IGFBP1 and IL6 was involved in liver cirrhosis. IGFBP1 [56], AREG (amphiregulin) [57], IL6 [58], NR4A3 [59], EEF1A2 [60] and AKR1B10 [61] are implicated in the growth and metastasis of hepatocellular carcinoma. IGFBP1 [62], IL6 [63], RGS1 [64], NR4A3 [65], EEF1A2 [66] and MYH7 [67] acts in the progression of cardiovascular diseases, but these genes might be novel targets for NAFLD. IGFBP1 [68], IL6 [69], NR4A3 [70] and SLC22A12 [49] plays in the development of obesity, but these genes might be novel targets for NAFLD. Increasing evidence suggested that IGFBP1 [71] and IL6 [72] are involved in diabetes mellitus development and progression, but these genes might be novel targets for NAFLD. Recent studies found that AREG (amphiregulin) [73], IL6 [74] and RGS1 [75] plays an important role in the occurrence and development of cerebrovascular diseases, but these genes might be novel targets for NAFLD. IL6 [76] has been demonstrated to function in kidney disease, but this gene might be novel target for NAFLD.

Subsequently, GO annotation and REACTOME pathway enrichment analysis were performed to elucidate the significance of these identified altered expressed genes. Pathways and GO annotation include signal transduction [77], GPCR ligand binding [78], metabolism of lipids [79], fatty acid metabolism [80], signaling [81], chromatin [82], cell periphery [83] and membrane [84] were linked with progression of NAFLD. EGR1 [85], NR4A1 [86], DLK1 [87], SIK1 [88], CCN1 [89], SOCS2 [90], IRS2 [91], GADD45B [92], SOCS3 [93], MMP1 [94], CXCR4 [95], THBS1 [96], FST (follistatin) [97], KLF4 [98], CCL2 [99], OSM (oncostatin M) [100], MSTN (myostatin) [101], EPO (erythropoietin) [102], IL10 [103], JUN (Jun proto-oncogene, AP-1 transcription factor subunit) [104], FOXO1 [105], DKK1 [106], PTN (pleiotrophin) [107], ATF3 [108], SOCS1 [109], GPR183 [110], AKR1B10 [111], TREM2 [112], SCD (stearoyl-CoA desaturase) [113], GCK (glucokinase) [114], LPL (lipoprotein lipase) [115], ANGPTL8 [116], UGT1A1 [117], IL2RA [118], CDKN1A [119], GAS6 [120], ACE2 [121], NLRP6 [122], S1PR4 [123], FADS2 [124], FASN (fatty acid synthase) [125], TIMP3 [126], CHI3L1 [127], ADAMTSL2 [128], FNDC5 [129], MMP9 [130] and TMC4 [131] were involved in progression of NAFLD. EGR1 [132], NR4A2 [133], DUSP1 [134], DLK1 [135], SIK1 [136], CCN1 [137], CEBPD (CCAAT enhancer binding protein delta) [138], SOCS2 [139], IRS2 [91], GADD45B [140], CXCL2 [141], ZFAND5 [142], EGR3 [143], PTGS2 [144], KL (klotho) [145], SOCS3 [146], MMP1 [147], CXCR4 [148], THBS1 [149], SLC7A11 [150], FAM83A [151], EGR2 [152], ZFP36 [153], NR0B2 [154], CCN2 [155], FST (follistatin) [156], PCK1 [157], CTLA4 [158], KLF4 [159], PRDM1 [160], CCL2 [161], OSM (oncostatin M) [162], ERBB4 [163], MSTN (myostatin) [164], ADAMTS1 [165], PDK4 [166], INHBB (inhibin subunit beta B) [167], EPO (erythropoietin) [168], TSPAN5 [169], RASSF10 [170], GATA3 [171], IL10 [172], ESR2 [173], VIP (vasoactive intestinal peptide) [174], CEBPB (CCAAT enhancer binding protein beta) [175], FOXC1 [176], FOXO1 [177], DKK1 [178], RNF152 [179], SOX17 [180], PTN (pleiotrophin) [181], GAS1 [182], PMAIP1 [183], CCL3 [184], FGF2 [185], ATF3 [186], FZD7 [187], ANGPT2 [188], SOCS1 [189], MAP3K8 [190], LPAR6 [191], EYA4 [192], FOSL1 [193] MAFF (MAF bZIP transcription factor F) [194], KLF6 [195], BASP1 [196], CTCFL (CCCTC-binding factor like) [197], AKR1B10 [198], S1PR2 [199], COMP (cartilage oligomeric matrix protein) [200], IGF2BP1 [201], ACHE (acetylcholinesterase) [202], TREM2 [203], SCD (stearoyl-CoA desaturase) [204], FOXP3 [205], SCIN (scinderin) [206], EPHB2 [207], ANGPTL8 [208], HSPA8 [209], KCNJ11 [210], PFKL (phosphofructokinase, liver type) [211], EYA2 [212], GLIS2 [213], CDKN1A [214], ITGB3 [215], GAS6 [216], PIEZO1 [217], SYT7 [218], AGRN (agrin) [219], DAB2IP [220], EXO1 [221], WNK2 [222], ACE2 [223], SPP1 [224], ANXA6 [225], E2F1 [226], CLDN3 [227], ALDH3A1 [228], ROS1 [229], FASN (fatty acid synthase) [230], NKD2 [231], TIMP3 [232], CHI3L1 [233], RAB25 [234], SUSD2 [235], LAMB3 [236], LYPD1 [237], ADAMTSL5 [238], FAT1 [239], GPBAR1 [240], FNDC5 [241], HLA-G [242], TRAF7 [243], MMP9 [244], TMC4 [245], TNFRSF12A [246], CABYR (calcium binding tyrosine phosphorylation regulated) [247], BMP10 [248] and LOXL4 [249] contributes to the progression of hepatocellular carcinoma. Previous studies have demonstrated that EGR1 [250], SIK1 [251], PTGS2 [252], RGS2 [253], KL (klotho) [254], SOCS3 [255], MMP1 [256], CXCR4 [257], THBS1 [258], CHRM3 [259], KLF4 [260], CACNA1A [261], CCL2 [262], AVPR1A [263], PDK4 [264], EPO (erythropoietin) [265], IL10 [266], ESR2 [267], VIP (vasoactive intestinal peptide) [268], FOXC1 [269], FOXO1 [270], S100B [271], FGF7 [272], SOX17 [273], FGF2 [274], ATF3 [275], IMPA1 [276], HPGDS (hematopoietic prostaglandin D synthase) [277], F2RL3 [278], SCUBE1 [279], COMP (cartilage oligomeric matrix protein) [280], ADAMTS16 [281], ACHE (acetylcholinesterase) [282], SLC6A2 [283], TREM2 [284], GCK (glucokinase) [285], FOXP3 [286], CYP7A1 [287], CMA1 [288], SLC22A2 [289], LPL (lipoprotein lipase) [290], ANGPTL8 [291], HSPA8 [292], MB (myoglobin) [293], KCNJ11 [294], MUC6 [295], UGT1A1 [296], KCNJ5 [297], GDF2 [298], ARHGEF18 [299], ITGB3 [300], P2RY2 [301], PIEZO1 [302], PDGFA (platelet derived growth factor subunit A) [303], ACE2 [304], NLRP6 [305], ROS1 [306], FASN (fatty acid synthase) [307], TIMP3 [308], CHI3L1 [309], TREH (trehalase) [310], SLC6A19 [311], FNDC5 [312], HLA-G [313], MMP9 [314], SLC22A18 [315] and BMP10 [316] are an independent prognostic factors in hypertension, but these genes might be novel targets for NAFLD. EGR1 [317], NR4A1 [318], NR4A2 [319], DUSP1 [320], GADD45G [321], DLK1 [322], SOCS2 [323], GADD45B [324], CXCL2 [325], EGR3 [326], PTGS2 [327], RGS2 [328], KL (klotho) [329], SOCS3 [330], MMP1 [331], CXCR4 [332], THBS1 [333], SLC7A11 [334], EGR2 [335], ZFP36 [336], KCNE4 [337], CCN2 [338], FST (follistatin) [339], CLEC4E [340], OGN (osteoglycin) [341], CTLA4 [342], KLF4 [343], CYTL1 [344], IL1RL1 [345], HBEGF (heparin binding EGF like growth factor) [346], CCL2 [347], OSM (oncostatin M) [348], PRRX1 [349], ERBB4 [350], MSTN (myostatin) [351], ADAMTS1 [352], PDK4 [353], EPO (erythropoietin) [354], GATA3 [355], IL10 [356], ADTRP (androgen dependent TFPI regulating protein) [357], ESR2 [358], VIP (vasoactive intestinal peptide) [359], KCNIP1 [360], PLA2G2A [361], FOXC1 [362], FOXO1 [363], S100B [364], DKK1 [365], SOX17 [366], SLC6A4 [367], CCL3 [368], NPHP3 [369], ATF3 [370], ANGPT2 [371], SOCS1 [372], MYZAP (myocardial zonulaadherens protein) [373], EYA4 [374], F2RL3 [278], TFAP2A [375], MYH7 [376], SLC22A12 [377], SCUBE1 [378], S1PR2 [379], COMP (cartilage oligomeric matrix protein) [380], EDN2 [381], ACHE (acetylcholinesterase) [382], SLC6A2 [383], TREM2 [384], LMOD2 [385], GCK (glucokinase) [386], FOXP3 [387], CYP7A1 [388], LPL (lipoprotein lipase) [389], ANGPTL8 [390], HCN2 [391], HSPA8 [392], MB (myoglobin) [393], KCNJ11 [394], HRH2 [395], TNFSF13 [396], UGT1A1 [397], IL2RA [398], KCNJ5 [399], ITGB3 [400], P2RY2 [401], GAS6 [402], PIEZO1 [403], DAB2IP [404], ACE2 [405], KCNE1 [406], S1PR4 [407], E2F1 [408], ROS1 [409], FADS2 [410], TIMP3 [411], CHI3L1 [412], PHLDA3 [413], MMP17 [414], RASAL1 [415], SLC5A10 [416], SUSD2 [417], OLFM2 [418], LAMA3 [419], BCAR1 [420], TREH (trehalase) [421], FNDC5 [422], HLA-G [313], MMP9 [423] and MYL7 [424] are an important members in the progress of cardiovascular diseases, but these genes might be novel targets for NAFLD. Previous studies have shown that EGR1 [425], NR4A1 [426], DUSP1 [427], DLK1 [428], SIK1 [429], SOCS2 [430], IRS2 [431], SOCS3 [432], CXCR4 [433], THBS1 [434], GPAT3 [435], ZFP36 [436], DUSP2 [437], CCN2 [438], FST (follistatin) [439], PCK1 [440], KLF4 [441], IL1RL1 [442], CCL2 [443], OSM (oncostatin M) [444], MSTN (myostatin) [445], PDK4 [446], EPO (erythropoietin) [447], IL10 [448], VIP (vasoactive intestinal peptide) [449], CEBPB (CCAAT enhancer binding protein beta) [450], FOXO1 [451], S100B [452], DKK1 [453], SLC6A4 [454], CCL3 [455], FGF2 [456], ATF3 [457], SOCS1 [109], IL1RAP [458], MAP3K8 [459], SIM1 [460], SCUBE1 [461], CIDEA (cell death inducing DFFA like effector a) [462], ACHE (acetylcholinesterase) [463], TREM2 [464], SCD (stearoyl-CoA desaturase) [465], GCK (glucokinase) [466], CYP7A1 [467], LPL (lipoprotein lipase) [468], ANGPTL8 [469], KCNJ11 [470], UGT1A1 [471], KCNJ5 [472], GAS6 [473], ACE2 [474], MCHR1 [475], SPP1 [476], E2F1 [477], FADS2 [478], FASN (fatty acid synthase) [479], TIMP3 [480], CHI3L1 [481], MFAP5 [482], LAMB3 [483], SERPINA12 [484], GPBAR1 [485], TREH (trehalase) [486], ANGPTL7 [487], FNDC5 [488], HLA-G [489] and MMP9 [490] are involved in the progression of obesity, but these genes might be novel targets for NAFLD. Studies indicated that EGR1 [491], NR4A1 [492], DLK1 [493], SOCS2 [494], IRS2 [495], PTGS2 [496], RGS2 [497], SOCS3 [498], MMP1 [499], CXCR4 [500], FST (follistatin) [439], PCK1 [501], CHRM3 [502], CTLA4 [503], CCL2 [504], OSM (oncostatin M) [505], EPO (erythropoietin) [506], GATA3 [507], IL10 [508], ESR2 [509], VIP (vasoactive intestinal peptide) [510], KCNIP1 [511], PLA2G2A [361], FOXO1 [451], S100B [512], DKK1 [513], FGF7 [514], RGS16 [515], SLC6A4 [516], CCL3 [517], FGF2 [518], ATF3 [519], SOCS1 [372], COMP (cartilage oligomeric matrix protein) [520], IGF2BP1 [521], ACHE (acetylcholinesterase) [522], TREM2 [523], SCD (stearoyl-CoA desaturase) [524], GCK (glucokinase) [525], EIF2S3B [526], PDX1 [527], FOXP3 [528], CYP7A1 [529], LPL (lipoprotein lipase) [530], ANGPTL8 [531], HSPA8 [532], MB (myoglobin) [533], KCNJ11 [534], UGT1A9 [535], IL2RA [536], CDKN1A [537], SLC29A4 [538], GAS6 [539], PIEZO1 [540], ACE2 [541], E2F1 [542], ALDH3A1 [543], FADS2 [544], FASN (fatty acid synthase) [545], CHI3L1 [546], VTCN1 [547], SERPINA12 [548], GPBAR1 [549], BCAR1 [550], TREH (trehalase) [551], CFB (complement factor B) [552], ANGPTL7 [553], FNDC5 [554], HLA-G [313] and MMP9 [555] played important roles in promoting the diabetes mellitus, but these genes might be novel targets for NAFLD. Németh et al. [556], Westbrook et al. [557], Hu et al. [558], Hu et al. [559], Duitman et al. [560], Landau et al. [561], Jang et al. [562], Kaludjerovic et al. [563], Li et al. [564], Zhang et al. [565], Dwivedi et al. [566], Mehta et al. [567], Baek et al. [341], Grywalska et al. [568], Xu et al. [569], Yu et al. [570], Sakai et al. [571], Feng et al. [572], Elbjeirami et al. [573], Feng et al. [574], Li et al. [575], Bataille et al. [576], Schrimpf et al. [577], Mitra et al. [578], Sun et al. [579], Hanudel et al. [580], Peda et al. [581], Yamaguchi et al. [582], Wang and He [583], Neto et al. [584], Luna-Antonio et al. [585], Hoff et al. [586], Li et al. [587], Cheng and Lin, [588], Tsai et al. [589], Chen et al. [590], Niitsuma et al. [591], Atanasio et al. [592], Sanchez-Niño et al. [593], Shaw et al. [594], Ichida et al. [595], Mazumder et al. [596], Cao et al. [597], Niida et al. [598], Hishida et al. [599], Hirooka et al. [600], Sallinen et al. [289], Huang et al. [601], Ćwiklińska et al. [602], Zou et al. [603], Tsantoulas et al. [604], Oguri et al. [292], Lenglet et al. [605], Pietrzak-Nowacka et al. [606], Lu et al. [607], Zhong et al. [608], Iatrino et al. [609], Yanagita, [610], Yard et al. [611], Maksimowski et al. [612], Valiño-Rivas et al. [613], Guan et al. [614], Wang et al. [615], Hoste et al. [616], Mao, [617], Fabretti et al. [618], Xiao et al. [619], Atwood et al. [620], Lu et al. [621], Khidr [554], Hauer et al. [622], Laucyte-Cibulskiene et al. [623], Yang et al. [624] and Matejas et al. [625] showed that EGR1, NR4A1, SIK1, CCN1, CEBPD (CCAAT enhancer binding protein delta), SOCS2, RGS2, KL (klotho), SOCS3, MMP1, CXCR4, CCN2, FST (follistatin), OGN (osteoglycin), CTLA4, KLF4, PTGER3, HBEGF (heparin binding EGF like growth factor), CCL2, OSM (oncostatin M), ERBB4, TNFSF11, MSTN (myostatin), ADAMTS1, CALCR (calcitonin receptor), INHBB (inhibin subunit beta B), EPO (erythropoietin), IL10, CEBPB (CCAAT enhancer binding protein beta), FOXO1, DKK1, GAS1, NPHP3, FGF2, ATF3, ANGPT2, SOCS1, IL1RAP, C9ORF72, BASP1, AKR1B10, SLC22A12, ACHE (acetylcholinesterase), TREM2, SCD (stearoyl-CoA desaturase), GCK (glucokinase), FOXP3, SLC22A2, EPHB2, LPL (lipoprotein lipase), ANGPTL8, HCN2, HSPA8, MB (myoglobin), KCNJ11, GLIS2, TNFSF13, LSS (lanosterol synthase), GAS6, AGRN (agrin), ACE2, NLRP6, DPEP1, TIMP3, CHI3L1, RASAL1, FAT1, GPBAR1, TREH (trehalase), CFB (complement factor B), FNDC5, HLA-G, MMP9, C4A and LAMB2 were associated with kidney disease, but these genes might be novel targets for NAFLD. Xu et al. [626], Moles et al. [627], Ogata et al. [628], Okamoto et al. [629], Liu et al. [630], Nakamura et al. [631], Liu et al. [632], Zhu et al. [633], Kovalenko et al. [634], Yang et al. [635], Li et al. [636], Znoyko et al. [637], Gong et al. [638], Diamantis et al. [639], Guo et al. [640], Lee et al. [641], Saleh et al. [642], Heinrichs et al. [643], Kurniawan et al. [644], She et al. [645], Mafanda et al. [646], Norman et al. [200], García-Ayllón et al. [647], Zhao et al. [648], Butler et al. [649], Urbánek et al. [650], Smirne et al. [651], Wu et al. [652], Yang et al. [653], Nishimura et al. [654], Han et al. [655] and Barascuk et al. [656] revealed that IRS2, CXCL2, SOCS3, MMP1, CXCR4, SLC7A11, CCN2, KLF4, CCL2, OSM (oncostatin M), PRRX1, ADAMTS1, IL10, VIP (vasoactive intestinal peptide), S100B, CCL3, FGF2, MSMP (microseminoprotein, prostate associated), SOCS1, COMP (cartilage oligomeric matrix protein), ACHE (acetylcholinesterase), FOXP3, EPHB2, UGT1A1, GAS6, ACE2, E2F1, CHI3L1, PHLDA3 and MMP9 are associated with liver cirrhosis. Altered expression of KL (klotho) [657], SOCS3 [658], MMP1 [659], CXCR4 [660], RASD1 [661], SLC7A11 [662], OGN (osteoglycin) [341], KLF4 [663], IL1RL1 [664], CCL2 [665], EPO (erythropoietin) [666], IL10 [667], ESR2 [668], VIP (vasoactive intestinal peptide) [669], FOXC1 [670], FOXO1 [671], S100B [672], DKK1 [673], SOCS1 [674], C9ORF72 [675], CTCFL (CCCTC-binding factor like) [676], S1PR2 [677], ACHE (acetylcholinesterase) [678], TREM2 [679], EPHB2 [680], LPL (lipoprotein lipase) [681], P2RY2 [682], ACE2 [683], CHI3L1 [684], HLA-G [685] and MMP9 [686] are found in cerebrovascular diseases, but these genes might be novel targets for NAFLD.

By analyzing the PPI network and modules, a number of hub genes were identified that may provide new approaches for therapeutic studies of NAFLD. Thus, IKBKG (inhibitor of nuclear factor kappa B kinase regulatory subunit gamma), SFN (stratifin), FOSB (FosB proto-oncogene, AP-1 transcription factor subunit), JUNB (JunB proto-oncogene, AP-1 transcription factor subunit) and JUND (JunD proto-oncogene, AP-1 transcription factor subunit) can be a novel molecular markers of NAFLD and provide new insight to improve therapeutic strategies for NAFLD complications.

A miRNA-hub gene regulatory network and TF-hub gene regulatory network were constructed with hub genes, and then miRNAs and TFs were identified. CDT1 [687], ZWINT (ZW10 interacting kinetochore protein) [688], hsa-mir-3613-3p [689], GATA2 [690], RUNX1 [691], SUZ12 [692], NUCKS1 [693], FOXP2 [694], TP53 [695], AHR (aryl hydrocarbon receptor) [696] and TRIM28 [697] were revealed to be correlated with disease outcome in patients with hepatocellular carcinoma. Hsa-mir-199a-5p [698], hsa-mir-22-5p [699], hsa-mir-338-5p [700], GATA2 [701], RUNX1 [702], SUZ12 [703], TP53 [704] and AHR (aryl hydrocarbon receptor) [705] were revealed to be associated with cardiovascular diseases, but these molecular markers might be novel targets for NAFLD. Hsa-mir-199a-5p [698], GATA2 [706], FOXP2 [707] and AHR (aryl hydrocarbon receptor) [708] have been demonstrated to function in hypertension, but these genes might be novel targets for NAFLD. Previous studies have demonstrated that hsa-mir-22-5p [709], hsa-mir-150-5p [710], RUNX1 [711], TP53 [712], AHR (aryl hydrocarbon receptor) [713] and TRIM28 [714] are an independent prognostic factor in obesity and might be associated with obesity pathogenesis, but these genes might be novel targets for NAFLD. Wang et al. [715], Lu et al. [716], Shi et al. [717], Yu et al. [718], Zhang et al. [719] and Ren et al. [720] reported that altered expression of hsa-mir-3613-3p, hsa-mir-103a-3p, hsa-mir-150-5p, GATA2, RUNX1 and AHR (aryl hydrocarbon receptor) correlated in patients with kidney disease, but these genes might be novel targets for NAFLD. Previous reports have demonstrated that hsa-mir-150-5p [721] and RUNX1 [722] are associated with liver cirrhosis. Increasing evidence demonstrated that hsa-mir-150-5p [723] and TP53 [724] have a function in diabetes mellitus, but these genes might be novel targets for NAFLD. Bertran et al. [725], Yan et al. [726] and Zhu et al. [727] revealed that RUNX1, TP53 and AHR (aryl hydrocarbon receptor) may be the potential targets for NAFLD diagnosis and treatment. Thus, hsa-mir-33a-3p, hsa-mir-561-3p, hsa-mir-30b-3p and hsa-mir-202-3p can be novel molecular markers of NAFLD and provide new insight to improve therapeutic strategies for NAFLD complications.

In conclusion, the large-scale data analysis of the current investigation provides a comprehensive bioinformatics analysis of DEGs that might be associated in the progress of NAFLD. This investigation provides a set of important targets for future studiws into the molecular mechanisms and molecular markers involved in NAFLD. The exploration of potential key genes of NAFLD might provide some potential help in further identification of novel molecular markers for the susceptibility of NAFLD and useful treatment targets. The real function of key genes needs to be explored further to determine the clinical and biological mechanism of NAFLD.

## Acknowledgement

I thank Simon Cockell, Newcastle University, Newcastle, United Kingdom, very much, the author who deposited their profiling by high throughput sequencing dataset GSE135251, into the public GEO database.

## Conflict of interest

The authors declare that they have no conflict of interest.

## Ethical approval

This article does not contain any studies with human participants or animals performed by any of the authors.

## Informed consent

No informed consent because this study does not contain human or animals participants.

## Availability of data and materials

The datasets supporting the conclusions of this article are available in the GEO (Gene Expression Omnibus) (https://www.ncbi.nlm.nih.gov/geo/) repository. [(GSE135251) https://www.ncbi.nlm.nih.gov/geo/query/acc.cgi?acc=GSE135251)]

## Consent for publication

Not applicable.

## Competing interests

The authors declare that they have no competing interests.

## Author Contributions

B. V. - Writing original draft, and review and editing

C. V. - Software and investigation

## Notes

### Competing Interest Statement

The authors have declared no competing interest.

